# Primary nasal viral infection rewires the tissue-scale memory response

**DOI:** 10.1101/2023.05.11.539887

**Authors:** Samuel W. Kazer, Colette Matysiak Match, Erica M. Langan, Marie-Angèle Messou, Thomas J. LaSalle, Elise O’Leary, Jessica Marbourg, Katherine Naughton, Ulrich H. von Andrian, Jose Ordovas-Montanes

**Author notes:** These authors contributed equally to the work (Samuel W. Kazer, Colette Matysiak Match). These senior authors contributed equally to the work (Ulrich H. von Andrian, Jose Ordovas-Montanes). Correspondence to Jose Ordovas-Montanes.

## Abstract

The nasal mucosa is frequently the initial site of respiratory viral infection, replication, and transmission. Recent work has started to clarify the independent responses of epithelial, myeloid, and lymphoid cells to viral infection in the nasal mucosa, but their spatiotemporal coordination and relative contributions remain unclear. Furthermore, understanding whether and how primary infection shapes tissue-scale memory responses to secondary challenge is critical for the rational design of nasal-targeting therapeutics and vaccines. Here, we generated a single-cell RNA-sequencing (scRNA-seq) atlas of the murine nasal mucosa sampling three distinct regions before and during primary and secondary influenza infection. Primary infection was largely restricted to respiratory mucosa and induced stepwise changes in cell type, subset, and state composition over time. Type I Interferon (IFN)-responsive neutrophils appeared 2 days post infection (dpi) and preceded transient IFN-responsive/cycling epithelial cell responses 5 dpi, which coincided with broader antiviral monocyte and NK cell accumulation. By 8 dpi, monocyte-derived macrophages (MDMs) expressing *Cxcl9* and *Cxcl16* arose alongside effector cytotoxic CD8 and *Ifng*-expressing CD4 T cells. Following viral clearance (14 dpi), rare, previously undescribed **K**rt13+ **n**asal **i**mmune-**i**nteracting **f**loor **e**pithelial (KNIIFE) cells expressing multiple genes with immune communication potential increased concurrently with tissue-resident memory T (TRM)-like cells and early IgG+/IgA+ plasmablasts. Proportionality analysis coupled with cell-cell communication inference, alongside validation by in situ microscopy, underscored the CXCL16–CXCR6 signaling axis between MDMs and effector CD8 T cells 8dpi and KNIIFE cells and TRM cells 14 dpi. Secondary influenza challenge with a homologous or heterologous strain administered 60 dpi induced an accelerated and coordinated myeloid and lymphoid response without epithelial proliferation, illustrating how tissue-scale memory to natural infection engages both myeloid and lymphoid cells to reduce epithelial regenerative burden. Together, this atlas serves as a reference for viral infection in the upper respiratory tract and highlights the efficacy of local coordinated memory responses upon rechallenge.

## INTRODUCTION

As the primary passage to the lower airway, the nasal mucosa balances the complex roles of olfaction, filtration and conditioning of inhaled air, and host defense. To accomplish these diverse functions, the nose contains distinct anatomical structures, harbors a varied yet organized cellular composition, and secretes a multitude of proteins with varied roles (Harkema et al., 2006). In the face of pathogens, the nasal mucosa is thought to mount a variety of incompletely understood defense mechanisms to protect against infection and limit spread to the lower respiratory tract (Bosch et al., 2013). Nevertheless, many respiratory pathogens manage to infect or colonize the upper airways and disseminate into the lungs, causing millions of cases of severe disease, hospitalizations, and deaths annually (Clark, 2020; Roth et al., 2018; Shinya et al., 2006).

There is a growing appreciation for how the inflammatory state of nasal tissue affects respiratory viral infection outcomes. The COVID-19 pandemic has helped accelerate research to understand the roles of interferons (IFNs) in nasal protection and disease trajectory, with studies highlighting the importance of sample timing and location, viral burden, and strain (Bastard et al., 2022; Kim and Shin, 2021; Park and Iwasaki, 2020; Sposito et al., 2021). Single-cell analysis of the human nasopharynx during SARS-CoV-2 infection showed muted IFN-responses in epithelial cells from severe disease relative to mild cases (Ziegler et al., 2021). Expression of specific IFN stimulated genes (ISGs) like *OAS1* associate with protection from severe COVID-19 and may even drive viral mutations to overcome host protection (Wickenhagen et al., 2021). More generally, evidence of a recent prior infection in children receiving a live-attenuated influenza vaccine was associated with enhanced ISG signaling and lower viral shedding (Costa-Martins et al., 2021). Collectively, this suggests that the present nasal state, cellular composition, and antiviral signaling capacity, as informed by the cumulative history of environmental exposures, may drive disease outcomes (Bastard et al., 2020; Habibi et al., 2020; Ordovas-Montanes et al., 2020; Weisberg et al., 2021; Zhang et al., 2020).

Following viral infection or intranasal (i.n.) vaccination, immune memory in the nasal mucosa can provide long-term protection both systemically and at the mucosal barrier, reducing pathology and infection burden in the lower airways and elsewhere (Johnson et al., 1986; Johnson Jr. et al., 1985; Rutigliano et al., 2010). Local protection is afforded by both cellular and humoral immune mechanisms. For example, CD8+ tissue-resident memory (TRM) cells that form following upper respiratory tract influenza A virus (IAV) infection correlate with enhanced protection against heterologous IAV strain rechallenge (Pizzolla et al., 2017). Protective mucosal IgA producing plasma cells, and antibodies capable of neutralizing virus, can be generated in the nasal mucosa following IAV, vesicular stomatitis virus, respiratory syncytial virus, and SARS-CoV-2 infections (Johnson Jr. et al., 1985; Liew et al., 2023; Sterlin et al., 2021; Wellford et al., 2022; Weltzin et al., 1996). Even so, respiratory infections like IAV remain epidemic and kill up to 500,000 people each year (Iuliano et al., 2018).

To develop protective, durable, and efficacious vaccines for respiratory viruses, we must reach a deeper understanding of the establishment, timing, and cooperation of tissue-scale memory following natural infection (Morens et al., 2023). Immune responses to pathogens often occur in stepwise fashion: recognition of pathogen-associated molecular patterns by immune and/or epithelial cells leads to cytokine production that broadly activates innate immune cells that in turn recruit pathogen-specific effector lymphocytes, some of which will develop into circulating and tissue-resident memory cells (Iwasaki and Medzhitov, 2015). Following prior infection or vaccination, however, these local circuits can be re-ordered and even inverted in the barrier tissues that re-encounter infection (Kadoki et al., 2017; Kaufmann et al., 2018; Ols et al., 2020; Ordovas-Montanes et al., 2020; Schenkel et al., 2014). During viral rechallenge, TRM cells exhibit antiviral effector functions and can act as sentinels that send antigen-specific inflammatory “alarms” to local immune cells to activate multicellular anti-microbial responses at the site of infection (Ariotti et al., 2014; McMaster et al., 2015; Schenkel et al., 2014; Steinbach et al., 2016). In mucosal vaccination, IFNγ produced by antigen-specific T cells is sufficient to induce increased inflammatory cytokine production by both distal (Bosch-Camós et al., 2022; Stary et al., 2015) and local (Yao et al., 2018) antigen presenting cells, suggesting that recruited and/or tissue-resident cells can contribute to rapid memory responses. Similarly, antibodies can directly neutralize virus and also orchestrate a variety of antiviral effector functions through antibody Fc-receptor mediated binding by NK cells, macrophages, and neutrophils (Boudreau and Alter, 2019). However, most of these studies to date have focused on the role of individual cell types or limited interactions during a memory response.

Here, we present a tissue-scale single-cell RNA-sequencing (scRNA-seq) atlas of the murine nasal mucosa before and during primary IAV infection and secondary rechallenge. By sampling multiple regions, timepoints, and cell lineages, we develop a compositional landscape of the tissue and reveal how the diversity of cell subsets and states dynamically changes in response to infection and during a memory response. Primary IAV infection induced reproducible stepwise shifts in cell composition starting with increased IFN-responsive neutrophil subsets followed by broader antiviral/IFN-stimulated responses in epithelial, myeloid, and lymphoid immune cells. Next, monocyte-derived macrophages (MDMs) accumulated along with effector CD8 and CD4 T cells. Following viral resolution, early TRM cells and plasmablasts are established alongside increased frequencies of rare **K**rt13+ **n**asal **i**mmune-interacting **f**loor **e**pithelial (KNIIFE) cells expressing genes for several ligands and receptors known to modulate immune cell activity. Learning cell cluster identity in samples generated in memory and during either homologous or heterologous IAV rechallenge, we demonstrate the applicability of our atlas to inform newly generated data and show that the nasal memory response to IAV is accelerated and coordinated compared to primary infection. Collectively, our spatial and temporal datasets enumerate and characterize the diversity of cell types, states, and subsets in the murine nasal mucosa and highlight those recruited and local cell subsets that exhibit memory and respond to viral infection.

## RESULTS

### Nasal mucosa infection with influenza virus and tissue processing

We administered 10^4^ plaque forming units (pfu) of IAV H1N1 strain PR8 i.n. to awake, naïve mice in a small volume (5 µl/nostril) to restrict infection to the upper respiratory tract (Klinkhammer et al., 2018; Pizzolla et al., 2017), and collected and processed nasal mucosa tissue (n=3 biological samples/timepoint) by scRNA-seq at 0, 2, 5, 8, and 14 days post infection (dpi; “primary”) (**Figure 1A**).

**Figure 1:**
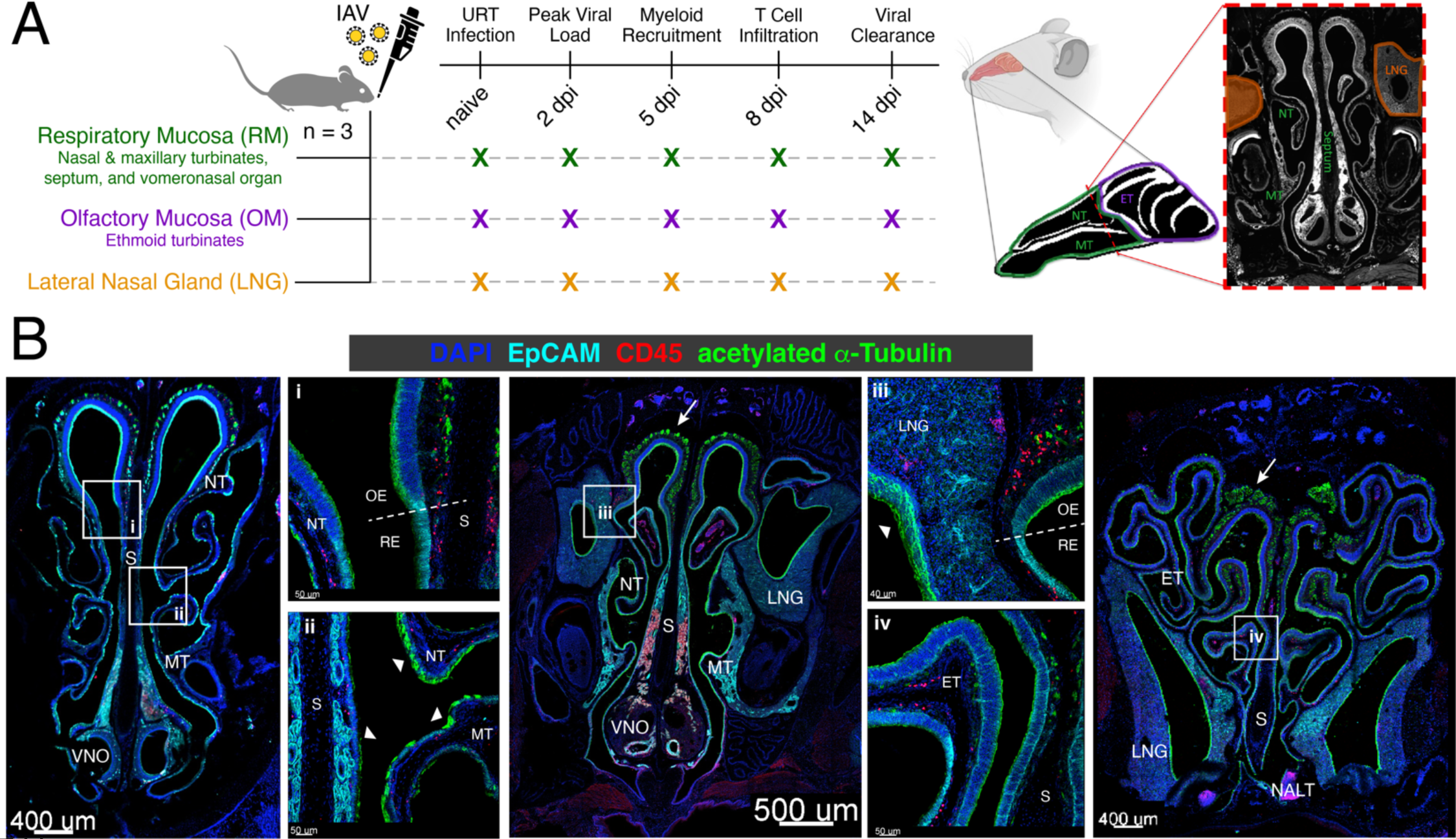
Experimental design and the structure of the murine nasal mucosa. (A) Schema depicting the sampling scheme and three tissue regions collected and processed for scRNA-seq: respiratory mucosa (RM), olfactory mucosa (OM), and lateral nasal gland (LNG). To infect, 10^4^ pfu Influenza A Virus (IAV) PR8 was administered intranasally (5µl/nostril). dpi = days post infection (B) Representative immunofluorescence images of coronal slices of the nasal mucosa from a naïve mouse moving from anterior (left) to dorsal (right) staining for epithelial cells (EpCAM, teal), immune cells (CD45, red), and ciliated cells/neurons (α-acetylated tubulin, green). Distinct regions of the mucosa are labeled. Labeled white boxes outline higher resolution images below. White arrows point to olfactory sensory nerve bundles; white arrowheads point to cilia. The olfactory epithelium (OE) and respiratory epithelium (RE) both reside within the collected RM tissue and are differentiated by morphology and the presence of olfactory sensory neurons. NT = nasoturbinate; S = septum; MT = maxillary turbinate; VNO = vomeronasal organ; LNG = lateral nasal gland; ET = ethmoid turbinate; NALT = nasal-associated lymphoid tissue.

Anatomically, the murine nasal mucosa can be divided into several distinct morphological, histological, and functional tissue regions (Harkema et al., 2006). We mapped the cellular and structural diversity in the naïve nasal mucosa by immunofluorescence imaging, observing broad heterogeneity in epithelial, immune, and neural distribution throughout the tissue (**Figure 1B**). Thus, to capture region-specific changes in cell composition and response following IAV infection, we micro-dissected the tissue and separated into three different regions: (1) respiratory mucosa (RM), inclusive of the nasal and maxillary turbinates, septum, and vomeronasal organ; (2) olfactory mucosa (OM), inclusive of the ethmoid turbinates; and (3) the lateral nasal gland (LNG), which sits underneath the RM and OM in the maxillary sinus (see **Methods**).

### Single-cell spatiotemporal atlas of primary influenza infection in the nasal mucosa

Across all primary infection timepoints (0-14 dpi), regions, and replicates (n=45), we collected 156,572 single-cell transcriptomes after filtering low-quality cell barcodes and hash-annotated cell multiplets (**Methods**). Top-level clustering on the entire primary infection dataset captured 42 clusters belonging to neural, epithelial, immune, and stromal (endothelial, fibroblast, and others) cell lineages demarcated by known lineage-restricted genes (**Figures 2A and S1A**). Neurons and epithelial cells comprised over half of the dataset, with immune and stromal cells comprising the rest (**Figure S1B**). Major cell types were found to be distributed differently across nasal regions, with enrichment of neurons, granulocytes, B cells, and hematopoietic stem cells (HSCs) in the OM and more epithelial cells, fibroblasts, myeloid cells, and T & NK cells in the RM (**Figures 2B and S1C**). As highly structured tissues, the turbinates in the nasal mucosa, and especially the ethmoid turbinate (OM), consist of substantial pieces of bone with sizeable bone marrow. Thus, the relative enrichment of specific immune cell types including HSCs and other immune progenitors in the OM likely reside in this bone marrow and may directly enter the mucosa and engage in pathogen defense (see **Figure 1B**). We note that succinctly depicting relative proportions of the data across multiple replicates is complex; stacked bar charts like **Figure 2B** show relative proportions within a cell type grouping, but do not reflect proportions within the cells captured at each time point/region.

**Figure 2:**
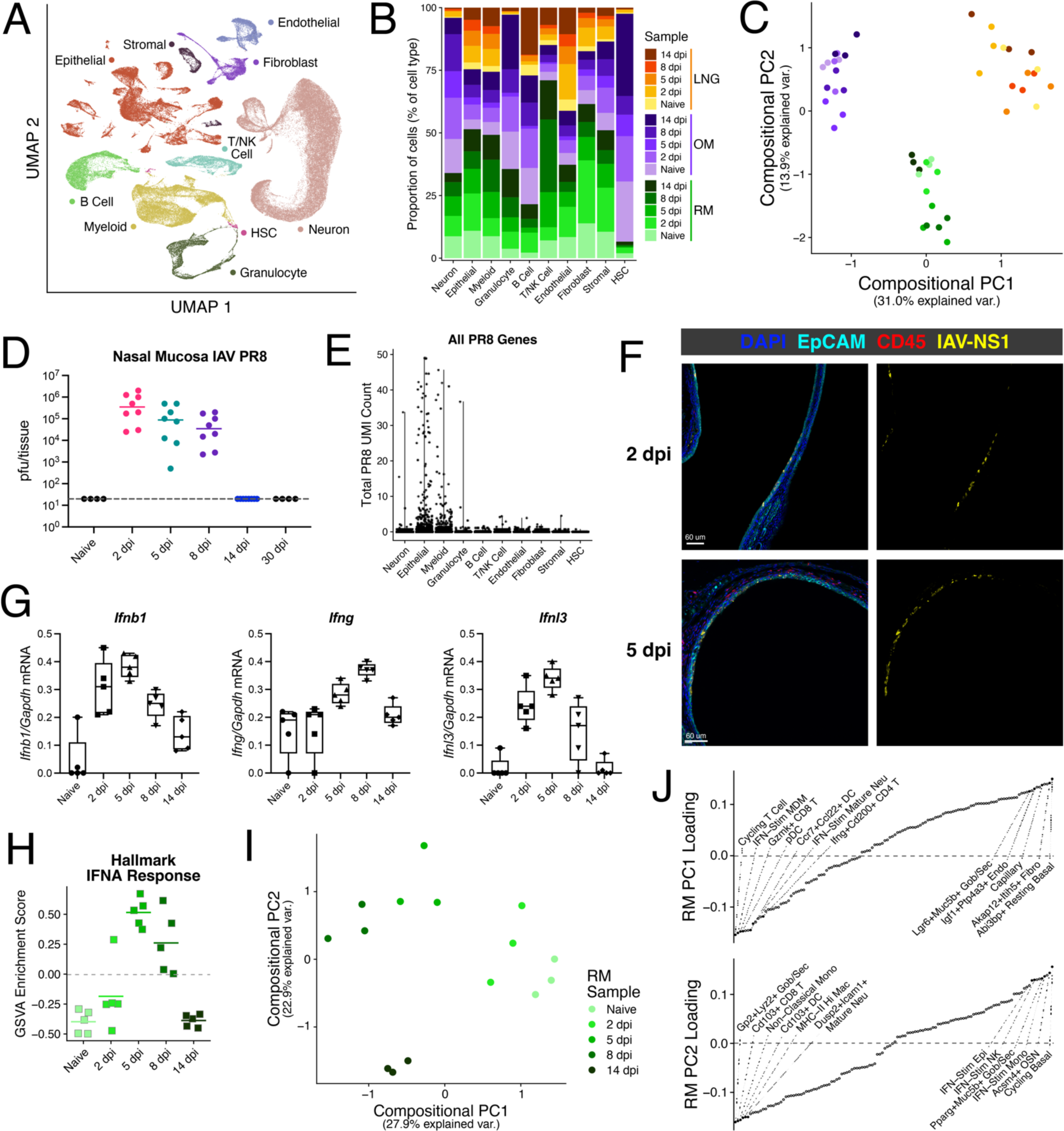
Single-cell atlas of the nasal mucosa during primary IAV infection. (A) UMAP embedding of 156,572 nasal cells across three tissue regions and five time points (n=3 per region/timepoint) colored by cell type. HSC = hematopoietic stem cell. (B) Stacked bar chart depicting the relative proportions of cells annotated for each cell type by region and time point. (C) Compositional principal component analysis (PCA) of all acute infection sample replicates. Each point represents a sample replicate and distance reflects variation in cell cluster abundance (scaled cell counts). Dots are colored by region and time point as in **B**. (D) Infectious IAV PR8 quantification in plaque forming units (pfu) of the entire nasal mucosa. (E) Summative scTransform-corrected UMI counts per cell across all IAV PR8 genes split by cell type. (F) Representative images of IAV infection in RM taken from mice 2 dpi (top) and 5 dpi (bottom). Staining for EpCAM (teal), CD45 (red), and IAV-NS1 (yellow). Images on the right depict only the signal in the IAV-NS1 channel. (G) qPCR of *Ifnb1* (left), *Ifng* (center), and *Ifnl3* from RNA extracted from RM tissue. C_q_ ratios are normalized by *Gapdh* C_q_. n = 5 per time point. (H) Gene Set Variation Analysis (GSVA) Enrichment score for Hallmark Response to Interferon-Alpha on total RM tissue lysate bulk RNA-seq data (n = 5/time point). (I) Compositional PCA of all cell clusters from only RM samples. (J) Cell cluster abundance loadings for PC1 (left) and PC2 (right) from (G). Cell cluster names for several of the most negative and most positive weights for each PC are depicted.

Sample replicates were called by demultiplexing oligo-hashtag count tables (Li et al., 2020) and did not exhibit strong batch effects (**Figure S1D**). While clustering and differential expression were performed on all singlets regardless of successful hash assignment, we counted only those cells with annotated sample replicates for downstream compositional analyses (see **Figure S1E** for assignment breakdown by cell type).

To delineate the diversity of cell subsets and states present in the nasal mucosa, we split the dataset by cell type and conducted new clustering analyses, yielding a total of 127 clusters across the dataset (**Figure S1F and Supplementary Table 1**). By counting the number of cells assigned to each cluster in each sample replicate and scaling across samples by cell capture, we calculated cell cluster abundances to interrogate the relationship between samples in cell compositional space (see **Methods and Supplementary Table 2**). Performing principal component analysis (PCA) on samples over center log-ratio (clr) transformed cell cluster abundances, we saw strong separation by region (**Figure 2C**), reinforcing that the nasal mucosa contains distinct regions with specific functions. Examination of the PCA loadings revealed that LNG is defined by higher abundances of odorant binding protein (OBP)-expressing cells, serous cells, and capillary endothelial cells (**Figure S1G**). OM has relatively more Schwann cells, HSCs, and glandular, while RM is enriched for vomeronasal sensory neurons, chondrocytes, and infection responsive epithelial cells. Collectively, this atlas represents a high-resolution, comprehensive view of the mouse nasal mucosa enabling characterization of the dynamic differences in cellular composition within and between nasal regions during infection.

### Influenza infection is largely restricted to the RM and induces reproducible changes in cellular composition

While viral, immune cell, and epithelial cell dynamics following IAV infection of the lung have been partially mapped (Bouvier and Lowen, 2010; Boyd et al., 2020; Manicassamy et al., 2010; Matsuoka et al., 2009; Steuerman et al., 2018), responses in the nasal mucosa are less studied. Viral titers of entire nasal mucosa showed robust infection 2 dpi that waned through 8 dpi and was completely cleared by 14 dpi (**Figure 2D**), while data from lungs showed sporadic spread of virus from the nasal mucosa only occurring between 5 and 8 dpi (**Figure S2A**). Aligning to a joint IAV and mouse genome, we also captured viral transcripts by scRNA-seq and thus could identify which cells may have been infected or contained virus. Individual genes like NP and HA were detected most strongly in epithelial and myeloid cells (**Figure S2B**). We calculated a summative IAV unique molecular identifier (UMI) count for every single cell to assess “positivity” for IAV (**Figure 2E**). Looking across time points and regions in epithelial and myeloid cells, we captured low, but reproducible numbers of IAV+ cells 2, 5, and 8 dpi in the RM aligning with the detectible plaque assays at these time points, but no positive cells in the OM, and only at 5 dpi in the LNG (**Figures S2C-D**). Bulk RNAseq of whole RM tissue lysate better matched the plaque forming assay results with higher IAV read counts than single-cell, suggesting non-cellular viral RNA and/or potential loss of IAV+ cells during processing for scRNA-seq **(Figure S2E**).

Staining for IAV NS1 at 2 and 5 dpi confirmed that infection was largely restricted to epithelial cells in the RM (**Figures 2F** and **S2F-G**) and is consistent with expression of binding receptors marked by α2,3-linked sialic acid in mucous producing cells (Ibricevic et al., 2006). We performed qPCR of total RM to validate an early response to infection and found robust upregulation of type I and III IFNs 2 dpi, with even higher expression 5 dpi despite relatively lower viral titers (**Figure 2G**). As expected, *Ifng* expression exhibited delayed kinetics peaking 8 dpi. To understand the dynamics of the global IFN-induced response during infection, we calculated enrichment scores for response to IFNα and IFNγ from bulk RNAseq data (**Figures 2H** and **S2H**). Despite elevated levels of *Ifnb1* at 2 dpi, IFN-response signaling was not enriched until the samples measured at 5 dpi; however, we cannot exclude that IFN-responses could start to occur between 2 and 5 dpi given sampling limitations.

To understand how infection remodels each nasal region, we applied PCA to the sample replicates within each region. In OM, 5 and 14 dpi samples separated from each other and the other time points (**Figure S2I**). In LNG, only 14 dpi samples separated from the rest (**Figure S2J**). Comparatively, PCA of RM samples showed clear separation between all timepoints in chronological order across the first two PCs (**Figure 2I**) suggesting dynamic and linked responses occur over the course of infection in this region. PCA loadings from RM highlight a shift in composition from resting basal cells, fibroblasts, and ciliated cells in naïve mice and 2 dpi to diverse activated myeloid and lymphoid clusters 5 and 8 dpi that gave way to specific goblet/secretory cells, TRM-like cells, and mature myeloid cells following viral resolution 14 dpi (**Figure 2J**). Even though virus had been cleared by 14 dpi, we note that all three nasal regions reached compositions distinct from naïve mice at this timepoint.

### Regional epithelial diversity in the nasal mucosa dynamically changes during IAV infection

#### Cellular diversity of the nasal epithelium

Having acquired high-level knowledge of how IAV infection broadly impacts the nasal mucosa, we next sought to understand the variety of epithelial cells present across the tissue and how they respond during infection as the main target of infection. Subclustering on all epithelial cells yielded 28 clusters encompassing diverse differentiation states and functions (**Figures 3A, S3A, and Supplementary Table 1**). We categorized these clusters into broader subsets including basal (*Krt5, Krt14*), ciliated (*Foxj1, Dnah5*), serous (*Ltf*, *Ccl9*), glandular (*Bpifb9b, Odam*), goblet/secretory (*Reg3g, Selenom, Scgb1c1,* and mucin-encoding genes), ionocyte (*Cftr, Coch*), tuft (*Trpm5, Il25*), and sustentacular cells (*Sec14l3, Cyp2g1*) (**Figure 3B**). In addition to known subsets, we also identified unique clusters of epithelial cells potentially specific to the nasal mucosa and present in naïve mice and throughout primary infection that separated distinctly in UMAP space: *Scgb-b27*+*Cck*+, *Klk1*+*Fxyd2*+, *Meg3*+*MHC-II*+, and *Krt13*+*Il1a*+ cells. Comparison to human nasal biopsy and swab datasets (Deprez et al., 2020; Ziegler et al., 2021), readily annotated known epithelial subsets, and suggested that the *Krt13*+*Il1a*+ cluster was squamous-like (**Figure S3B**). The other unique clusters did not map reliably to human subsets, potentially due to limitations in sampling human tissue. Epithelial clusters were differentially distributed across regions (**Figure 3C**): recently described nasal tuft cells, ionocytes, and *Dclk1*+ cells (Ualiyeva et al., 2024), as well as olfactory sensory neuron supportive sustentacular cells (Brann et al., 2020), were enriched in OM, whereas serous and glandular cells were specific to LNG.

**Figure 3:**
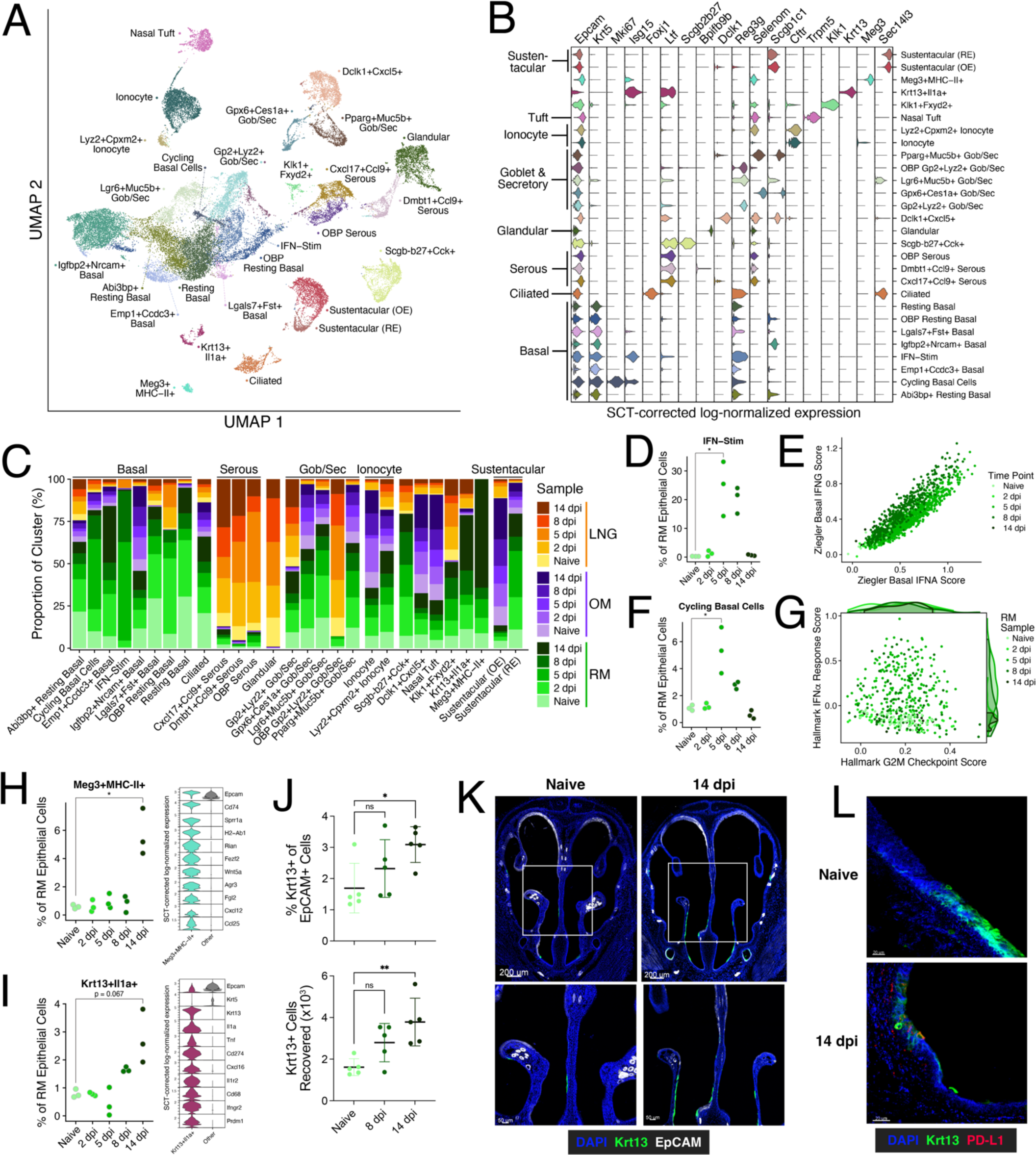
Epithelial cell subclustering reveals diverse subsets including transient IFN-responsive/cycling cells and rare cells with immune-interaction potential arising 14 dpi. (A) UMAP embedding of 38,367 epithelial cells across 27 clusters. (B) Violin plot depicting epithelial lineage and subset marker expression levels (scTransform-corrected log-normalized UMI counts) across all discovered clusters (see **Supplementary Table 1**). (C) Stacked bar chart depicting the relative proportions of cells annotated for each cluster by region and time point. (D) Relative frequencies of cells clustered as IFN-Stim as a proportion of all epithelial cells per replicate RM sample. Only cells with assigned hash calls are included. Welch’s t test, *p < 0.05. (E) IFN-Stim epithelial cell scores for signatures derived from airway basal cell cultures stimulated with IFNα or IFNγ (Ziegler et al., 2020). (F) Same as in (D) for the cycling basal cell cluster. (G) Scatter plot of gene module scores for the Hallmark IFNα Response and Hallmark G2M Checkpoint gene lists (MsigDB v7.5.1) in cycling basal cells. Density plots represent the scatter plot data. (H & I) Relative frequency plots of *Meg3*+MHC-II+ (G) and *Krt13*+*Il1a*+ (KNIIFE cells) (H) clusters (left) as a proportion of all epithelial cells per replicate RM sample. Violin plots of select cluster specific/enriched genes, except for *Epcam* and *Krt5* (FDR corrected p-values ≤ 10^–242^ by 1-vs-rest Wilcoxon Rank Sum Test) (right). (J) Mice were infected with 10^4^ PFU IAV PR8 and RM tissue was collected and stained intracellularly for Krt13+ epithelial cells (n = 5/timepoint). Kruskal-Wallis test, *p < 0.05, **p < 0.01. (K) Representative immunofluorescence images of the very anterior nasal mucosa in naïve mice (left) and 14 dpi (right) staining for Krt13 (green) and EpCAM (white). (L) Representative images within the region shown in (I) in naïve mice (top) and 14 dpi (bottom) staining for Krt13 (green) and PD-L1 (red). Welch’s t test, *p < 0.05.

#### Cycling and IFN-responsive epithelial cells arise in the RM during infection

To understand the impact of IAV infection on the nasal epithelial compartment, we first determined which clusters harbored viral reads. Looking at the distribution of IAV UMIs across all epithelial clusters, we found that IAV+ cells were most prevalent in the IFN-stimulated cluster followed by cycling basal and ciliated cells (**Figure S3C**). The IFN-stimulated cluster, largely restricted to the RM, exhibited high levels of *Krt5*, a basal cell marker, but also *Cxcl17*, which in our dataset is expressed at steady state in serous cells in the LNG and some goblet/secretory subsets. Comparing cells within the IFN-stimulated cluster by presence of IAV transcripts, we found relatively higher levels of ISGs but lower expression of transcription factors *Atf3*, *Egr1*, and *Junb* in IAV+ cells (**Figure S3D**). While infection was well-established at 2 dpi, IFN-stimulated epithelial cells only arose at 5 dpi and made up ∼20% of all RM epithelial cells 5 and 8 dpi (**Figure 3D**). The substantial detection of IAV transcripts by bulk RNA-seq (**Figure S2E**) but lack of an IFN-response signature in total RM tissue at 2 dpi (**Figure 2H**) suggests that besides selective loss of IAV+ cells during processing, host silencing mechanisms by IAV may contribute to the measured muted IFN-response as well (Kochs et al., 2007). To understand the relative contribution of type I/III and type II IFNs to the IFN-stimulated epithelial cluster, we scored these cells with gene lists derived from nasal basal cells stimulated with IFNα or IFNγ in vitro (Ziegler et al., 2020). While IFNα and IFNγ induce partially-overlapping ISGs, cells at 5 dpi scored higher for the IFNα signature, cells at 8 dpi scored higher for the IFNγ signature (**Figure 3E**). Closer exploration of genes with stronger induction following IFNα vs IFNγ stimulation confirmed the sequential response timing within IFN-stimulated epithelial cells (**Figure S3E**).

Cycling basal cells demonstrated a similar transient increase in abundance, peaking 5 dpi (**Figure 3F**). Interestingly, differential expression across timepoints revealed many ISGs were upregulated in these cells 5 dpi (**Figure S3F**). Given recent work demonstrating that the epithelial IFN-response can impede proliferation and tissue repair in the lower airway (Broggi et al., 2020; Major et al., 2020), we leveraged our single-cell resolution to assess if individual nasal basal cells co-express pathways for cell cycle and IFN-response. Gene set analysis confirmed significant enrichment for both cell cycle and IFN-response pathways in cycling basal cells. Additionally, a largely mutually exclusive apoptosis pathway was also significant (**Figures S3G**). Pathway module scoring showed that while the IFNα response score changed over time, the G2M checkpoint score was equally distributed across time points and independent of IFNα response (**Figure 3G**). Thus, unlike their lower respiratory tract counterparts, nasal cycling basal cells proliferate during primary infection and may concurrently support ISG expression alongside non-proliferative IFN-stimulated basal cells.

#### Rare unique epithelial cell subsets with lymphoid and myeloid communication potential accumulate following viral clearance

Looking at the compositional PCA, RM samples from 14 dpi separated from other timepoints (**Figure 2G**). Among epithelial cells, two clusters, *Emp1*+*Ccdc3*+ basal cells and *Gp2*+*Lyz2*+ goblet/secretory cells, accumulated to 9-12% of all epithelial cells following viral clearance by 14 dpi (**Figure S3H**). In addition to *Gp2* and *Lyz2*, the goblet/secretory subset was also enriched for *Il33*, *Muc1*, *Isg20*, and *Cd14* suggesting a potential shift in the epithelium toward an antibacterial state. Additionally, two rare and transcriptionally distinct subsets of epithelial cells (*Meg3*+MHC-II+ and *Krt13*+*Il1a*+), each only making up ∼1% of all RM epithelial cells prior to viral clearance, accumulated at 14 dpi (p=0.032 and p=0.067, respectively). The *Meg3*+MHC-II+ subset expressed high levels of maternally imprinted Meg genes (*Meg3*, *Rian*) alongside *Cd74*, H2 class II genes, *Wnt5a*, *Cxcl12*, and *Ccl25* (**Figure 3H**). They also uniquely expressed *Fezf2*, a transcription factor studied in the context of thymic self-antigen expression and immune tolerance induction by thymic epithelial cells (Takaba et al., 2015), but not *Aire,* which is necessary in the thymus for presentation of self-antigens. Whether these cells can promote tolerance in the upper respiratory tract, like their thymic counterparts, remains to be determined.

The *Krt13*+*Il1a*+ subset uniquely expressed *Krt13* (93.8% expressing within cluster vs 0.6% expressing in other clusters), a keratin previously described on “hillock” cells in the mouse trachea with undetermined functions (Montoro et al., 2018) and squamous/suprabasal cells in the human nasal turbinate (Deprez et al., 2020). These *Krt13*+*Il1a*+ cells exhibited several of the markers specific to tracheal hillock cells including *Lgals3*, *Ecm1,* and *Anxa1* but were not enriched for *Cldn3* or club cell marker *Scgb1a1*. Unlike hillock cells, this nasal subset expressed genes for several secreted and membrane bound immune cell regulatory factors including *Il1a*, *Tnf*, *Cd274* (PD-L1), *Ifngr2*, and *Cxcl16* as well as secretory proteins like *Defb1*, *Muc4*, and *Muc1* (**Figure 3I**). Flow cytometry confirmed the accumulation of Krt13+ cells 14 dpi (p=0.036, **Figure 3J**). Given the potential for these *Krt13*+ cells to communicate with immune cells, we stained the mucosa for Krt13 to understand their distribution throughout the nasal cavity. We found the strongest signal for Krt13 along the nasal floor in the anterior RM (**Figure 3K**) and more distally where the nasal mucosa meets the oral mucosa (**Figure S3I**). Comparing samples from naïve mice and 14 dpi, we saw increased Krt13 staining and colocalization with PD-L1 in the post-infection samples (**Figure 3L**). Thus, following resolution of IAV infection, rare subsets of nasal epithelial cells with immune communication potential accumulate in the RM.

### Neutrophils mature and activate in the RM immediately following IAV infection

Neutrophil accumulation in IAV infected lung has largely been associated with severe disease and poor prognosis (Brandes et al., 2013; Johansson and Kirsebom, 2021; Tang et al., 2019), but their role in the nasal viral infection is unknown. Given recent work showing a relationship between increased neutrophil activation in the nose prior to RSV infection and higher symptom occurrence (Habibi et al., 2020), and intrigued by the large number of neutrophils and mast cells captured across the nasal mucosa (n=7,987), we investigated the transcriptional programs and change in frequency of granulocytes in the nasal mucosa. Subclustering and UMAP embedding revealed a continuum of granulocyte development starting with granulocyte-myeloid precursor like cells (*Elane, Mpo)*, differentiating through immature states (*Camp, Mmp8, Retnlg*), and ending with several clusters of mature neutrophils (*Il1b, H2-D1, Siglecf*) (**Figures 4A, S4A, and Supplementary Table 1**). Mast cells expressing *Il6*, *Gata2*, and *Il4* were also detected in small numbers. Many neutrophils originated from OM samples (**Figures 2B and S4B**) and are likely present in high numbers in the bone marrow from that region. Pseudotime analysis across both the OM and RM recapitulated known maturation gene expression patterns in the blood (Grieshaber-Bouyer et al., 2021) and systemically (Xie et al., 2020) (**Figures 4B,C and S4C**). Precursors and immature neutrophils present largely in OM samples, likely in bone marrow, may give rise to activated and mature subsets in the RM that begin to accumulate in high frequencies only during infection.

**Figure 4:**
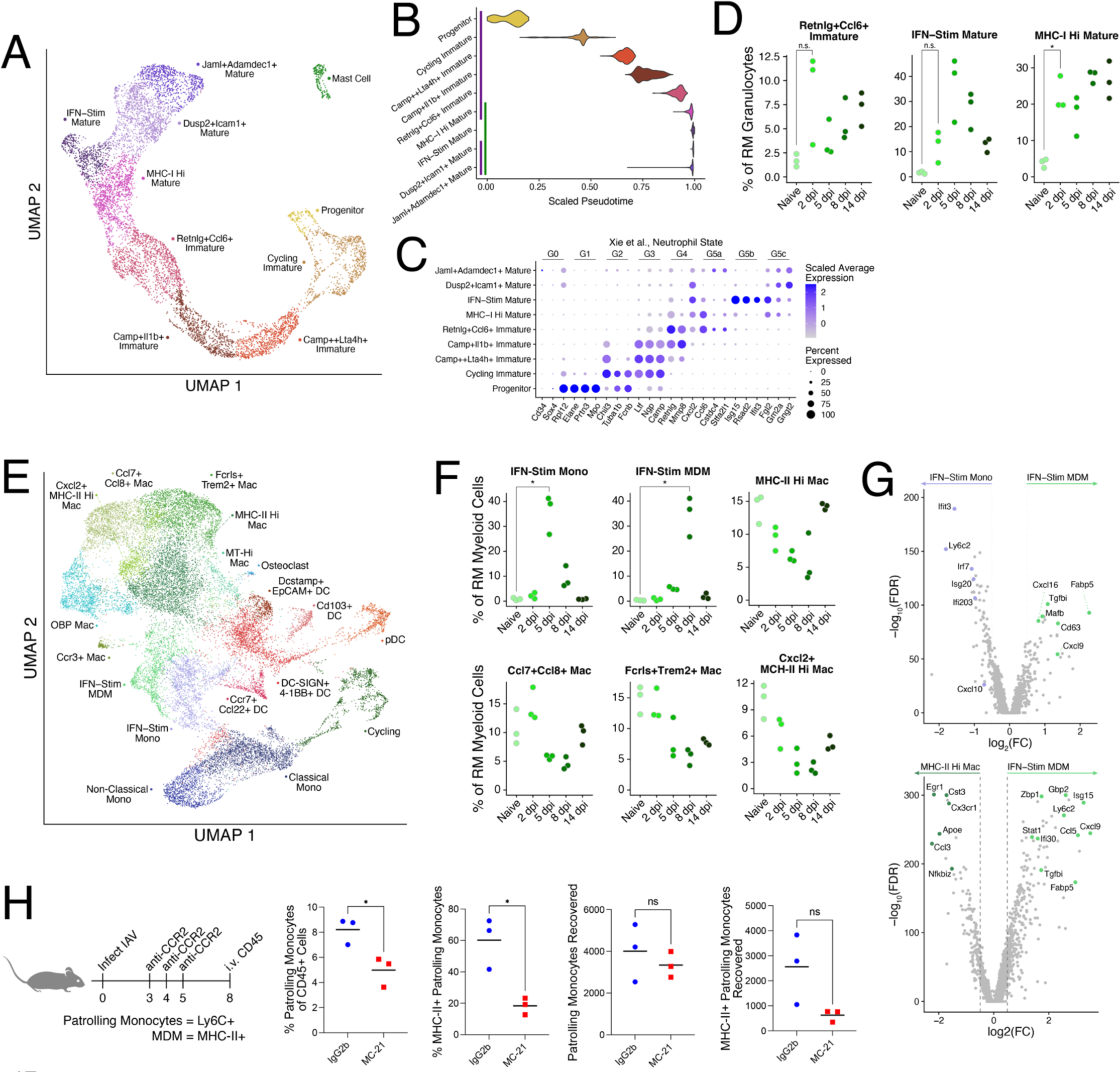
Immediate neutrophil responses are bolstered by recruited antiviral monocytes that differentiate into antiviral monocyte-derived macrophages. (A) UMAP embedding of 7,987 granulocytes across 10 clusters. (B) Violin plot of assigned pseudotime values to all granulocytes (except mast cells) split by cluster identity. Purple (OM) and green (RM) bars depict in which regions each cluster can be found at high frequencies (see **Figure S4B**). (C) Expression of blood neutrophil development genes annotated in (Xie et al., 2020) across nasal mucosa neutrophil clusters. (D) Relative frequencies of various neutrophil clusters as a proportion of all granulocytes per replicate RM sample. Only cells with assigned hash calls are included. Welch’s t test, *p < 0.05. (E) UMAP embedding of 22,654 macrophages, monocytes, and dendritic cells (DCs) across 17 clusters. (F) Relative frequencies of various myeloid cell clusters as a proportion of all macrophages, monocytes, and DCs per replicate RM sample. Welch’s t test, *p < 0.05. (G) Volcano plots depicting differentially expressed genes (|log2FC| ≥ 0.5; FDR < 0.01) between IFN-Stim MDMs and IFN-Stim Mono (top) and between IFN-Stim MDMs and MHC-II-Hi Macs (bottom). Only cells from RM were used in the differential expression analysis. Genes of interest are labeled. (H) Mice were infected with 10^4^ pfu IAV PR8 and then treated on 3, 4, and 5 dpi with either MC-21 (anti-CCR2; n=3) or IgG2b (isotype control; n=3). At 8 dpi mice received anti-CD45 intravascularly immediately prior to euthanasia to distinguish cells in the tissue from those in circulation. Patrolling monocytes were distinguished from circulating monocytes based on lower expression of Ly6C. Flow cytometry statistics are gated on CD45-EV+ cells; Welch’s t test, *p < 0.05.

By 2 dpi, neutrophil composition in the RM transitioned into mature IFN-stimulated and MHC-I-Hi states alongside an antimicrobial immature subset near the end of the pseudotime development trajectory (**Figure 4D**). The accumulation of these neutrophil clusters is one of the earliest changes in the RM following infection and may make up some of the earliest responses to local viral molecules and/or type-I IFN, with epithelial and RM-wide IFN-stim responses not arising until 5 dpi. Interestingly, the OM exhibited increased frequencies of mast cells, progenitors, and cycling immature granulocytes in 2-of-3 mice at 5 dpi, likely within bone marrow, indicating IAV infection may induce changes to adjacent hematopoiesis (**Figure S4D**). Thus, neutrophil activation and maturation mark the earliest detectable responses using our sampling strategy in the nasal mucosa to IAV infection.

### Stepwise recruitment of monocytes and differentiation of monocyte-derived macrophages follow neutrophil activation

Next, we explored heterogeneity among non-granulocyte myeloid cells (**Figure 4E, S5A and Supplementary Table 1**) — i.e., macrophages, monocytes, and dendritic cells (DCs). We captured a spectrum of macrophage (*Cd74, C1qb, Ccl4*) clusters including a *Trem2+* subset expressing *Fcrls, Il1a,* and *Pf4* (CXCL4), an innate immune recruiting subset expressing *Ccl7, Ccl8,* and *Pf4* (CXCL4), and a small cluster of osteoclasts (*Ctsk, Mmp9*). Monocytes clustered into classical (*Ly6c2, Ccr2, Chil3*) and non-classical (*Ace*, *Ear2, Itgal*) subsets (Jung et al., 2022) alongside IFN-stimulated monocytes and monocyte-derived macrophages (MDMs). DCs separated into distinct clusters including Langerhans-like (*Epcam, Ccl17, Ccl22*), intraepithelial (*Cd103, Xcr1, Tlr3*), migratory (*Ccr7, Ccl22, Cd274)*, and a subset uniquely expressing *Cd209a* (DC-SIGN), *Tnfsf9* (4-1BB), and *Klrd1*. We also captured plasmacytoid DCs (*Siglech, Irf8*; pDCs). Most myeloid clusters were present in all nasal regions with some exceptions like the IFN-stimulated monocytes and MDMs, which were restricted to the RM (**Figure S5B**).

While non-existent at baseline, upward of 30-40% of all myeloid cells belonged to these antiviral monocyte and MDM clusters 5 and 8 dpi, respectively (**Figure 4F**). The appearance and accumulation of monocytes and MDMs is concordant with lower frequencies of several tissue macrophage clusters, likely reflecting an overall increase in the total number of myeloid cells in the tissue as monocytes infiltrate from circulation. To understand the difference between the IFN-stimulated monocytes and MDMs, we performed differential expression analysis between clusters (**Figure 4G**). While the monocyte cluster had higher ISG expression than the MDM cluster, the MDMs still had relatively high ISG expression when compared with resting tissue macrophages. Notably, IFN-stimulated MDMs expressed higher levels of *Cxcl9* and *Cxcl16*, whose receptors (CXCR3 and CXCR6, respectively) have been implicated in TRM cell development in the lung (Slütter et al., 2013; Wein et al., 2019). Comparison of response scores to IFNα and IFNγ stimulation derived from in vitro macrophage cultures (Liu et al., 2012) on all IFN-stimulated monocytes and MDMs showed relatively higher expression of the IFNα score at 5 dpi and the IFNγ score at 8 dpi (**Figure S5C-D**), a similar pattern noted for IFN-stimulated epithelial cells (**Figure 3E**).

The IFN-stimulated MDM cluster also had the largest number of IAV+ cells of all myeloid cells (**Figure S5E**). Bystander analysis showed higher expression of some ISGs (*Isg15, Ifit3, Rsad2*) in IAV+ myeloid cells, like IAV+ epithelial cells (**Figure S5F**). However, the small fraction of IAV+ cells, compared to their bystander counterparts, had lower expression of MHC-II genes and other ISGs such as *Ccl6* and *Cxcl9* suggesting reduced antigen presentation and immune cell recruitment capacity. This is consistent with prior research in the lung showing that IAV suppresses myeloid cell activation and maturation (Moriyama et al., 2016; Zhang et al., 2022a).

The rapid shift in the myeloid compartment from IFN-stimulated monocytes at 5 dpi to a predominance of IFN-stimulated MDMs 3 days later (**Figure 4F**) suggested that recruited monocytes differentiated into MDMs within the RM during this interval. To test this idea, we treated mice with an anti-CCR2 antibody known to deplete circulating monocytes from the blood for 48 hours (Mack et al., 2001; Schneider et al., 2005) (**Figure S5G**). In the nasal mucosa, we stained for differentiating monocytes using MHC-II, CD11c, F4/80, CD64, and an intravascular CD45 stain to separate cells that had extravasated into the tissue from those in circulation (**Figure S5H**). Patrolling (Ly6C+) monocytes upregulate MHC-II, in addition to F4/80 and CD64, as they differentiate into MDMs. To deplete monocytes during their recruitment to the nasal mucosa (between 3-7 dpi), we treated animals with anti-CCR2 from 3 to 5 dpi. Depletion led to reduced frequencies of patrolling monocytes at 8 dpi (**Figures 4H**). Moreover, the proportion and number of MHC-II+ patrolling monocytes (i.e., differentiating MDMs) were considerably lower (18.4% vs 60.1% and 2558 vs 625, respectively), suggesting that the large proportion of IFN-stim MDMs measured at 8 dpi by scRNA-seq are derived from monocytes. Together, these data show that IAV infection induces a large recruitment of antiviral monocytes 5 dpi that differentiate into MDMs by 8 dpi in the RM.

### Antiviral NK cell responses precede transient effector T cells that are replaced by durable TRM cells following viral clearance

Following the accumulation of inflammatory and chemokine secreting monocytes and MDMs at 5 and 8 dpi, we anticipated a strong lymphocyte response during IAV infection. Thus, we next further investigated NK and T cells, the latter of which have been shown to be essential in clearing IAV infection in the lungs (Hufford et al., 2015) and nasal mucosa (Pizzolla et al., 2017). Subclustering revealed NK cell subsets (*Klrb1c, Ncr1*), type 2 (*Areg, Il13*) and type 3 (*Il22, Rorc*) innate lymphoid cells (ILC), γδT cells (*Trdc, Cd163l1, Cd3e*), and a spectrum of αβT cell subsets and states including naïve/central memory CD8 (*Ccr7, Dapl1*), effector CD8 (*Gzmb, Gzmk*), TRM-like CD8 cells (*Itgae* [CD103]), resting CD4 (*Cd4, Tnfrsf4* [OX40]), Th1 CD4 (*Ifng, Cd200*), Th17 (*Cd40lg, Il17a*), and a cluster of Helios (*Ikzf2*) expressing cells (**Figures 5A, S6A, and Supplementary Table 1**). Most T and NK cell clusters were enriched or restricted to the RM, but *Ccr7*+ CD8 T cells, ILCs, and γδT cells were also found in the OM and LNG (**Figure S6B**).

**Figure 5:**
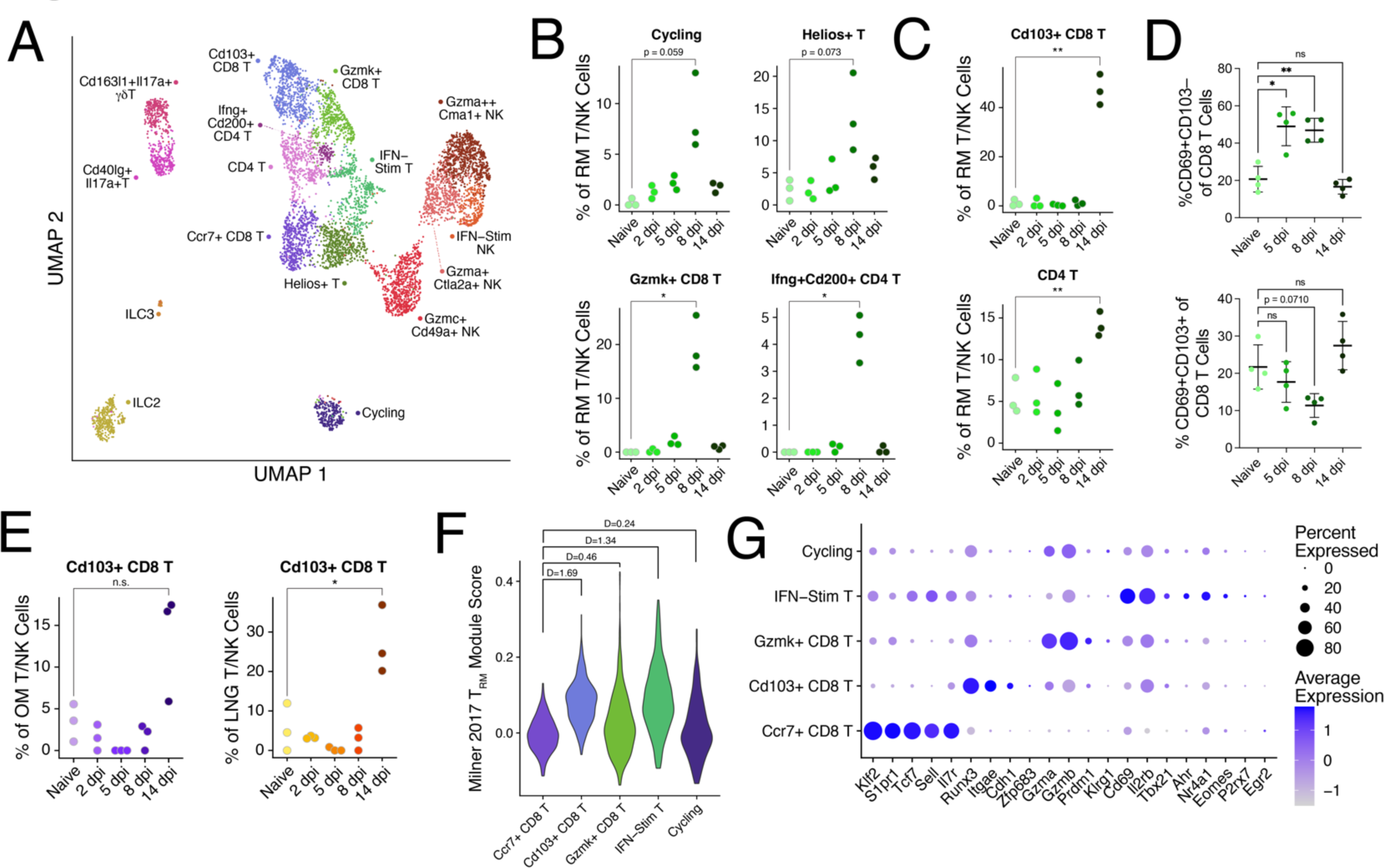
Effector CD4 and CD8 T cells 8 dpi are replaced by T_RM_ cells following viral clearance. (A) UMAP embedding of 6,573 T, NK, and innate lymphoid cells across 16 clusters. (B & C) Relative frequencies of various T cell clusters as a proportion of all T and NK cells per replicate RM sample. Only cells with assigned hash calls are included. (D) Mice were infected with 10^4^ PFU IAV PR8 and RM tissue was collected to stain for T cells. Kruskal-Wallis, *p < 0.05, **p < 0.01. (E) Relative frequencies of *Cd103*+ CD8 T cells as a proportion of all T cells, NK cells, and innate lymphocytes per OM replicate sample (left) and LNG replicate sample (right). (F) Violin plot depicting a gene module score derived from the universal T_RM_ signature as published in (Milner et al., 2017) across all CD8 T cell clusters for cells collected from RM. Cohen’s D for effect size is reported between *Ccr7*+ CD8 T cells and each other cluster. Welch’s t test, *p < 0.05, **p < 0.01. (G) Dot plot of genes encoding for canonical surface markers, proteases, and transcription factors enriched or absent in T_RM_ cells from circulating memory and naïve CD8 T cells. Gene list derived from (Crowl et al., 2022).

Looking at T and NK cell frequencies, we found that IFN-stimulated T and NK cells accumulated 5 dpi alongside cytotoxic NK cells that remained elevated through 8 dpi (**Figure S6C**). By 8 dpi, the T and NK cell compartment completely shifted towards effector antiviral T cell subsets, with high abundances of *Gzmk*+ CD8, Th1-like *Ifng*+*Cd200*+ CD4, cycling, and Helios+ T cell clusters (**Figure 5B**). These effector responses were short-lived, however, and were followed by increased frequencies of TRM cells and resting CD4 T cells (**Figure 5C**). Flow cytometry of RM tissue at matched time points confirmed an influx of CD69+CD103– activated CD8 T cells at 5 and 8 dpi that receded by 14 dpi alongside the accumulation of CD69+CD103+ TRM-like cells (**Figure 5D and S6D**). Notably, while detectable infection was largely restricted to the RM, TRM cells also increased in OM and LNG 14 dpi (**Figure 5E**), supporting the notion that even low levels of infection can result in TRM accumulation and development (Jiang et al., 2012).

To contextualize the TRM-like cells that arise following IAV clearance, we examined two recently published signatures separating resident memory from central/circulating memory. A universal TRM gene score (Milner et al., 2017) reasonably separated our TRM cluster from effector and naïve/memory CD8 T cells, but IFN-stimulated T cells also scored highly (**Figure 5F**). Conversely, the TRM-like cells also scored low for the associated circulating memory gene score, while naïve/memory CD8 T cells scored highest (**Figure S6E**). Closer inspection of known resident memory and central memory markers and transcription factors (Crowl et al., 2022) confirmed restriction of *CD103* (*Itgae*) expression to our TRM cluster, but *Cd69* was only highly expressed in IFN-stimulated cells. *Runx3* was also most highly expressed in TRM-like cells but also at lower levels in effector CD8 T cells. The *Cd103*+ CD8 cluster lacked the known naïve/central memory transcription factors *Klf2* and *Tcf7* (**Figure 5G**). In summary, effector T cell responses in the RM 8 dpi are replaced by TRM-like cells across all nasal mucosa regions following viral clearance.

### IgA+ cells populate throughout the NM following viral clearance

Following IAV infection, local mucosal plasma cells and activated B cells produce and secrete neutralizing soluble IgA into the airways (Rossen et al., 1970; Wellford et al., 2022; Woof and Mestecky, 2005). In the lungs, resident memory B cells form after primary infection and can be recruited upon secondary challenge to produce additional antibodies (MacLean et al., 2022), suggesting infection can lead to long term changes in both local and distal B cell subsets. Clustering of B cells in the nasal mucosa (**Figures S7A-B and Supplementary Table 1**) revealed mature subsets (*Ighd, Cd74*), IgG+/IgA+ early plasmablast cells (*Aicda, Jchain, Igha*), lambda-chain-high expressing cells (*Iglc1, Iglc2*), nucleoside diphosphate kinase (NME) expressing cells (*Nme1*, *Nme2*) and, primarily in OM, developing subsets including pro-B (*Dntt, Vpreb1, Rag1*), pre-B (*Bub1b, Mki67, Sox4*), and immature B cells (*Ms4a1, Ifi30*). The preponderance of precursor and developing B cell subsets in OM likely reflects bone marrow cells, whereas IgG+/IgA+ cells were found at highest frequency within LNG tissue, but class-switched B cells were also detectable in RM and OM (**Figure S7C**).

Looking at changes in cluster frequency over the course of IAV infection, we found that pro-B and pre-B cells collectively comprised up to 80-90% of all B cells in the OM 5 dpi increasing from 5-15% at baseline, suggesting that IAV infection may induce local B cell proliferation and differentiation in the bone marrow following infection and/or egress of mature B cells from this region (**Figure S7D**). This increase in B cell precursor frequency in OM 5 dpi paralleled that of granulocyte precursors (**Figure S4D**), supporting the notion of activation in nasal bone marrow. By 14 dpi, IgG+/IgA+ early plasmablast cells were detected in the RM in all three replicate samples. However, their recovery was more variable in OM and LNG samples, which could be due to biological variability and/or inconsistent cell capture. Flow cytometry staining for intracellular IgA confirmed the increase of IgA+ cells in both RM and LNG (**Figure S7E**) at 14 dpi. Moreover, imaging of the RM and LNG at 30 dpi confirmed the presence of IgA+ cells in both regions following infection (**Figure S7F**). Thus, B cells may undergo proliferative development during acute IAV infection and IgA+ plasmablasts accumulate in the RM and LNG following clearance.

### Proportionality guided cell-cell communication analysis highlights the CXCL16-CXCR6 signaling axis in effector and memory T cell responses

To understand how compositional changes in the tissue during primary infection may be coordinated across multiple cell subsets, we next characterized relationships between pairs of cell clusters over time using our compositional data. We employed proportionality analysis (Lovell et al., 2015), an alternative to correlation that avoids intra-sample abundance dependence present in compositional data (Quinn et al., 2018), to find cell clusters with significantly similar abundance trajectories (see **Methods**). Given that the RM was the major site of infection and showed temporally structured changes in cell composition over time (**Figure 2I**), we applied our proportionality analysis to all samples from this region. We discovered a highly structured proportionality landscape with 101 significantly proportional cluster pairs (FDR < 0.05) (**Figures 6A and S8A**). To understand coordination among larger groups of cell clusters, we built a network of all significantly proportional cluster pairs (**Figure S8B**). The network revealed larger groups of proportional responses made up of clusters from several different cell types and smaller and single-pair groups with 1-2 contributing cell types. Next, to further characterize the coordination among immune cells and between immune and epithelial cells, we investigated subsets of clusters with high proportionalities by cell-cell communication analysis.

**Figure 6:**
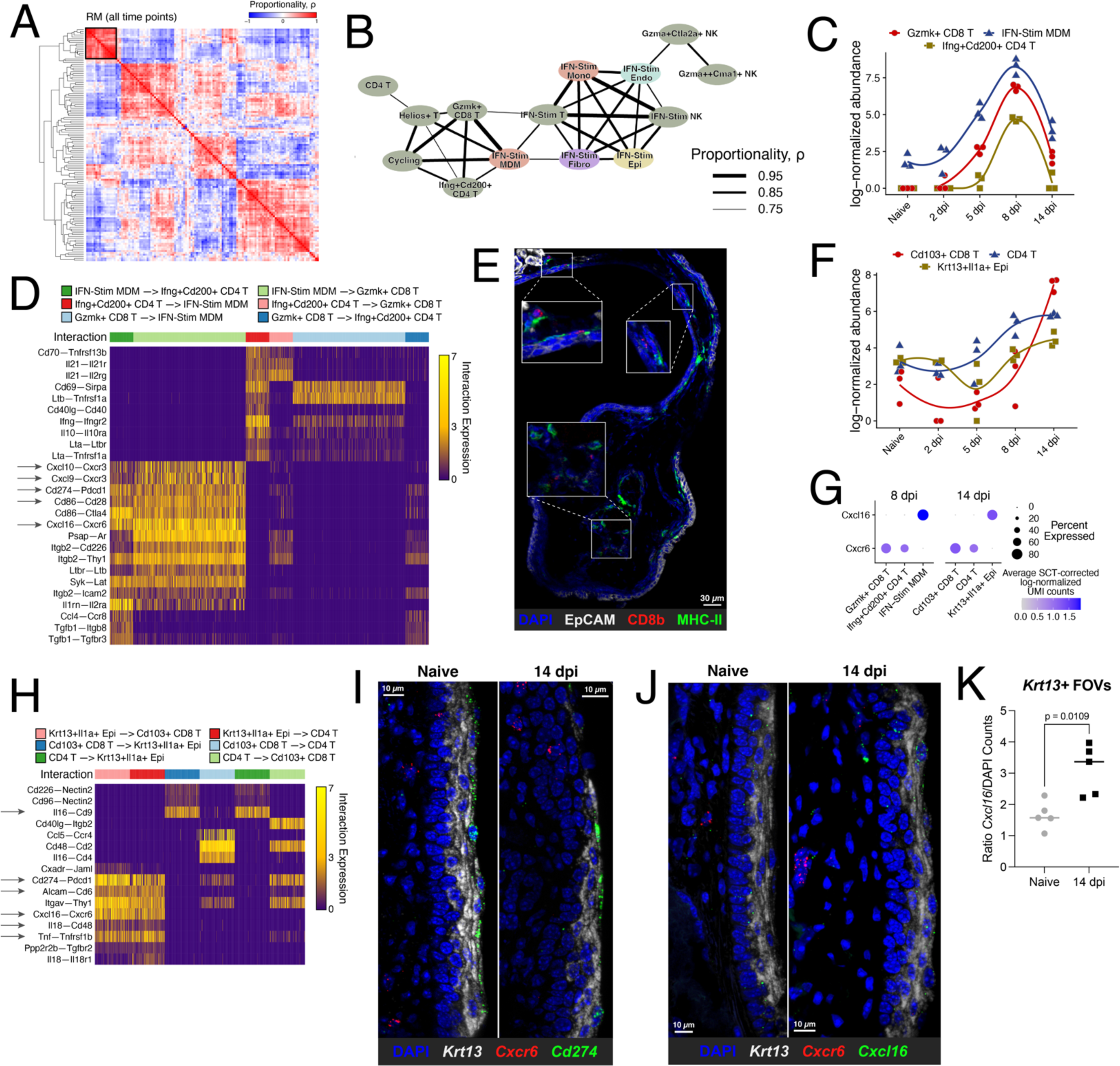
Proportionality and cell-cell communication analyses highlight the CXCL16–**CXCR6 signaling pathway in T cell:MDM and T cell:KNIIFE cell interactions**. (A) Hierarchically clustered similarity heatmap of sample replicate proportionality calculated across all RM primary infection time points. Black box surrounds the proportionality results for the cell clusters depicted in (B). (B) Network of significantly proportional (FDR<0.01) cell clusters as in (A). Nodes are colored by cell type and edge weight is representative of proportionality. (C) Abundance plot of *Gzmk*+ CD8 T cells, IFN-Stim MDMs, and *Ifng*+*Cd200*+ CD4 T cells in replicate RM samples. Smoothed lines are calculated using local polynomial regression fitting. (D) Heatmap depicting a subset of differentially expressed receptor-ligand interaction pairs between single-cell pairs identified by NICHES (Raredon et al., 2023) for the clusters depicted in (C); see **Supplementary Table 2**. Interaction expression is the multiplicative expression of receptor and ligand gene expression for each member of a single-cell pair. See for all receptor-ligand interaction pairs. Arrows highlight interactions described in the text. (E) Representative immunofluorescence imaging of maxillary turbinate at 8 dpi staining for EpCAM (gray), Cd8b (red), and MHC-II (green). White boxes show enlarged insets. (F) Abundance plot of *Cd103*+ CD8 T cells, CD4 T cells, and *Krt13*+*Il1a*+ epithelial (KNIIFE) cells in replicate RM samples. (G) Dot plot of scTransform-corrected log-normalized *Cxcl16* and *Cxcr6* expression 8 dpi (left) in the clusters depicted in (C) and 14 dpi (right) in the clusters depicted in (F). (H) Heatmap like (D) for the clusters shown in (F). (I) Representative RNAscope in situ staining for *Krt13* (gray)*, Cxcr6* (red)*, and Cd274* (i.e., PD-L1; green) of the nasal floor in a naïve mouse and 14 dpi mouse. Images depict a maximal intensity projection across 5 µm (10 slices) in the z-plane. (J) RNAscope as in (I) staining with *Krt13* (gray)*, Cxcl6* (red)*, and Cxcl16* (green). (K) Quantification of co-localization of *Krt13* and *Cxcl16* RNAs across multiple RNAscope images from the nasal floor (n = 5/timepoint). Ratio of the number of *Cxcl16* spots per nucleus within each *Krt13*+ region is reported. Welch’s t test, *p < 0.05.

#### IFN-stimulated MDMs – Gzmk+ CD8 T cells – Ifng+Cd200+ CD4 T cells

The strongest proportionality was observed among highly networked IFN-stimulated clusters and effector T cell clusters (**Figure 6B**). Given the strong myeloid and T cell responses 8 dpi and the possibility that activated MDMs may function as APCs, alongside DCs, within the nasal mucosa, we focused on the relationship between the IFN-stimulated MDM, *Gzmk*+ CD8 T cell, and *Ifng*+*Cd200*+ CD4 T cell clusters. Plotting abundance values confirmed synchronous trajectories of these three clusters with transient accumulation starting 5 dpi, peaking at 8 dpi, and waning by 14 dpi (**Figure 6C**). We next assessed cell-cell communication potential using NICHES, an approach that finds single-cell pairs with multiplicative high expression of known interacting ligands and receptors (Raredon et al., 2023). Differential ligand-receptor expression between groups of cell-pairs was then used to identify interactions specific to pairs of clusters (see **Methods** and **Supplementary Table 3**). Applied to cells from these three clusters at 8 dpi, we found several predicted, literature supported, interactions between the MDM cluster and both effector T cell clusters including *Cd274–Pdcd1* (PD-L1–PD-1), *Cd86–Cd28*, *Cxcl9/10–Cxcr3,* and *Cxcl16–Cxcr6* (**Figure 6D**). Imaging confirmed the spatial proximity of CD8 T cells and MDMs/DCs RM 8 dpi (**Figure 6E**).

#### Cd103+ DCs – Dusp2+Icam1+ mature neutrophils – Gp2+Lyz2+ goblet/secretory cells

The second largest networked group included various myeloid, granulocyte, and epithelial cell clusters. Intrigued by the inclusion of the late arising Gp2+Lyz2+ goblet/secretory cell cluster (**Figure S3G**), we took a closer look at this cluster and the two clusters most proportional with it: *Cd103*+ DCs and *Dusp2*+*Icam1*+ mature neutrophils. Plotting cluster abundances confirmed that all three clusters peaked following viral clearance 14 dpi (**Figure S8C**). Closer inspection of differentially expressed ligand-receptor pairs revealed potential pro-inflammatory signaling by Gp2+Lyz2+ Gob/Sec cells to mature neutrophils via *Sftpd–Sirpa*, which blocks Cd47 binding (Gardai et al., 2003), *Cirbp–Trem1* which has been shown to occur in sepsis (Denning et al., 2020), and *Tgfb2–Tgfbr1*. DC–neutrophil interactions included *Ccl2–Ccr1* and *Il18–Il18rap* suggesting mutual immune recruitment/homeostasis (**Figure S8D**). These interactions suggest that late arising *Gp2*+*Lyz2*+ goblet/secretory cells may recruit and regulate *Cd103*+ DCs and mature neutrophils in the RM following viral clearance.

#### Krt13+Il1a+ epithelial cells express Cxcl16 and increase when TRM-like cells accumulate

Like IFN-stim MDMs producing *Cxcl16* 8 dpi in concert with abundant *Cxcr6* expressing effector CD8 and CD4 T cells, the late arising *Krt13*+*Il1a*+ epithelial cell subset expressed *Cxcl16* alongside increasing frequencies of *Cxcr6+* TRM and CD4 T cells 14 dpi (**Figures 6F-G**). Although not significantly proportional over all time points (π = 0.53, 0.32, 0.77), we applied cell-cell communication analysis to cells from 14 dpi in these three clusters given the role of CXCL16– CXCR6 signaling in TRM localization in the lower airways (Morgan et al., 2008; Wein et al., 2019). In addition to discovering the *Cxcl16-Cxcr6* interaction, the analysis also captured additional interactions like *Cd274–Pdcd1*, *Tnf–Tnfsrsf1b*, *Il18–Cd48*, *Alcam–Cd6*, and *Il16–Cd9* (**Figure 6H**). RNAscope of the nasal floor in a naïve mouse and 14 dpi mouse confirmed expression of *Cd274* (PD-L1) and *Cxcl16* by *Krt13*+ cells in the vicinity of cells expressing *Cxcr6* (**Figure 6I-J**). Comparison between time points showed higher relative abundance of Cxcl16 transcripts within *Krt13*+ regions at 14 dpi compared to naïve samples (**Figures 6K and S8E**). Considering the transcriptional programming and localization of these cells, we propose the name ***K****rt13*+ **n**asal **i**mmune-**i**nteracting **f**loor **e**pithelial (KNIIFE) cells. Notably, KNIIFE cells are a fraction of many cells on the nasal floor expressing *Cxcl16* at 14 dpi, suggesting that this region of the nasal mucosa may be important in instructing TRM cells following viral clearance and/or tissue damage.

In summary, proportionality analysis coupled with cell-cell communication approaches reveal temporally synced cell cluster abundance changes over the course of primary infection and highlight potential cell-cell interactions contributing to T cell function and residual inflammation following viral clearance.

### Evaluating the reference capacity of the nasal mucosa atlas to learn compositions of additional scRNA-seq datasets

To assess the ability of our primary IAV infection atlas to contextualize and analyze additional scRNA-seq data generated from murine nasal mucosa, we leveraged the replicate structure of our dataset to test label transfer methods. Separating one RM replicate from each timepoint as a query set, we compared Seurat’s built-in weighted nearest neighbors method (Hao et al., 2021) to the generative model approach used in single-cell Annotation using Variational Inference (scANVI) (Xu et al., 2021). First trying cluster annotation on the entire RM query dataset, we found poor accuracy in assigning correct cluster identity in both methods and several clusters were completely lost in the predicted annotations (**Figure S9A**). Since the primary infection atlas clusters were found following multiple rounds of clustering, we next applied the same stepwise approach for label transfer: assign a cell type label and then split into cell types for cluster label predictions (**Figure 7A**). Using this stepwise approach, we correctly labeled 99.66% cell types using Seurat and 99.27% using scANVI (**Figure S9B**). Cluster identity calling, however, was more accurate in Seurat with 89.11-95.19% of cells correctly annotated across cell types, whereas scANVI had 69.33-92.31% properly labeled (**Figure S9C**). Moving forward with Seurat given its superior predictions, we repeated the analysis two more times using the other sets of replicates as the query dataset, finding robust reproducibility across models for cell type and cluster calling (**Figure S9D-E)**. Finally, we validated the output by calculating cell cluster abundances using the predicted cell cluster labels and projecting these “query” replicates into the primary infection RM compositional PCA. Remarkably, the query replicates aligned very closely to their real sample replicate counterparts in compositional space (**Figure S9F**). Thus, we validated a label transfer approach to learn cell cluster identities and cellular composition of new scRNA-seq data using our primary infection atlas as a reference.

**Figure 7:**
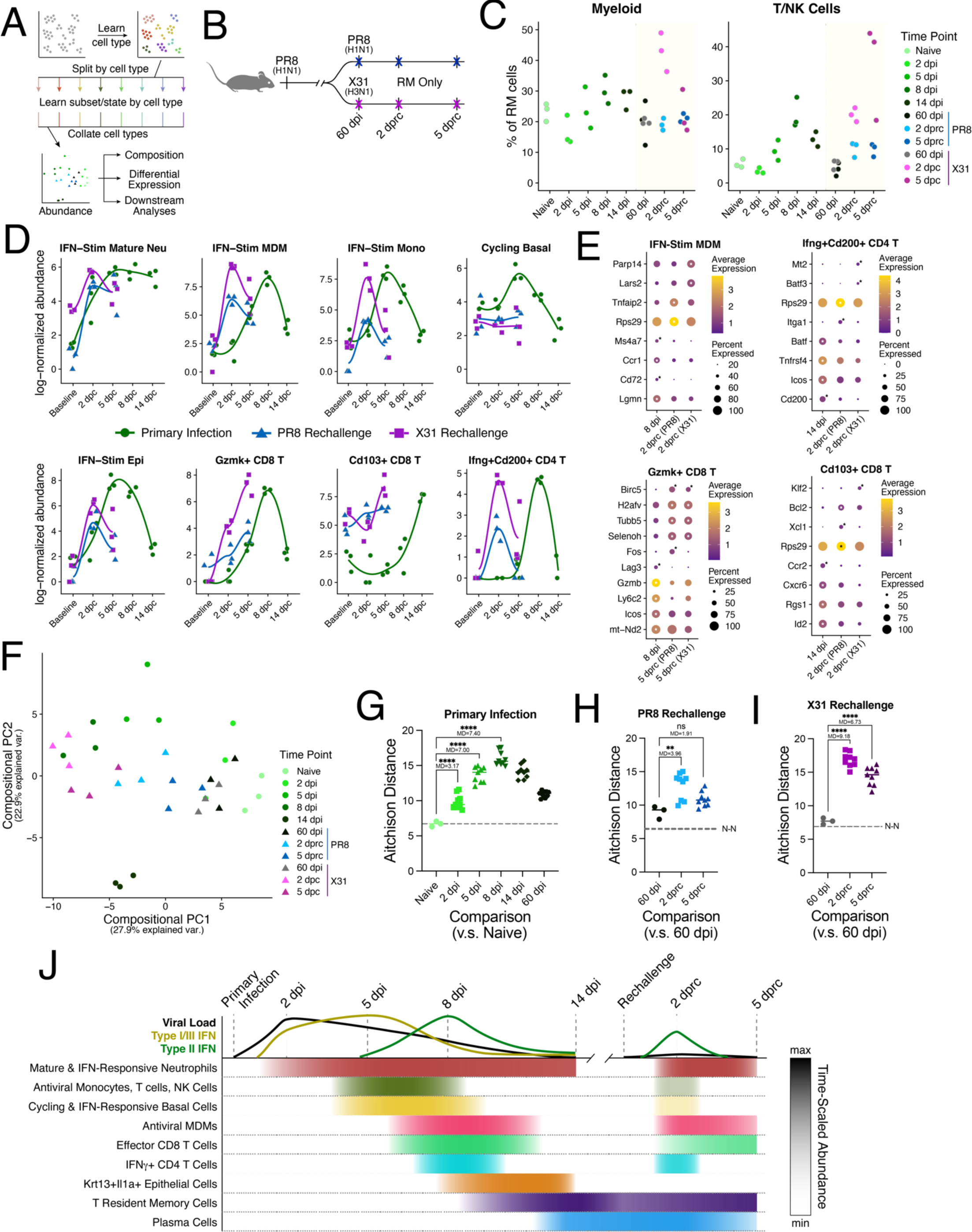
IAV rechallenge induces accelerated and coordinated memory immune responses. (A) Analysis scheme applied to RM samples to learn cell cluster identity. (B) Schema depicting experimental design for IAV rechallenge experiments. Mice previously infected with PR8 were administered either PR8 (H1N1) or X31 (H3N2) 60 dpi and RM was sampled prior to rechallenge, 2 and 5 days post rechallenge (dprc). (C) Relative frequencies of myeloid cells and T & NK cells as a proportion of all sequenced cells per RM replicate sample in primary infection and following rechallenge. (D) Abundance plots of various clusters showing overlaid primary infection (green) and rechallenge responses with PR8 in blue and X31 in pink. Baseline refers to samples from naïve mice in primary infection and to samples from 60 dpi in rechallenge. dpc = days post challenge. Smoothed lines are calculated using local polynomial regression fitting. (E) Dot plots depicting select cluster-specific and differentially expressed genes between primary infection and homologous and heterologous rechallenge in IFN-Stim MDMs (top left), *Ifng*+*Cd200*+ CD4 T cells (top right), *Gzmk*+ CD8 T cells (bottom left), and *Cd103*+ CD8 T cells (bottom right). Time points were chosen from peak responses in each challenge. Significantly enriched genes are labeled at each timepoint (*FDR<0.01). See **Supplementary Table 4** for all differentially expressed genes in each comparison. (F) Compositional PCA of primary infection and secondary challenge RM sample replicates. Here, secondary challenge samples were projected into the PC space calculated across only primary infection samples (see Figure 2H). (G, H, & I) Euclidean distances calculated between center log ratio transformed abundance values (Aitchison distance). The farther the Aitchison distance between two sample replicates, the less similar their compositions. (G) All pairs of naïve replicates and primary infection + 60 dpi replicates. (H) All pairs of 60 dpi replicates with matched PR8 rechallenge replicates. (I) All pairs of 60 dpi replicates with matched X31 rechallenge replicates. Dotted line plotted at the median naïve-naïve Aitchison distance (N-N). P values reported for multiple hypothesis corrected Welch’s ANOVA. MD = mean difference; *p<0.05; **p<0.01; ***p<0.001, ****p< 0.0001. (J) Timeline schematic of primary IAV infection and rechallenge depicting viral load trajectory, IFN dynamics, and immune and epithelial cell cluster response timing and duration.

### IAV rechallenge is characterized by accelerated and concurrent myeloid and lymphocyte memory responses

Having developed a tissue-scale response timeline of acute IAV infection in the nasal mucosa, we next asked how strain matched and unmatched induced memory responses differ from primary infection on cluster and compositional levels. After priming mice with IAV PR8 infection, we rechallenged 60 dpi with either the same virus or IAV X31, a H3N2 strain (**Figure 7B**). In the PR8→PR8 arm, both antibody and T-cell mediated mechanisms of protection will occur; in the PR8→X31 arm, however, antibodies from the primary infection will fail to neutralize IAV, but T cells will still exhibit memory responses (Pizzolla et al., 2017). Sampling RM prior to rechallenge at 60 dpi, and 2- and 5-days post rechallenge (dprc), we applied our label transfer approach — using the primary infection data as a reference — to annotate an additional 76,159 cells with cluster labels derived from primary infection (**Figures 7A**). All cell types present in the primary infection dataset were captured in the rechallenge samples and UMAP projection showed strong overlap between the datasets (**Figure S10A,B**). Plaque assays following PR8 rechallenge detected infectious virus in 1-of-5 mice in nasal mucosa, and in 0-of-6 mice in lung at 2 dprc, and none at 5 dprc, suggesting immediate control of infection or baseline resistance (**Figure S10C**).

Inspection of cell type frequencies in rechallenge demonstrated substantial accumulation of immune cells in both the homologous and heterologous settings. Even with neutralizing antibody mediated protection in the PR8 challenge and nearly undetectable viral titers, we measured increased proportions of granulocytes, T & NK cells, and B cells following rechallenge (**Figures 7C and S10D**). Following X31 challenge, T & NK cell frequencies increased even higher than following PR8 challenge alongside a substantial but transient accumulation of myeloid cells, supporting a bigger role for T cell-mediated responses in the absence of antibodies. Since plasma cells were not captured in our primary infection time course and thus could not be identified by label transfer, we re-clustered all the B cells in our acute and memory dataset. We readily resolved a small cluster (n = 75 cells) of plasma cells expressing *Slpi*, *Jchain*, *Igha,* and *Xbp1* present at low, variable frequencies at 60 dpi and during rechallenge (**Figures S10E,F**).

Following annotation, we compared changes in cluster abundance over time between the primary and secondary responses to IAV infection (**Figures 7D and S10G**). Like primary infection, the IFN-Stim neutrophil subset accumulated immediately and maintained elevated levels through 5 dprc. Interestingly, in homologous rechallenge IFN-stimulated MDMs rapidly accumulated while monocytes only slightly increased; in heterologous rechallenge, both increased substantially. Given the total proportion of myeloid cells in the dataset was similar between 60 dpi, 2 dprc, and 5 dprc following PR8, these data suggest that MDMs already inside the RM prior to homologous rechallenge quickly responded. Effector Th1 CD4 T cells were also elevated 2 dprc, but effector CD8 T and TRM cells were slower to accumulate. Cycling basal cells showed no change in abundance following rechallenge regardless of rechallenge strain, and the increase in IFN-responsive epithelial cells was also stunted. Notably, both the *Meg3*+*MHC-II*+ subset and KNIIFE cells that arose following viral clearance in primary infection remained at low levels throughout rechallenge. While we do not yet understand whether their roles may be restricted to resolving primary infection, there may be a necessary inflammation threshold for their expansion, or they may take longer to increase in frequency than the relatively early sampling timepoints post re-challenge. In summary, both PR8 and X31 rechallenge induced the rapid accumulation of several anti-viral myeloid and T cell subsets and states despite little detectible virus, with even greater induction following heterologous challenge.

To understand if cell state and the quality of antiviral effector responses differs depending on prior exposure and viral strain, we performed differential expression analysis within cell clusters across primary infection, PR8, and X31 rechallenge (**Figure 7E** and **Supplementary Table 4**). IFN-stimulated MDMs at 2 dprc in both PR8 and X31 rechallenge compared to 8 dpi had lower expression of *Lgmn, Cd72, and Ccr1*, but higher levels of *Tnfaip2*, *Lars2*, or *Parp14*. *Ifng*+*Cd200*+ CD4 T cells may become more prone to cell death during a memory response, with lower levels of *Tnfrsf4* (OX40), *Icos*, and *Cd200*. However, they expressed higher levels of *Itga1*, which has been shown to mark a subset of CD4 T cells that rapidly secrete IFN-γ in the airways following IAV infection (Chapman and Topham, 2010). Compared to 8 dpi, *Gzmk*+ CD8 T cells at 5 dprc in both PR8 and X31 rechallenge exhibited reduced expression of cytotoxic and activation genes, but higher levels of cell survival genes *Birc5* and *Selenoh*, and histone *H2afv*, suggesting induction of epigenetic modifications. During rechallenge, Cd103+ CD8 T cells expressed lower levels of *Id2*, *Cxcr6*, and *Ccr2* relative to 14 dpi, when these cells were likely just starting exhibit a TRM phenotype; during rechallenge, expression of *Xcl1*, *Bcl2*, and *Klf2* was moderately, but significantly increased.

While changes in abundance or gene expression on an individual cell subset/state level highlight specific differences between primary and secondary responses, we sought to understand how the collective RM tissue-scale response differs. To contextualize on the compositional level, we projected the memory and rechallenge sample replicates into the previously derived compositional PC space for RM in primary infection (**Figure 7F**). The RM 60 dpi samples were separated from naïve samples and most resembled 2 dpi, suggesting that even though IAV was cleared by 14 dpi, the nasal mucosa sustained significant changes in composition. For both memory arms, the 2 dprc time point recapitulated the variance described by PC1, with X31 rechallenge samples moving further negative in PC1 and overlapping 8 dpi. No significant shift along PC2, unlike in primary infection, suggests increases in effector immune responses occur upon rechallenge but not broad antiviral activation across all cell types like those seen 5 dpi (i.e., IFN-Stim clusters) (**Figure 2H**). Notably, by 5 dprc in the PR8. rechallenge, the tissue had almost returned to “memory baseline” at 60 dpi in PC-space, indicating that responses had already largely resolved; however, in the X31 challenge, a high level of immune infiltration is still taking place at 5 dprc. To assess variation between the primary infection distinct secondary response arms, we re-ran PCA with abundances from all timepoints, finding some separation between experiments, potentially arising from variation in the primary infection (**Figure S10H**). Nevertheless, infection in the naïve setting and both rechallenge paradigms resulted in a concerted and parallel directional shift in compositional PC space, supporting the conclusions of induced effector myeloid and T cell clusters from the projected PCA. Notably, PC4 captured the aspects unique to a “optimal” memory response, with increased abundances of several B cell clusters during homologous rechallenge (**Figure S10I**).

To quantify the overall difference between timepoints, we calculated compositional Aitchison distances between all pairs of sample replicates (**Methods**). In primary infection, RM increasingly separated from the naïve state up through 8 dpi but then became closer as infection is resolved (**Figure 7G**). Corroborating the PCA, the nasal mucosa 60 dpi was still distinct from its naïve state. Upon rechallenge with PR8, RM also separated from 60 dpi; however, the extent of that difference (MD=3.96) was less than between naïve and 5 dpi (MD=7.00) and 8 dpi (MD=7.40) indicating that the memory response to the same IAV strain was more succinct (**Figure 7H**). In comparison, X31 rechallenge led to the largest distance increase (MD=9.18), highlighting the massive accumulation of immune cells induced without antibody-mediated protection (**Figure 7I)**. Comparing primary infection timepoints with peak memory response, each primary infection timepoint was substantially distinct from 2 dprc, suggesting that prior infection rewired the RM response to IAV infection (**Figure S10J**). Summarizing the primary and secondary responses to infection described here, we present a timeline of the key immune and epithelial cell responses during IAV infection and rechallenge illustrating that many of the stepwise changes seen in primary infection occur in a more coordinated and accelerated fashion in memory (**Figure 7J**).

## DISCUSSION

Comprehensively understanding airway mucosal immunity is an urgent unmet need in the face of emerging and recurring respiratory pathogens (Lavelle and Ward, 2022; Morens et al., 2023; Roth et al., 2018; Russell et al., 2020). In particular, the nasal mucosa is at the forefront of mammalian host responses to airborne pathogens and functions as both an entry site and the primary barrier for infections of the respiratory tract. Consequently, the nasal mucosa is thought to be the site of initial engagement of respiratory viruses to generate both local T cell memory (Pizzolla et al., 2017) and neutralizing antibodies (Liew et al., 2023; Sterlin et al., 2021; Wellford et al., 2022; Weltzin et al., 1996). Determining how these responses occur following primary infection, and how immune and non-immune cells in the nasal mucosa contribute to viral clearance and subsequent memory, is critical to inform the design of next-generation nasal vaccines and therapeutics.

Here, we present a longitudinal, multi-region, scRNA-seq atlas of the murine nasal mucosa during primary and secondary IAV infection. Cataloguing the distribution and temporal dynamics of the diverse cell types, subsets, and states present, we develop and apply a compositional framework to understand tissue-scale changes occurring throughout primary and memory responses to viral infection. Neutrophil activation responses following infection precede broader type I/III IFN-stimulated responses in epithelial, myeloid, and lymphoid cells. By 8 dpi, effector CD8 and *Ifng*+ CD4 T cell subsets accumulate alongside recently differentiated MDMs. Following viral clearance at 14 dpi, TRM-like cells and IgG+/IgA+ B cells appear in the nasal mucosa, which has achieved distinct cellular composition from the naïve state with these adaptive immune subsets being sustained until 60 dpi. Careful investigation of the epithelial cell compartment also revealed a rare, previously undescribed KNIIFE cell subset. We validate and localize the presence and accumulation of these cells, provide evidence for their interaction with *Cxcr6*-expressing lymphocytes, and show co-expression of *Krt13*, *Cd274* (PD-L1), and *Cxcl16* on the nasal floor following viral clearance. Employing the primary infection atlas to annotate and interpret new secondary infection samples using both homologous and heterologous IAV strains showed that rechallenge induces surprisingly widespread yet coordinated and accelerated changes to cellular composition. We identify accelerated neutrophil, macrophage, and T cell responses in memory with a reduced burden on epithelial cells to express the joint interferon and proliferative response programs of primary infection.

The nasal mucosa consists of a multi-faceted epithelium that exhibits diverse responses to IAV infection. Basal cells in the nasal mucosa give rise to epithelial cells reminiscent of pseudostratified epithelium found in the trachea (Davis and Wypych, 2021), but we and others have also captured several additional epithelial cell subsets not found in other parts of the airway. Our atlas validates recent work in mice describing sustentacular cells, ionocytes, nasal tuft cells, and serous cells (Brann et al., 2020; Ualiyeva et al., 2024). Moreover, we describe for the first time several clusters of epithelial cells with undetermined function: (1) *Scgb-b27+Cck+*, (2) *Meg3+*MHC-II+, (3) *Klk1+Fxyd2+*, and (4) KNIIFE cells. Except for the *Scgb-b27*+*Cck*+ cluster, the remaining previously undescribed clusters all exist at low frequencies (< 1% of all epithelial cells) in naïve mice and may have been missed in experiments without sufficient cell numbers or utilized cell sorting.

The viral signaling and proliferative capacities of epithelial cells associate with COVID-19 disease trajectory (Sposito et al., 2021; Ziegler et al., 2021). Our data confirm that transient IFN-responsive epithelial cell subsets, including cycling basal cells, arise in the nasal mucosa during IAV infection. At 5 dpi, type I/III IFNs drive the response, while at 8 dpi, IFNγ levels are increased alongside Th1-like cells. At 5 dpi, nasal basal cells co-expressed cell cycle and IFN-response programs, which have been previously described as non-compatible in lungs (Broggi et al., 2020; Major et al., 2020). Given the diverse roles of nasal epithelial cells and the need to protect olfactory sensory neurons (Dumm et al., 2020), nasal basal cells may be more tolerant of IFN-response signaling during proliferation than basal cells in the lower airways. Notably, epithelial IFN-induced responses were significantly reduced upon rechallenge regardless of IAV strain. This difference could reflect several non-exclusive mechanisms during recall, including an overall reduction or shortening in IFN production or signaling, lower viral load, or a potential tolerized basal cell state. Airway basal cells can develop transcriptional memory in vitro (Adamson et al., 2022), but whether primary infection can confer durable memory to viral immunity, as has been seen for allergic inflammation (Ordovas-Montanes et al., 2018), requires further investigation.

In addition to epithelial responses, IAV infection also precipitated a highly dynamic, stepwise response by immune cells that was initially dominated by myeloid subsets. Specifically, we observed a substantial early influx of neutrophils followed by monocytes responding to type I/III IFNs that then differentiated into MDMs concurrent with the arrival of effector T cells expressing IFNγ. The evidence for both resident (Yao et al., 2018) and recruited (Aegerter et al., 2020) macrophages in the lungs to engage in memory responses suggests that a similar phenomenon may occur in the nasal mucosa. After depleting circulating monocytes during acute infection, MDM formation was markedly reduced, indicating that the majority of nasal MDMs at 8 dpi differentiated from newly recruited monocytes. Interestingly, during homologous memory response MDMs, but not monocytes, increased in abundance 2 dprc even though overall myeloid frequencies remained unchanged, suggesting that either recruited MDMs replaced local myeloid cells or MDMs already present in the tissue expanded to exert antiviral effector functions. If the latter, understanding the mechanisms by which enhanced myeloid function is maintained and recalled in the nasal mucosa could yield a new avenue for designing improved mucosal vaccines (Iijima and Iwasaki, 2014).

The role of adaptive immune responses to IAV infection have been well described in the lower respiratory tract (Chapman and Topham, 2010; Krammer, 2019; McMaster et al., 2015; Onodera et al., 2012; Slütter et al., 2013; Wein et al., 2019), but their dynamics and quality in the nasal mucosa are less understood. Antibody-mediated immunity following primary and secondary IAV infection has been described (Chen et al., 2018) and falls outside the scope of the present study. Antibody-producing B cell response dynamics were variable in our model with sizeable frequencies of IgA+ B cells detected in 1- or 2-of-3 mice at 14 dpi and low numbers of plasma cells at 60 dpi during rechallenge. Flow cytometry validated increased frequencies of IgA+ cells in RM and LNG at 14 dpi. Wellford et al., recently showed in an influenza B model that the OM requires mucosa resident plasma cells to prevent transmission to the brain; the RM, alternatively, can receive neutralizing antibodies from both serum and local plasma cells (Wellford et al., 2022). To what extent local nasal plasma cell derived IgG and IgA play roles in stymying infection during rechallenge, and whether non-neutralizing antibody functions help activate other immune subsets (e.g., Fc receptor mediated signaling), must be further explored.

IAV infection of the nasal mucosa resulted in classical T cell responses with both antiviral effector CD8 T cells expressing cytotoxic genes and Th1 CD4 T cells expressing *Ifng* and *Tnfrsf4* (OX40) arising at 8 dpi. Proportionality analysis revealed their coordination with an influx of IFN-Stim MDMs expressing *Cxcl9* and *Cxcl16*, and NICHES predicted several modes of communication between all three clusters, suggesting MDMs, alongside DCs, may provide activation signals for T cells in the nasal mucosa. While infectious titers waned between 2 and 8 dpi, effector T cell responses likely played a critical role in completely extinguishing IAV infection by 14 dpi. Finding upward of ∼50% of all T and NK cells by 14 dpi belonged to the TRM-like cluster in the RM tissue, we validate their presence and phenotype in the nasal cavity following IAV infection (Pizzolla et al., 2017; Wiley et al., 2001). Unlike in the lung where TRM cells quickly wane following infection (Slütter et al., 2017), our data demonstrate robust frequencies out to 60 dpi that are further amplified during rechallenge; moreover, in addition to the RM, we find TRM in OM and LNG tissue, where virus is only detected at low levels or not at all. While TRM contribute to an effective memory response upon rechallenge (Ariotti et al., 2014; McMaster et al., 2015; Schenkel et al., 2014; Steinbach et al., 2016), differential expression across primary and rechallenge responses of the *Cd103*+ CD8 T cell cluster in our dataset showed few significant gene expression differences between time points; moreover, their relative proportion only increased by ∼2-3x over levels at 60 dpi depending on the strain of IAV used. Together, T cell accumulation was substantially higher in a heterologous rechallenge setting compared to the matched strain, suggesting potential compensatory mechanisms of protection without functional neutralizing antibodies. Given recent work highlighting the importance of TRM in mitigating nasal viral infections and transmission (Mao et al., 2022; Pizzolla et al., 2017; Uddbäck et al., 2024), understanding which cell subsets and signals establish, maintain, and expand the TRM niche could help guide mucosal vaccine strategies with heterotypic protection.

Following viral clearance, subsets of epithelial cells with potential immune signaling and inflammatory regulation capacity substantially increased in abundance. In addition to a large cluster of goblet/secretory cells with predicted DC/neutrophil communication ability, we also discovered a subset of epithelial cells uniquely expressing *Krt13* and *Krt6a* in the nasal mucosa, which we identify as ***K****rt13*+ **n**asal **i**mmune-**i**nteracting **f**loor **e**pithelial (KNIIFE) cells. Phenotypically, KNIIFE cells were reminiscent of the recently described “hillock” cells in the trachea expressing *Krt13, Ecm1*, and *Lgals3* (Montoro et al., 2018) and squamous cells in human nasal swabs and biopsies (Deprez et al., 2020; Ziegler et al., 2021). However, KNIIFE cells also expressed several genes often found in macrophages including *Cd274* (PD-L1), *Ifngr2*, *Tnf*, and *Cxcl16*. This cluster was present at low levels throughout infection until expanding 14 dpi and remained stable during rechallenge. At 14 dpi, we measured *Krt13* and *Cxcl16* co-expression in situ nearby *Cxcr6* expressing cells, especially along the nasal floor. These results raise the possibility that KNIIFE cells, by providing a source for CXCL16 beyond that expressed by myeloid cells, may contribute to the establishment and/or maintenance of the resident memory T cell pool in the nasal mucosa, as has been suggested for this chemokine pathway in other tissues (Clark et al., 2006; Heim et al., 2023; Morgan et al., 2008; Tse et al., 2014; Wein et al., 2019). We use “KNIIFE” as a convenient acronym for these cells, but their specific functions must still be elucidated. The enrichment of KNIIFE cells along nasal floor and below the vomeronasal organ prompts the question of whether these cells interact with particles or irritants just entering or settling in the nose and play a regulatory role in tissue tolerance and/or immunity.

Compositional scRNA-seq analyses are becoming more common to discern differences between disease trajectories (Ordovas-Montanes et al., 2018; Smillie et al., 2019; Zheng et al., 2021), treatment groups (Darrah et al., 2020; Zhang et al., 2022b), and/or species (Chen et al., 2022; Li et al., 2022). Current tools focus on identifying specific clusters or gene programs that are compositionally distinct between groups (Büttner et al., 2021; Cao et al., 2019; Dann et al., 2022). However, the power of compositional scRNA-seq data lies in its structure; namely, singular changes in composition cannot be independent and must correspond with mutual changes in other clusters/programs. Leveraging the biological replicates and multiple timepoints present in our atlas, we utilized straightforward tools for compositional analysis adapted from microbiome research (Gloor et al., 2017; Lin and Peddada, 2020; Quinn et al., 2018) to understand tissue-scale changes within the nasal mucosa throughout IAV infection. PCA of center-log ratio transformed cell cluster abundances across sample replicates separated nasal regions and depicted structured stepwise changes in epithelial and immune cell subsets throughout infection trajectory. Proportionality analysis, which avoids the spurious associations present in Pearson correlation applied to compositional data (Lovell et al., 2015), revealed pairs and groups of clusters with significantly similar compositional trajectories (e.g., IFN-stimulated MDMs and effector CD4 and CD8 T cells) and can be readily applied to discover similarities across various metadata. We note that proportionality will highlight those potential interactions where subsets are changing together in relative abundance. Thus, any interactions that occur between subsets where only changes in gene expression occur may be overlooked by this approach, but newer tools are in development to assess prior interaction potential based on gene expression (Li et al., 2023). Finally, metrics like Aitchison distance (Aitchison et al., 2000) capture holistic changes in tissue-scale cellular composition and support standard tests for differences between group means (e.g., Welch’s ANOVA) to assess global similarity and compositional distance traveled by a tissue. Applied to our datasets, the RM “travels” less during an “optimal” memory response to IAV than during primary infection, suggesting prior infection induces a coordination of responses that were previously unsynchronized. We propose that these approaches for analyzing scRNA-seq data constitute a new framework for understanding and summarizing whole tissue- and biopsy-scale changes in cellular composition at high resolution in health, disease, and/or under perturbation.

Collectively, our murine nasal mucosa atlas of primary IAV infection longitudinally catalogues the cell types, subsets, and states present throughout distinct nasal regions. We demonstrate the utility of our dataset to serve as an annotation reference for newly generated scRNA-seq datasets and apply it to understand how the response to infection in the RM differs during memory recall following distinct IAV rechallenges. These findings will help contextualize temporal studies of the nose in humans with more complex exposure histories and highlight key immune and epithelial cell responses to recapitulate in future nasal vaccines and therapeutics to drive increased synchronicity in nasal memory responses.

### Limitations of the study

First, we acknowledge that cluster abundance-based compositional analyses are inherently dependent on how clustering was performed, and thus implicitly incorporates, to some degree, operator bias. While we believe our approach to be as impartial as possible through use of iterative clustering, it will be imperative to implement robust, reproducible clustering analyses (Hu et al., 2019; Patterson-Cross et al., 2021; Zheng et al., 2023) prior to compositional analysis moving forward. Partial labeling of cells by hashing antibodies may also have obscured changes in composition over time. Second, detection of IAV transcripts by scRNA-seq was limited. Other studies have included spike-in primers to facilitate additional capture of viral nucleic acids (Ratnasiri et al., 2023); it is possible that we were not sufficiently sensitive to IAV transcripts without these spike-in primers. Also, cells productively infected with virus may not be sufficiently viable through our tissue processing pipeline, as suggested by the viral reads detected by bulk RNA-seq from tissue lysate, leading to artificially low numbers of cells containing IAV reads. Third, to increase the relative proportion of non-epithelial cells in our scRNA-seq dataset, we performed a partial EpCAM depletion using magnetic beads. This decision was made following experiments comparing this approach to un-depleted RM tissue in naïve mice; we found that while depletion reduced the relative abundance of some epithelial cell clusters, it did not result in the complete loss of any cluster. Thus, the cellular compositions throughout the study represent the nasal mucosa tissue after both dissociation and epithelial cell depletion and, therefore, do not reflect the true frequencies of cell types within intact nasal mucosa. Nevertheless, our atlas can still be used to assign cell cluster labels to new datasets where epithelial cells have not been depleted and could inform spatial transcriptomics approaches to derive more accurate cell abundances in vivo. While this dataset represents the largest scRNA-seq atlas of the murine nasal mucosa to date, we may still be under sampling this complex tissue. Indeed, spatial transcriptomics and/or multiplexed immunofluorescence approaches will help validate the spatial organization and quantification of cell clusters defined here; however, given the complexity of the nasal mucosa and difficulty in sectioning through the nasal bone, further work will need to be done to validate, adapt and refine imaging protocols for this unique tissue. Additional experiments to test how other respiratory pathogens and vaccination strategies impact the composition and timing of responses in the nasal mucosa to IAV challenge are warranted (Rutigliano et al., 2010).

## Acknowledgements

We would like to thank members of the Ordovas-Montanes and von Andrian labs for insightful discussions and advice. S.W.K was supported by the Cancer Research Institute’s Irvington Postdoctoral Fellowship. J.O.M is a New York Stem Cell Foundation – Robertson Investigator. J.O.M was supported by the AbbVie-Harvard Medical School Alliance, the Richard and Susan Smith Family Foundation, the AGA Research Foundation’s AGA-Takeda Pharmaceuticals Research Scholar Award in IBD – AGA2020-13-01, the HDDC Pilot and Feasibility P30 DK034854, the Leona M. and Harry B. Helmsley Charitable Trust, The Pew Charitable Trusts Biomedical Scholars, The Broad Next Generation Award, The Chan Zuckerberg Initiative Pediatric Networks, The Mathers Foundation, The New York Stem Cell Foundation, NIH R01 HL162642, NIH R01 DE031928 and The Cell Discovery Network, a collaborative funded by The Manton Foundation and The Warren Alpert Foundation at Boston Children’s Hospital. This work was supported by the AbbVie–HMS Alliance Program. Sequencing of the scRNA-seq libraries was performed with AbbVie Inc. at the Genomics Research Center and Immunoprofiling Center in collaboration with Steven Leonardo (former AbbVie employee), Abel Suarez-Fueyo, Neha Chaudhary, Jozsef Karman (former AbbVie employee), Aridaman Pandit, and Amlan Biswas. Imaging was performed with MicRoN and HMS Center for Immune Imaging at Harvard Medical School. Anti-CCR2 antibody MC-21 and the X-31 IAV strain (A/HKx31) were generously provided by Prof. Matthias Mack (Universität Regensburg) and Prof. Daniel Lingwood (Ragon Institute of MGH, MIT, and Harvard) respectively. Training for the RNAscope experiments was facilitated by Anoohya N. Muppirala and Meenakshi Rao at Boston Children’s Hospital. Daniel Lingwood (Ragon Institute of Mass General, MIT, and Harvard), Alex Shalek (MIT), Paolo Cadinu and Jeffrey Moffitt (BCH), Colin Bingle (Sheffield) provided helpful feedback and discussion on experimental design and interpretation of the results. Additionally, we would like to thank Susan Westmoreland for insightful discussions as well as Shankar Subramanian, Michelle Cordoba Gunter, Isabel Chico-Calero, Jochen Salfeld, and Mark Namchuk for supporting the AbbVie–HMS Alliance Program Area 1 which enabled the scope of this work.

## Author contributions

Conceptualization, S.W.K., C.M., U.H.vA., and J.O-M,;

Methodology, S.W.K., C.M., and M.M.;

Software, S.W.K., E.M.L., and T.J.L.;

Formal Analysis, S.W.K., C.M., E.M.L., and T.J.L.;

Investigation, S.W.K., C.M., E.M.L., M.M., E.O., J.M., K.N., and J.O-M.;

Data Curation, S.W.K.;

Writing – Original Draft, S.W.K and C.M.;

Writing – Review & Editing, S.W.K., C.M., E.M.L., U.H.vA., and J.O-M.;

Supervision, U.H.vA. and J.O-M.;

Funding Acquisition, S.W.K., U.H.vA., and J.O-M.

## METHODS

### Resource availability

#### Lead contact

Further information and requests for resources and reagents should be directed to and will be fulfilled by the lead contact Jose Ordovas-Montanes.

#### Material availability

All the mouse lines used in this study are available from Jackson Laboratories. The anti-CCR2 antibody MC-21 was provided as a gift by Prof. Matthias Mack. This study did not generate new unique reagents.

#### Data and code availability

All sequencing data reported in this paper will be available in FASTQ read format and cellbender corrected gene expression matrix format at Gene Expression Omnibus upon publication as we plan to submit this work for pre-print. The annotated data can also be explored at the Broad Institute Single Cell Portal under study numbers SCP2216 and SCP2221. All the code generated and used to analyze the data reported in this paper will be available on GitHub in the jo-m-lab/IAV-nasal-sc-atlas repository.

### Experimental model and subject details

#### Mice

All experiments were approved by the Harvard University Institutional Animal Care and Use Committee and run following NIH guidelines. C57BL/6J (B6) mice 6 to 8 weeks old were purchased from The Jackson Laboratory and experiments commenced 1 to 3 weeks following their arrival. Mice were infected with 10^4^ pfu PR8 in a 10µL volume that was administered by pipette dropwise to the nares to allow each drop to be inhaled (5µl/nostril). Mice were restrained during this administration but not anesthetized, to maintain the virus in the upper respiratory tract. For rechallenge experiments mice previous inoculated with 10^4^ pfu PR8 were administered with either 10^4^ pfu PR8 or 10^5^ pfu X31 60 days after the initial PR8 infection. A higher dose of X31 was used in rechallenge given its reduced pathology (Rutigliano et al., 2014). All mice were housed in a BSL-2+ facility with specific pathogen free conditions.

### Method details

#### Virus growth, quantification, and mouse infections

IAV strain A/Puerto Rico/8/1934 (PR8) and Madin-Darby canine kidney (MDCK) cells were generously provided by Dr. Daniel Lingwood and Dr. Maya Sangesland of the Ragon Institute of Mass General, MIT, and Harvard. Virus was propagated and quantified in MDCK cells. MDCK cells were grown at 37°C with 5% CO2 in cell growth media: Dulbecco’s modified eagle’s medium (DMEM) (Corning, #10-017-CV), 10% fetal bovine serum (FBS; Gemini #100-106), 1X Penicillin:Streptomycin (Gemini, 100X stock: 400109). PR8 was grown in MDCK cells in influenza growth media: Iscove’s DMEM (Corning, # 10-016-CV), 0.2% bovine serum albumin (BSA; EMD Millipore, EM-2960), 1Xm Penicillin:Streptomycin, and 2µg/mL TPCK treated Trypsin (Sigma, T1426).

For viral load quantification experiments, mice were sacrificed in CO2 and lungs and heads were separated. For the nasal cavity, fur and skin were removed and the lower jaws cut off. The entire nasal cavity or lungs were collected into 1mL PBS with 2.3mm Zirconia/Silica beads (Biospec Products, 11079125z) and stored on ice. The tissue was homogenized in an OMNI Bead Ruptor Elite at 3m/s for 30 seconds twice, centrifuged 500g for 5 minutes, and supernatant was collected and stored at -80°C until thawed for plaque assays. Virus titers were measured by plaque assays in confluent MDCK cells in 6-well plates. MDCK cells were grown in cell growth media, washed with sterile phosphate-buffered saline (PBS), then washed with influenza growth media. Media was removed and serial dilutions of viral supernatant in influenza growth media were added to each well in a 400µL volume, incubated for 1 hour at 37°C, then overlayed with 0.3% agarose in influenza growth media. Infected cells were incubated for three days at 37°C, fixed with 4% paraformaldehyde, stained with crystal violet, washed, and plaques were counted.

### Tissue harvesting, single-cell suspension preparation, and hashtag labeling

Three separate regions of the nasal tissue were harvested independently: (1) the respiratory mucosa (RM), (2) the olfactory mucosa (OM), and (3) the lateral nasal gland (LNG). The nasal tissue was collected by removing the skin and connective tissue from around the head, cutting off the lower jaw, and opening the nasal cavity by peeling away the nasal bone from the rest of the skull. Tissue separation and collection was performed using a dissection scope with a 4x objective. All nasal tissue surrounding the nasoturbinates, maxillary turbinates, and septum, including the mucosa that runs along the nasal lateral walls between the nasoturbinates and maxillary turbinate, and the mucosal tissue under the nasal bone that connects the nasoturbinates and septum, were collected together and constitute the RM. After removal of the RM the ethmoid turbinates were collected including both the mucosal tissue and the bone and cartilage of the turbinates, but not the surrounding skull, constituting the OM. After removal of the OM, the LNG was exposed and could be collected without any bone or cartilage. The nasal-associated lymphoid tissue (NALT) was not collected in any of the three regions. Matched RM, OM, and LNG regions were collected simultaneously from the three mice per time point, and each time point was processed on a different day.

Each nasal tissue region was collected into 750µL Wash Media (RPMI 1640, 2% FBS, 10 mM HEPES, and 100U/ml penicillin G, 100µg/ml streptomycin) and stored on ice. Tissues were chopped with scissors then 750µL Digestion Media (Wash Media with 100µg/mL Liberase (Sigma, #5401127001) and 100µg/mL of DNAse I (Roche, #10104159001)) was added. Tissues were incubated at 37°C with end-over-end rotation, 30 minutes for RM and OM, 20 minutes for LNG. 13.3µL EDTA (0.5M) was added to each sample and then cells were washed with HBSS Media (HBSS (Ca, Mg Free, 500 mL), 10mM EDTA, 10mM HEPES, 2% FBS) and filtered through a 70µm nylon cell strainer. Cells were pelleted by centrifugation 500g for 10 minutes, resuspended with ACK (Ammonium-Chloride-Potassium) lysis buffer for 1 minute on ice, and then diluted with 9mL HBSS Media and centrifuged 500g for 5 minutes twice. Cells were then resuspended in 1mL Isolation Buffer (PBS, 0.1% BSA, 2mM EDTA) pre-mixed with 25µL anti-EpCAM-biotin-Dynabeads (anti-EpCAM-biotin antibody (G8.8, Biolegend) bound to Dynabeads Biotin Binder (ThermoFisher)) for a light epithelial cell depletion, incubated for 15 minutes on ice, washed with Isolation Buffer and placed on a Dynamag for 2 minutes. Supernatants were collected, centrifuged 500g for 5 minutes, resuspended in 100µL Staining Buffer (PBS, 1% BSA, 0.01% Tween) and 10µL Fc block, and incubated on ice for 10 minutes. Next, 0.5µg Biolegend TotalSeq Hashing antibodies B0301, B0302, or B0303 were added so that each mouse had all three nasal regions (RM, OM, and LNG) stained with one of the three antibodies, and incubated on ice for 20 minutes. Cells were then washed extensively to remove excess antibody with 10mL Staining Buffer and centrifugation at 500g for 5 minutes twice. Cells were resuspended in Loading Buffer (PBS and 0.04% BSA), counted, and pooled equally (13,500 cells/sample) between three mice for each region. Finally, each set of pooled cells were centrifuged 500g for 5 minutes and resuspended in 42µl Loading Buffer for downstream scRNA-seq processing.

### Single-cell RNA-seq

Pooled samples from each nasal region (RM, OM, and LNG) were processed using the 10x Genomics Chromium Next GEM Single Cell 3’ Kit v3.1 and Feature Barcoding Kit with dual indices per the manufacturer’s instructions. Approximately 40,000 cells per pooled reaction were loaded on the 10x Genomics Chromium Controller. Library quality was evaluated using the Agilent TapeStation 4200 (Agilent). Prior to sequencing, the gene expression and hashtag libraries were pooled 20:1. Sequencing was performed on either the NovaSeq 6000 or NextSeq 2000 (Illumina) with an average RNA read depth of 16,000 reads/cell and hashtag read depth of 500 reads/cell.

### Immunofluorescence Microscopy

Mice were euthanized in CO2 and their heads following skin, fur, and lower jaw removal were placed in 4% paraformaldehyde for 1-4 hours on ice for fixation. Heads were transferred to 0.5M EDTA for 2-3 days at 4°C for bone decalcification. Heads were transferred to 30% sucrose in PBS for cryoprotection for 2 days at 4°C then rapidly frozen in NEG-50 using dry ice. Frozen heads were stored at -20°C until cryostat sectioning. Mouse nasal tissues were cut into 50-100μm sections, permeabilized with 0.3% Tween in PBS (PBST) for 1 hour, then incubated overnight at 4°C with antibodies, DAPI, and Fc block at a 1:200 dilution in PBST. Antibodies used: anti-Influenza A virus NS1 (PA5-32243, ThermoFisher), anti-acetyl-α-tubulin (Ly640, D20G3, Cell Signaling Technology), anti-CD45 (30-F11, Biologend), anti-EpCAM (G8.8, Biolegend), anti-Krt13 (EPR3671, Abcam), anti-PD-L1 (10F.9G2, Biolegend), and anti-IgA (mA-6E1, ThermoFisher). Samples were then washed 3 times with PBST in 15-minute intervals at room temperature, mounted on glass slides with Prolong Gold, and visualized with an Olympus FLUOVIEW FV3000 confocal laser scanning microscope.

### qPCR

RM tissue was collected as described above from mice and lysed in Buffer RLT (Qiagen) + 1% beta-mercaptoethanol (Sigma) via gentleMACS Octo Dissociator in M-Tubes (Miltenyi Biotec). RNA was extracted from tissue lysate by RNAEasy Mini column purification (Qiagen) following the manufacturer’s instructions. cDNA was then generated following the SmartSeq II protocol as previously described (Trombetta et al., 2014). qPCR was performed using TaqMan reagents and probes (ThermoFisher) on a CFX384 Real-Time PCR System.

### Antibody-based depletion

Naïve or PR8 infected B6 mice were administered daily 20µg anti-CCR2 depleting antibodies (MC-21 generously provided by Prof. Matthias Mack, Universität Regensburg) or rat IgG2b,κ isotype control (Biolegend, #400644) intraperitoneally (i.p.). 24h following one administration, blood was collected from naïve mice by tail vein bleed into FACS buffer (PBS, 0.5% BSA, 2mM EDTA) and stored on ice before processing for flow cytometry.

PR8 infected mice were administered antibodies 3, 4, and 5 dpi in 24h intervals. For this experiment, mice were euthanized at 8 dpi.

### Flow cytometry

Blood was processed for flow cytometry by pelleting cells by centrifugation and resuspending with ACK lysis buffer to remove RBCs. Cells were then washed with FACS buffer and stained in 50 µL for flow cytometry using the following antibodies: anti-CCR2 (475301, R&D Systems), anti-CD11b (M1/70, Biolegend), anti-CD19 (6D5, Biolegend), anti-CD3e (145-2C11, Biolegend), anti-CD45 (30-F11, Biologend), anti-Ly6C (HK1.4, Biolegend), anti-Ly6G (1A8, Biolegend), and anti-NK1.1 (PK136, Biolegend). Cells were analyzed using the Beckman Coulter CytoFLEX.

For RM tissue, mice were anesthetized i.p. with ketamine (100 mg/kg body weight) and xylazine (10 mg/kg body weight) prior to euthanasia and administered 1µg anti-CD45 antibody (30-F11, Biologend) by retroorbital intravascular injection to label CD45+ cells in circulation. Mice were then euthanized 3 minutes later in CO2. RM tissue was processed as described above for tissue harvesting and single-cell suspension preparation through ACK lysis and dilution. Cells were then centrifuged and resuspended in 100 µL FACS buffer LIVE/DEAD Fixable Aqua Dead Cell Stain (ThermoFisher #L34966) per manufacturer’s instructions. Cells were then washed and stained for 30min at 4°C in the dark with Fc block diluted 1:200 and subsets of the following antibodies: anti-CD103 (2E7, Biolegend), anti-CD11b (M1/70, Biolegend), anti-CD11c (HL3, BD Biosciences), anti-CD19 (6D5, Biolegend), anti-CD3e (145-2C11, Biolegend), anti-CD4 (GK1.5, Biolegend), anti-CD45 (30-F11, Biologend), anti-CD64 (X54-5/7.1, Biolegend), anti-CD69 (H1.2F3, Biolegend), anti-CD8b (YTS156.7.7, Biolegend), anti-Ly6C (HK1.4, Biolegend), anti-Ly6G (1A8, Biolegend), anti-F4/80 (BM8, Biolegend), anti-MHC-II (M5/114.15.2, Biolegend), and anti-NK1.1 (PK136, Biolegend). To stain for Krt13, IgA, and IgK/L intracellularly, cells were fixed and permeabilized using the eBioscience Transcription Factor Staining Buffer Set (ThermoFisher #00-5523-00) according to the manufacturer’s instructions. Cells were stained with anti-Krt13 (EPR3671, abcam) at 1:200, or anti-IgA (mA-6E1, eBioscience), anti-IgK (187.1, BD Biosciences), and anti-IgL (R26-46, BD Biosciences) at 1:100. Following staining, cells were washed in FACS buffer, and analyzed. To determine cell counts, AccuCheck Counting Beads (ThermoFisher #PCB100) were added to every sample.

### RNAscope Microscopy

RNA *in situ* hybridization was performed according to manufacturer’s instructions for the RNAscope Multiplex Fluorescent Reagent Kit v2 (Advanced Cell Diagnostics ACD, 323270) on 20 µm thin sections of fixed-frozen murine nasal mucosa tissue collected 14 dpi. We implemented the following modifications to preserve tissue integrity: 1) 5 min PBS wash preceding initial baking of slides was removed; 2) slides were baked for 30 min at 60°C following EtOH dehydration; 3) target retrieval time was reduced to 5 min; 4) slides were baked for 60 min at 60°C following target retrieval; and 5) tissue sections were incubated in Protease Plus instead of Protease III for milder protease digestion. Probes used included Mm-Krt13 (ACD, 575341), Mm-Cxcr6-C2 (ACD, 871991-C2), Mm-Cxcl16-C3 (ACD, 466681-C3), and Mm-Cd274-C3 (ACD, 420501-C3). Following signal amplification, Opal 520 (Akoya Biosciences, FP1487001KT), Opal 570 (Akoya Biosciences, FP1488001KT), and Opal 690 (Akoya Biosciences, FP1497001KT) dyes were used, diluted 1:1000 in TSA buffer (ACD, 322809). Nuclei were stained with DAPI and slides were mounted with VECTASHIELD PLUS (Vector Laboratories, H-1900). Confocal images were collected using an Olympus FLUOVIEW FV3000 confocal laser scanning microscope.

### Quantification and statistical analysis

#### Single-cell RNA-seq alignment, cleanup, and pre-processing

To detect reads originating from IAV, we built a combined genome of mm10 (GRCm39) and the sequences for PR8 (NCBI Taxonomy ID #211044). The eight PR8 genomic viral segment sequences (NC_002023.1, NC_002022.1, NC_002021.1, NC_002020.1, NC_00219.1, NC_2018.1, NC_002017.1, and NC_002016.1) and associated IAV gene annotations were added to the GRCm39 FASTA and GTF files and processed using the CellRanger’s built in “mkref” function. Sequences were then aligned and quantified using this combined genome with the CellRanger toolkit (v6.0.1) via Cumulus tools (Li et al., 2020) (https://cumulus.readthedocs.io/en/stable/). Cell sample identity was assigned from the measurement of TotalSeqB aligned counts using the cumulus demultiplexing tool for feature barcoding, calling identity for any cell with at least 100 barcodes. To correct for transcript spill-over, cellbender (Fleming et al., 2022) was applied to the raw output UMI matrices from CellRanger with the following parameters: *expected_cells=30000*, *fpr=0.01, total_droplets_included=50000*. Cellbender corrected cells were then filtered based on Unique Molecular Identifiers (UMI) count (>750 & <10000), number of detected genes (>500), and percentage of mitochondrial genes (<15%). Finally, cells labeled as doublets by demultiplexing were removed.

#### Iterative clustering, cell cluster annotation, and IAV+ cell calling

Downstream analysis was performed using Seurat (v.4.2.1) (Hao et al., 2021). Briefly, the entire primary infection dataset underwent normalization using the *scTransform* function followed by principal component analysis (PCA), shared nearest neighbors (SNN) graph generation, Louvain clustering, and UMAP embedding. Clustering was performed at multiple resolutions to help annotate similar and dissimilar clusters. Using clustering *resolution = 0.6,* cluster specific/enriched markers were calculated. Each cluster was labeled by major cell type based on the expression of known lineage markers (e.g., *Omp*, *Epcam*, *Ptprc, Flt1,* etc.). Doublet clusters were also annotated based on the lack of unique markers and/or the presence of multiple mutually exclusive lineage markers (e.g., *Omp+Ptprc+* cells). Following annotation, doublet clusters were removed, and the normalization/clustering/doublet removal process was repeated twice more (total of three times) until no doublet clusters were discernable.

The dataset was then divided into separate objects by cell type label for further subclustering. Following the same routine applied to the full dataset, clusters for each cell type were annotated with subset/state labels based on prior knowledge and previously published scRNAseq datasets of the nasal mucosa (Brann et al., 2020; Ualiyeva et al., 2024; Ziegler et al., 2021). After the first set of annotations in every cell type object, it was apparent that there were still intra-sample doublets present: mostly contaminating cell types, but also within cell type doublets (e.g., ionocyte/sustentacular doublets). These clusters were iteratively removed like in the analysis of the full dataset, for a total of three rounds in each cell type, yielding a total of 127 clusters across the whole dataset. To visualize these clusters’ relationships and distribution across nasal regions, we built a cell cluster “phylogenetic tree” using ARBOL (Zheng et al., 2021), where the first tier encodes major cell type, the second tier encodes defined subtypes, and the third tier encodes cluster identity (**Figure S1F**).

We note that neurons, mostly olfactory sensory neurons, make up a large number of cells in our primary infection dataset (n > 50,000). Given their importance in mouse olfaction, and their broad distribution through the OE and OM (**Figure 1B**), we believe that their relative abundance in our scRNA-seq data is concordant with the anatomy and biology of the nasal mucosa.

Since IAV transcript capture was sparse, we classified IAV+ cells as any cell with 2 or more UMIs aligned to any IAV PR8 gene.

#### Compositional analyses

After removing multiplets, immune (>97.5%) and endothelial cells (89%) had nearly all cells assigned a sample replicate while neurons (30.7%), epithelial cells (65.9%), fibroblasts (52.4%), and other stromal cells (56.4%) had lower sample annotation rates (**Figure S1E**). We note that cells without a sample replicate assignment were excluded from all compositional analyses. Within cell type frequencies were calculated on a per replicate basis by counting the number of cells within each cluster label and dividing by the total number of cells for that cell type captured in that replicate. For tissue- and region-level compositional analyses, cell cluster abundances were calculated by deriving cell cluster frequencies over all labeled cells in each sample replicate, scaling to 3,000 cells per replicate, and log transforming. Subsequent PCAs were calculated using the center-log-ratio (clr) transformed data.

Proportionality analysis was performed using the propr package (v4.2.6) (Quinn et al., 2017). We compared the proportionality statistic **ρ,** calculated from the clr transformed abundance data, to standard Pearson correlation across all pairwise comparisons and found **ρ** to be more stringent for significance cutoffs (FDR<0.05) generated by permutating testing (**Figure S8A**). We built a network using Cytoscape (v3.9.1) comprised of all significantly proportional cell cluster pairs to assess groups of cell clusters with similar cell abundance trajectories throughout the infection time course.

RM sample replicate distances were calculated using the Aitchison distance (Aitchison et al., 2000). Euclidean distance on the clr transformed data then calculated between all pairs of RM sample replicates, yielding three distances within a timepoint and nine distances between two timepoints. Statistics on Aitchison distances were performed using a one-sided non-parametric Welch’s ANOVA and Dunnett’s T3 test for multiple comparisons in Prism.

#### Neutrophil pseudotime analysis

The neutrophil pseudotime analysis was performed using diffusion mapping as previously described (Grieshaber-Bouyer et al., 2021). A principal component analysis was run on all cells assigned to granulocyte clusters, excluding mast cells. The first 20 principal components were used to compute a cell-to-cell distance matrix using *1 – Pearson correlation coefficient* as the distance metric. Using the destiny package in R (Angerer et al., 2016), we computed a diffusion map with standard parameters with density normalization and rotate enabled. We manually selected “Progenitor” cells as the root of the trajectory and used the DPT function to calculate the pseudotime values, manually scaling the values from 0 to 1.

#### Cell-cell signaling analysis

Three cell networks (IFN-stimulated MDMs : *Gzmk+* CD8 T cells : *Ifng+Cd200+* CD4 T cells; *Cd103*+ DCs : *Dusp2+Icam1+* mature neutrophils : *Gp2+Lyz2+* goblet/secretory cells*; Krt13+Il1a+* epithelial cells : *Cd103+* CD8 T cells : CD4 T cells) were selected based on high proportionality throughout the primary infection time-course or specific biological interest to the authors. Data for the clusters of interest in each network were then subset to RM and timepoint of interest to best capture individual cells with sufficient spatial and temporal proximity to plausibly interact, and re-normalized with Seurat’s *NormalizeData* function. Since NICHES calculates the multiplicative expression of ligand-receptor pairs from a random sampling of cells from each cell type to predict cell-cell communication, Adaptively thresholded Low-Rank Approximation (ALRA) imputation was applied to each cell network to reduce the impact of technical zeros due to potential dropout events (Linderman et al., 2022). NICHES was then run on each cell network individually, drawing from the OmniPath database of ligand-receptor pairs (Türei et al., 2016) to generate a cell interaction object whereby rows are ligand-receptor pairs and columns are cell type pairs (Raredon et al., 2023). These objects were then scaled and passed through principal component analysis and UMAP dimensionality reduction to generate low-dimensional embeddings of cell interactions. Differentially expressed interactions were identified using the Seurat *FindAllMarkers* function, and highly differentially expressed interactions validated by literature review were selected for display as heatmaps.

#### RNAscope co-localization quantification

RNAscope quantification was carried out using ImageJ software on five fields of view (FOV) from both the naïve and 14 dpi timepoints (n = 5/timepoint). Ten slices were selected in the z-plane from each FOV to generate maximum intensity projections across 5 µm, and color channels were separated using the “Split Channels” function. For each image, a threshold was set for signal intensity in the *Krt13* channel, allowing generation of a mask of *Krt13* expression that was subsequently overlaid on both the DAPI and *Cxcl16* channels to generate regions of interest for further analysis. Within the region of interest in the DAPI channel, nuclei were manually counted with the aid of the ImageJ plugin “Cell Counter,” with inclusion of most segmented nuclei. Within the region of interest in the *Cxcl16* channel, a threshold was set for *Cxcl16* expression with the aid of a signal intensity histogram and then punctuate dots were counted automatically with the “Analyze Particles” function. A measure of average *Cxcl16* puncta per DAPI-stained nuclei in the *Krt13+* region was calculated for each FOV.

#### Cluster annotation in memory samples

To assign cluster labels to new scRNA-seq datasets generated from the nasal mucosa, we leveraged the structure of our data to test the label transfer methods provided in Seurat (Hao et al., 2021) and scANVI (Xu et al., 2021). We separated the cells from one RM replicate from each time point as a query dataset, using the remaining cells as the reference. With Seurat, we implemented the FindTransferAnchors function on the scTransformed data, using the first 40 PCs from the PCA as the reference. Labels were then assigned with the TransferData function using either cell cluster or cell type identities. With scANVI, we first built a scVI model on the reference data, and then a scANVI model using either cell cluster or cell type identities as labels. The reference scANVI model was then used to train a model on the query data. We calculated the percentage of correctly called cell labels using each method for both sets of labels and found calling to be superior on the cell type level. Thus, we next repeated each procedure within each cell type to learn cell cluster labels and found Seurat to perform better across all cell types (**Figure S9C**). This approach was repeated using two other non-overlapping sets of sample replicates (one per time point) to assess reproducibility. To further validate, we calculated cell cluster abundances using the predicted cell cluster labels for the query replicates and projected into the PCA calculated across the RM samples.

We next applied the two-step label transfer approach using Seurat to new data generated from RM 60 dpi and 2 and 5 dprc. Before performing label transfer, we removed all hashtag annotated cell doublets from the new dataset and applied the same filtering criteria as above. We next performed *scTransform* and PCA on the new dataset. We then predicted cell type labels using all cells from RM samples in the primary infection dataset for reference. We next removed any cells from subsequent analysis that had a maximum prediction assignment score < 0.8 (i.e., 80% is the greatest confidence in label prediction), making up 5% of the total dataset. Given our loading strategy and number of intrasample doublets found in the primary infection dataset, we chose a more stringent cutoff following cell type label prediction. Separating into each cell type and using the processed data from the matching cell type in the primary infection dataset as reference, we performed the same procedure. Here, we were more liberal, keeping all cells with a maximum prediction assignment score ≥ 0.4 since very similar clusters within cell types could receive almost equal prediction probability (e.g., Resting Basal and Abi3bp Resting Basal). With cell cluster labels assigned, we then calculated cell cluster abundances as above and performed downstream differential expression analysis.

## SUPPLEMENTARY MATERIAL

**Figure S1:**
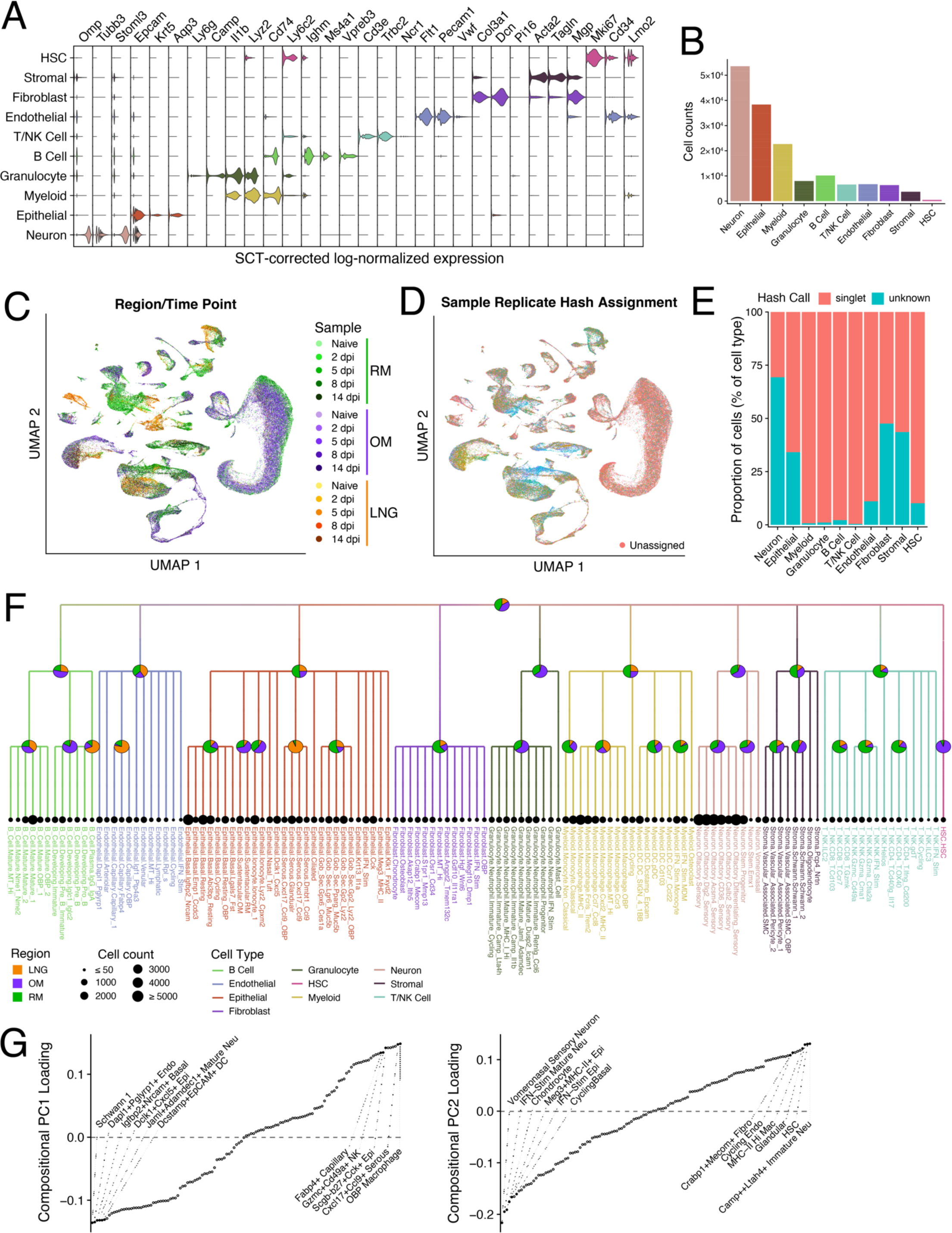
Clustering and sample replicate assignment across nasal mucosa regions and cell types. (A) Violin plots of representative genes used to assign cell type identity to clusters. (B) Numbers of cells classified as each cell type across all samples, including cells that did not receive a hash call. (C) UMAP embedding as in Figure 2A colored by region and time point. (D) UMAP as in (D) colored by hash call assignment. There are 45 sample replicates across the dataset in addition to cells without definitive hash identities (“Unassigned”). (E) Stacked bar chart depicting the relative proportion of cells with assigned sample replicate identity (i.e., hash call) by cell type. Singlet = single sample replicate call; unknown = too few barcodes measured to assign a sample replicate identity. (F) Cell lineage tree generated with ARBOL (Zheng et al., 2023) depicting all 127 clusters found in the dataset through cell type subclustering. Branches are colored by cell type. Pie charts at each branching point depict the relative proportion of cells from each nasal mucosa region. Dot size at each end node is proportional to the number of cells assigned to that cluster. See **Supplementary Table 1** for all differentially expressed markers across clusters within each cell type. (G) Cell cluster abundance loadings from the PCA shown in Figure 2C for PC1 (left) and PC2 (right) from (F). Cell cluster names for several of the most negative and most positive weights for each PC are depicted.

**Figure S2:**
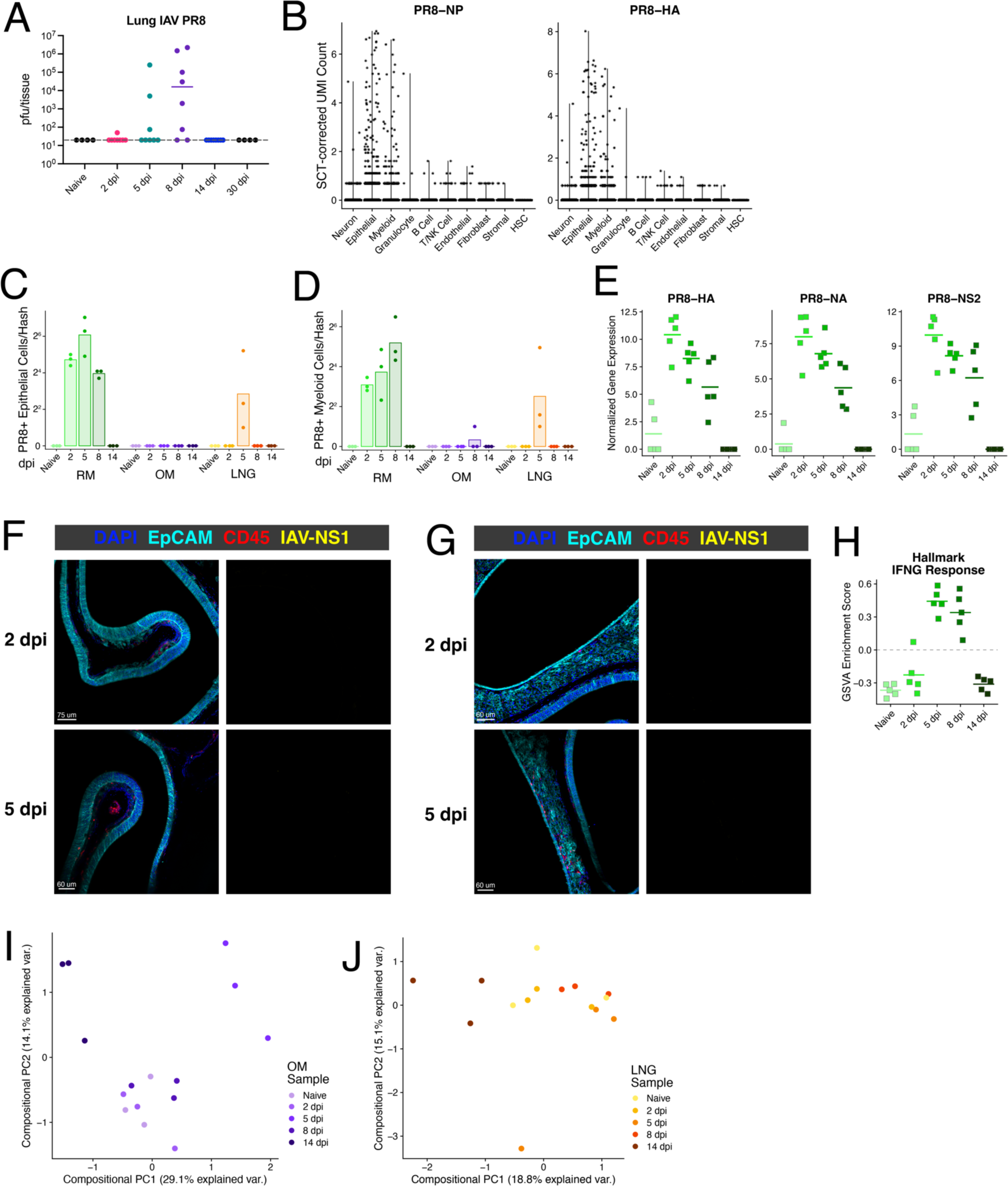
Viral transcript capture, global antiviral responses, and changes in OM and LNG composition. (A) Infectious IAV PR8 quantification in plaque forming units (pfu) of the entire lung. (B) scTransform-corrected UMI counts for the IAV PR8 genes encoding NP (left) and HA (right) by cell type. (C & D) Number of PR8+ epithelial cells (D) and myeloid cells (E) by time point and region. PR8+ cells are classified by having at least 2 UMI aligning to PR8 genes. (E) Log-normalized expression of PR8 genes in bulk RNA-seq samples generated from whole RM tissue lysate (n = 5/timepoint). (F & G) Representative images of IAV infection in OM (E) and LNG (F) taken from mice 2 dpi (top) and 5 dpi (bottom). Staining for EpCAM (teal), CD45 (red), and IAV-NS1 (yellow). Images on the right depict only the signal in the IAV-NS1 channel. (H) Gene Set Variation Analysis (GSVA) Enrichment score for Hallmark Response to Interferon-Gamma on total RM tissue lysate bulk RNA-seq data (n = 5/time point) (I & J) Compositional PCA of only OM samples (I) and only LNG samples (J).

**Figure S3:**
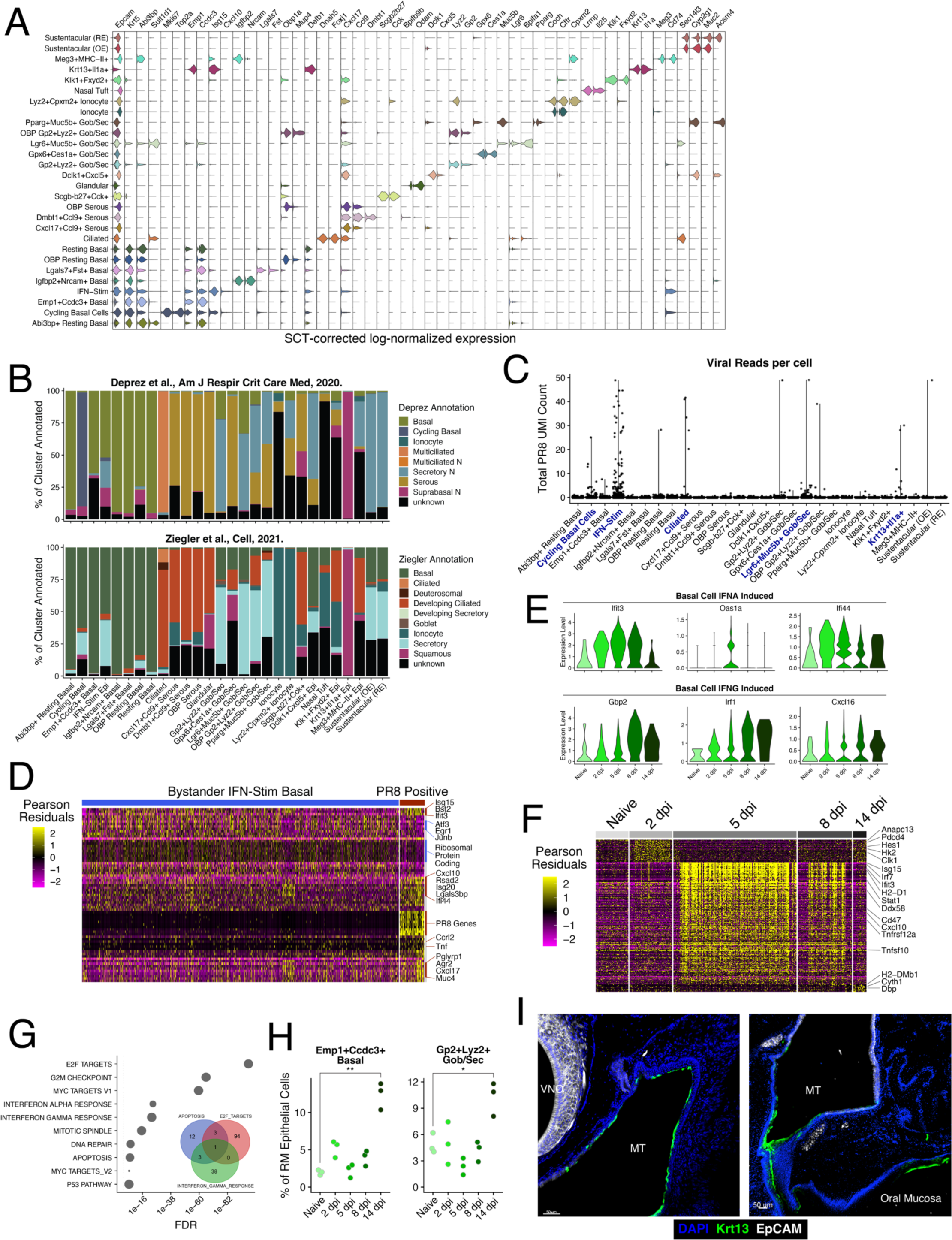
Epithelial cell heterogeneity and response cluster dynamics. (A) Violin plots depicting differentially expressed marker genes (FDR<0.01) across all 28 epithelial clusters (see **Supplementary Table 1**). (B) Stacked bar charts depicting the relative proportions of mouse nasal mucosa epithelial cells annotated for each human nasal epithelial cell type from (Deprez et al., 2020) and (Ziegler et al., 2021) by label transfer. Cells with poor assignments (maximum prediction score < 0.4) were labeled as “unknown”. (C) Summative scTransform-corrected UMI counts across all 8 IAV genes by epithelial cell cluster. Clusters with ≥5 cells with more than 2 PR8 UMIs have their cluster names bolded in blue. (D) Heatmap depicting all differentially expressed genes between PR8 positive (≥2 PR8 UMIs) and bystander IFN-Stim epithelial cells from RM 5 and 8 dpi. Scaled Pearson residuals from scTransform are plotted. (E) Expression in IFN-Stim epithelial cells of representative IFNα and IFNγ induced ISGs from the stimulation signatures derived from airway basal cell cultures (Ziegler et al., 2020). (F) Heatmap depicting all differentially expressed genes (FDR<0.01) in Cycling Basal cells from RM between timepoints. Scaled Pearson residuals from scTransform are plotted. (G) Gene set analysis (hypergeometric test) of all differentially enriched genes in Cycling Basal cells compared to all other epithelial cell clusters (FDR<0.01). The Hallmark pathways from MsigDB (v7.5.1) were used. Inset: venn diagram showing the number genes that are within the Hallmark Apoptosis, E2F Targets, and Interferon Gamma Response pathways. (H) Relative frequencies of *Emp1*+*Ccdc3*+ basal cells (top) and *Gp2*+*Lyz2*+ Gob/Sec cells (bottom) as a proportion of all epithelial cells per replicate RM sample. Only cells with assigned hash calls are included. Welch’s t test, *p < 0.05, **p < 0.01. (I) Representative immunofluorescence images of the nasal mucosa 14 dpi taken more posterior than Figure 3K where the nasal mucosa connects to the oral cavity.

**Figure S4:**
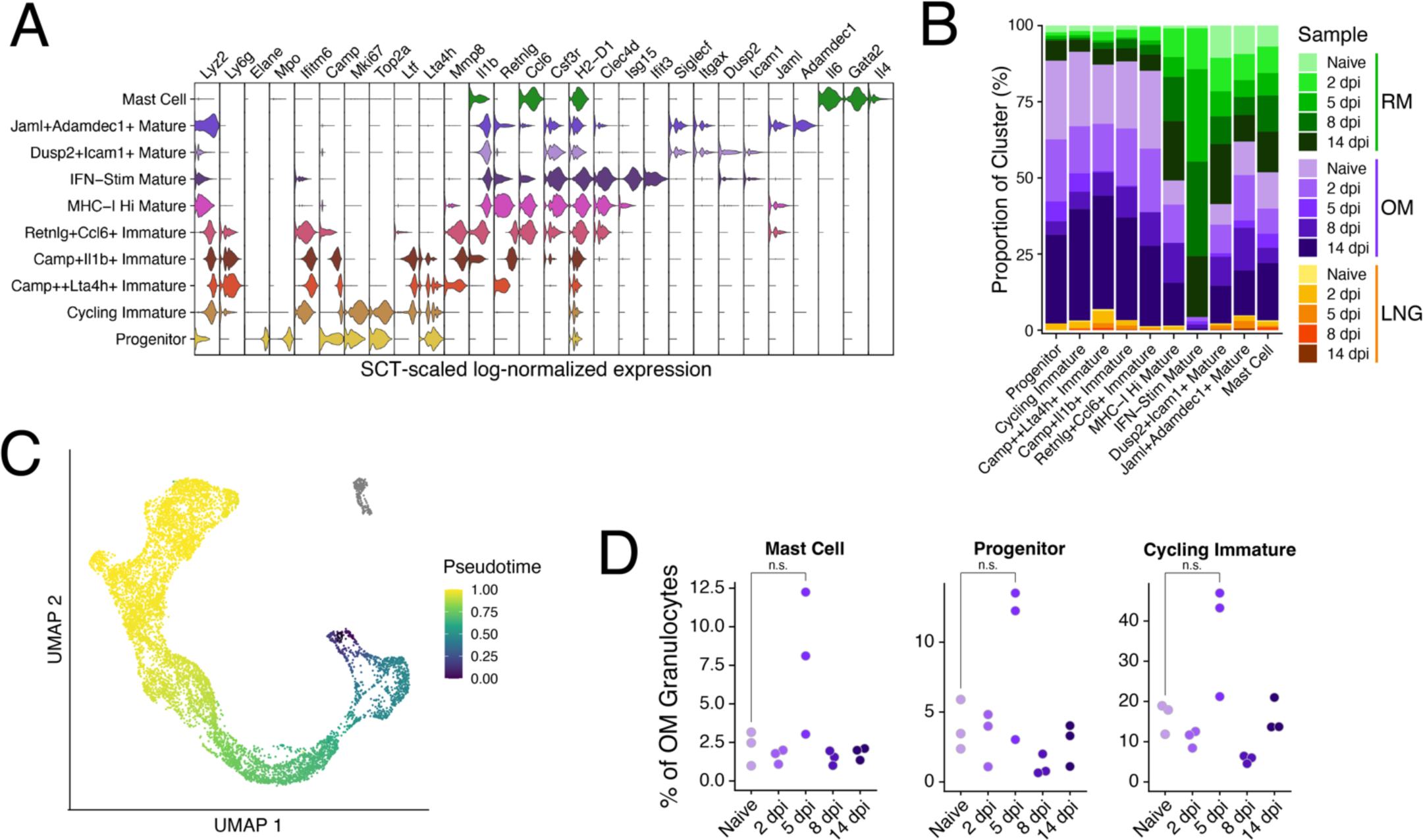
Granulocyte heterogeneity. (A) Violin plots depicting differentially expressed marker genes (FDR<0.01) across all 10 granulocyte clusters (see **Supplementary Table 1**). (B) Stacked bar chart depicting the relative proportions of cells annotated for each granulocyte cluster by region and time point. (C) UMAP of granulocytes colored by pseudotime. Mast Cells were not included in the pseudotime analysis and are colored gray. (D) Relative frequencies of Mast cells (left), progenitors (center), and cycling immature (right) as a proportion of all granulocytes per replicate OM sample.

**Figure S5:**
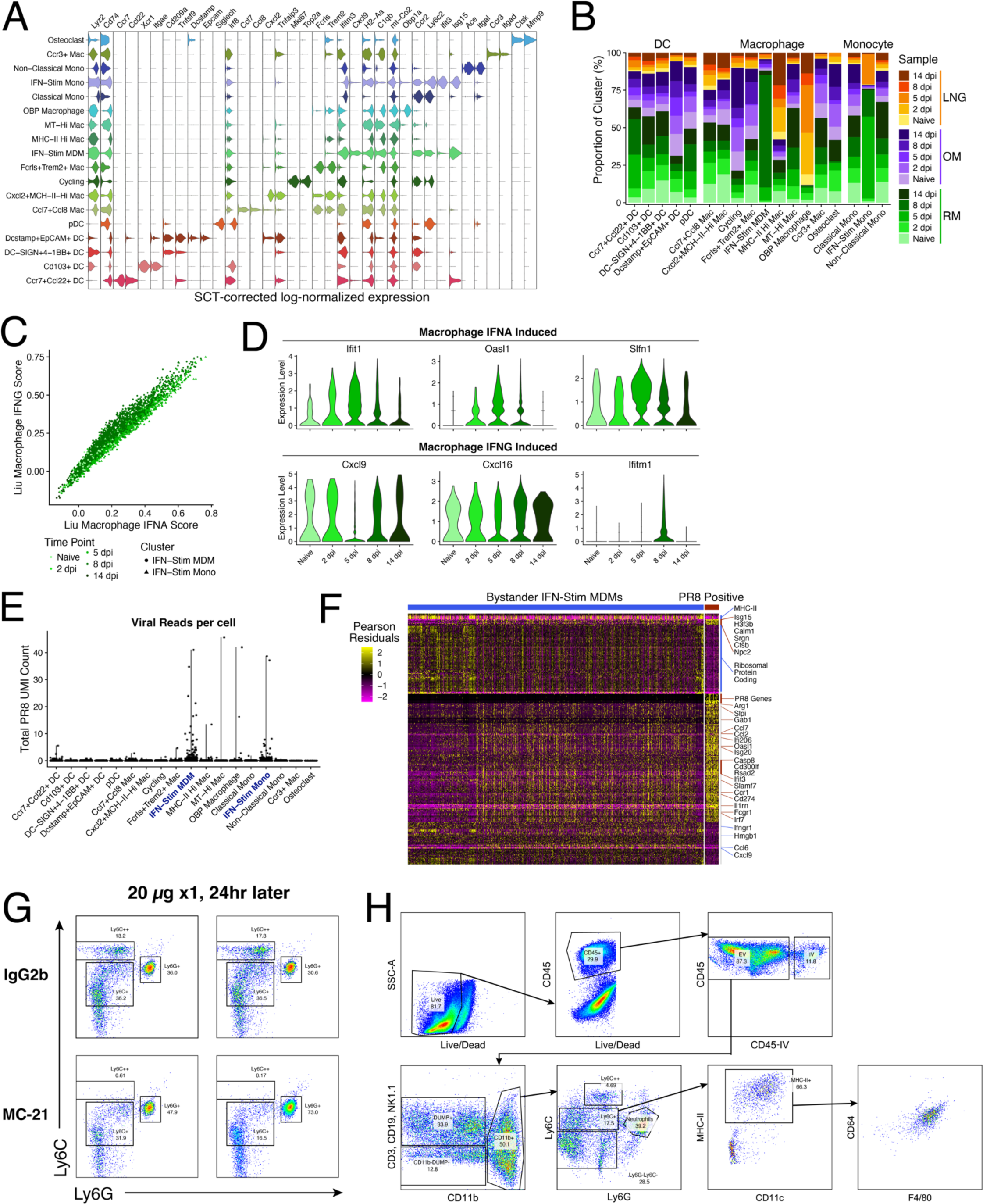
Myeloid heterogeneity, viral+ cells, and monocyte depletion. (A) Violin plots depicting differentially expressed marker genes (FDR<0.01) across all 18 macrophage, monocyte, and DC clusters (see **Supplementary Table 1**). (B) Stacked bar chart depicting the relative proportions of cells annotated for each myeloid cluster by region and time point. (C) IFN-Stim monocyte and MDM scores for signatures derived from bone marrow-derived macrophage cultures stimulated with IFNα or IFNγ (Liu et al., 2012). (D) Expression in representative IFNα and IFNγ induced ISGs from (C). (E) Summative scTransform-corrected UMI counts across all 8 IAV genes by myeloid cell cluster. Clusters with ≥5 cells with more than 2 PR8 UMIs have their cluster names bolded in blue. (F) Heatmap depicting all differentially expressed genes between PR8 positive (≥2 PR8 UMIs) and bystander IFN-Stim MDMs from RM 8 dpi. Scaled Pearson residuals from scTransform are plotted. (G) Mice (n = 4) were treated i.p. with control antibody (top) or anti-CCR2 antibody (bottom) and blood was collected 24 hours later for flow cytometry. Pre-gated on Dead–CD45+CD3– CD19–CD11b+. (H) Representative gating scheme for Ly6C+ and Ly6C++ monocytes in the nasal mucosa.

**Figure S6:**
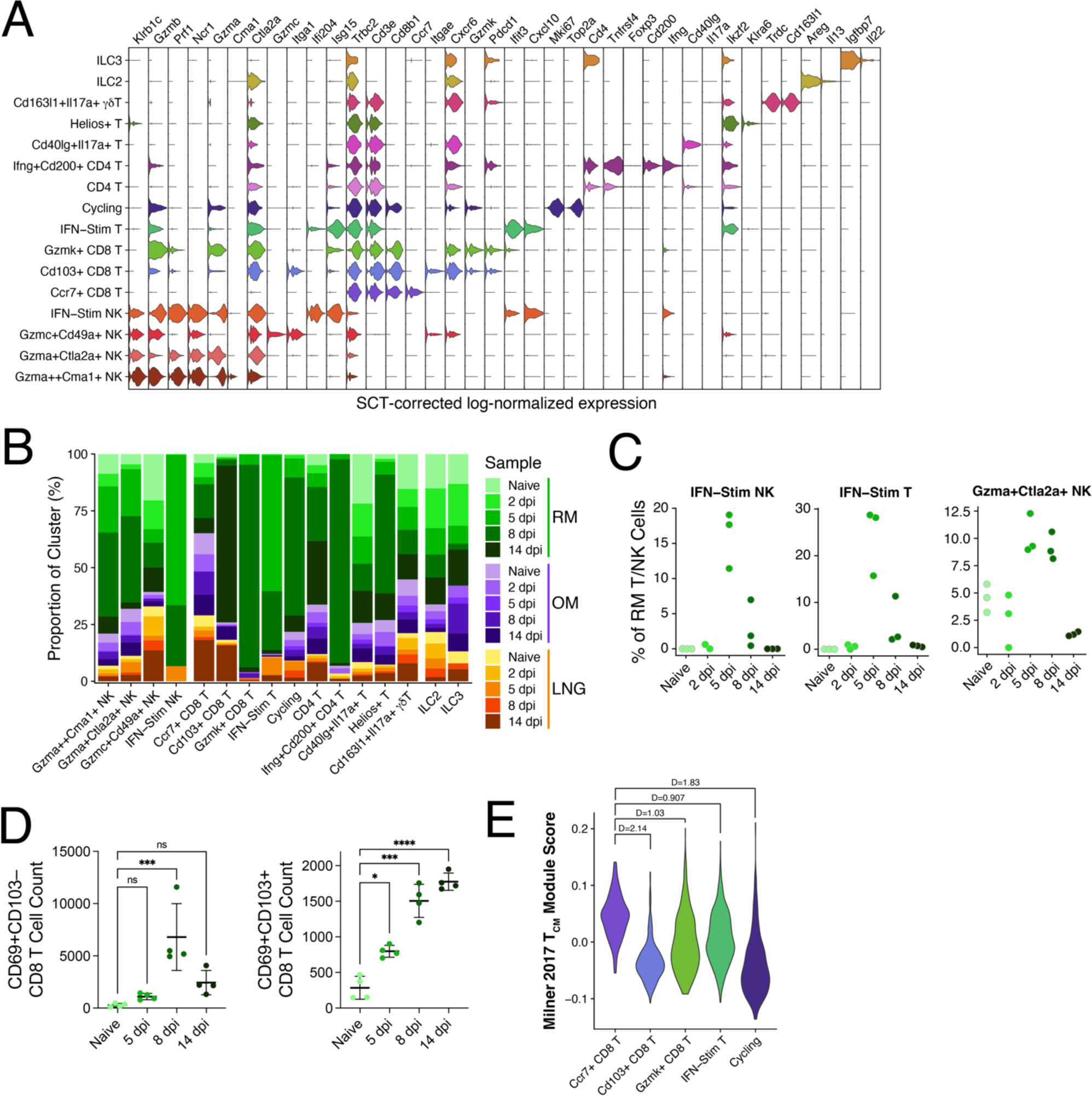
T cell, NK cell, and innate lymphocyte heterogeneity and T_RM_ responses. (A) Violin plots depicting differentially expressed marker genes (FDR<0.01) across all 16 T cell, NK cell, and innate lymphocyte cell clusters (see **Supplementary Table 1**). (B) Stacked bar chart depicting the relative proportions of cells annotated for each myeloid cluster by region and time point. (C) Relative frequencies of IFN-Stim NK cells (left), IFN-Stim T cells (middle), and *Gzma*+*Ctla2a*+ NK cells (right) as a proportion of all T cells, NK cells, and innate lymphocytes per RM replicate sample. (D) Mice were infected with 10^4^ PFU IAV PR8 and RM tissue was collected to stain for T cells. Kruskal-Wallis, *p < 0.05, **p < 0.01. (E) Violin plot depicting a gene module score derived from the universal T circulating memory (T_CM_) cell signature as published in (Milner et al., 2017) across all CD8 T cell clusters. Cohen’s D for effect size is reported between *Ccr7*+ CD8 T cells and each other cluster.

**Figure S7:**
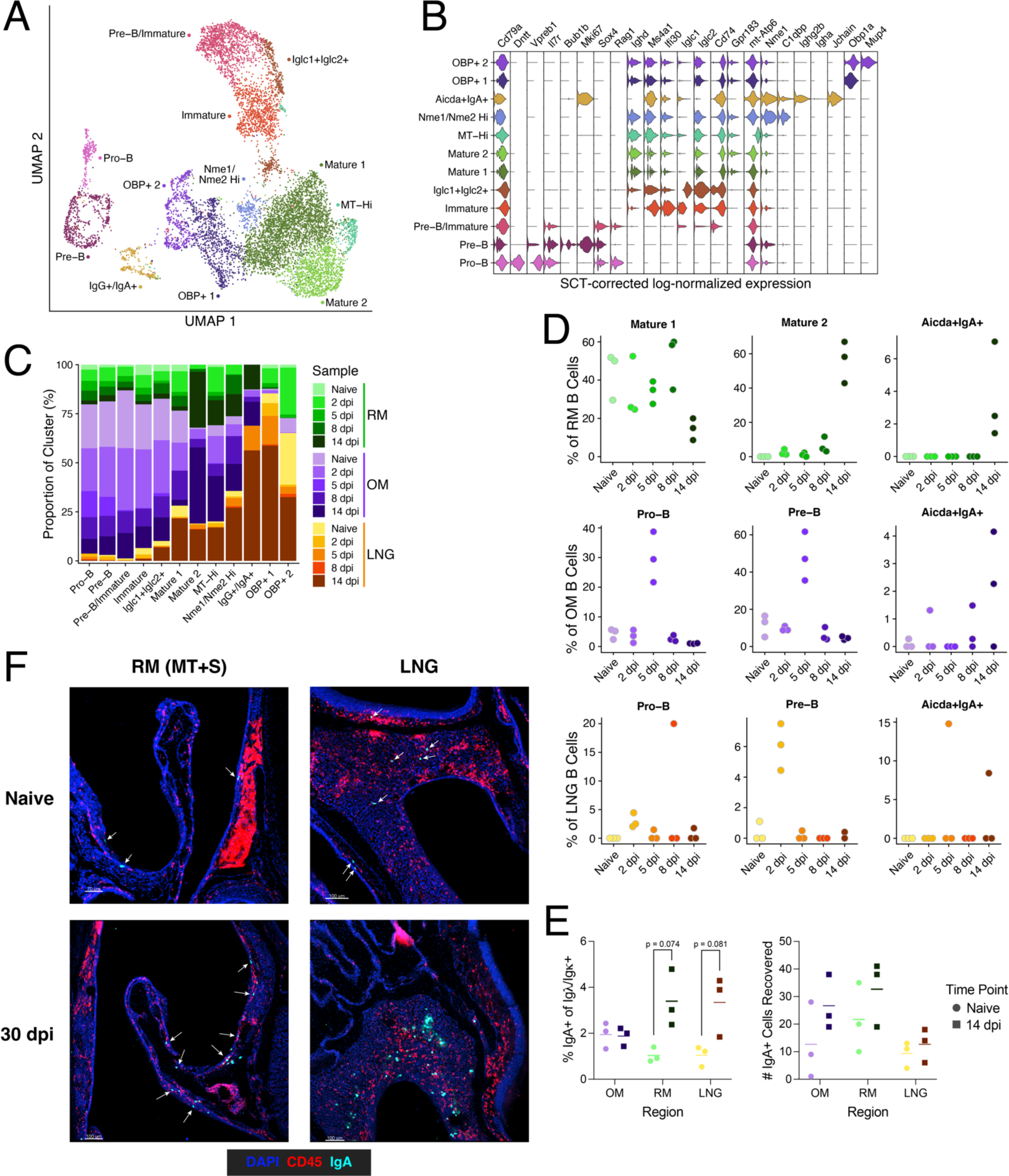
B cell heterogeneity and cluster dynamics. (A) UMAP embedding of 10,167 B cells across 12 clusters. (B) Violin plots depicting differentially expressed marker genes (FDR<0.01) across all 12 B cell clusters (see **Supplementary Table 1**). (C) Stacked bar chart depicting the relative proportions of cells annotated for each B cell cluster by region and time point. (D) Relative frequencies of several B cell clusters as proportions of all B cells per RM replicate sample (top), OM replicate sample (middle), and LNG replicate sample (bottom). (E) Mice were infected with 10^4^ PFU IAV PR8 and RM, OM, and LNG tissue were collected to stain intracellularly for IgA cells. Welch’s t test. (F) Representative immunofluorescence images staining for IgA producing cells in the RM (left) and LNG (right) in naïve mice (top) and 30 dpi (bottom). White arrows point to IgA+ cells in the sparser images.

**Figure S8:**
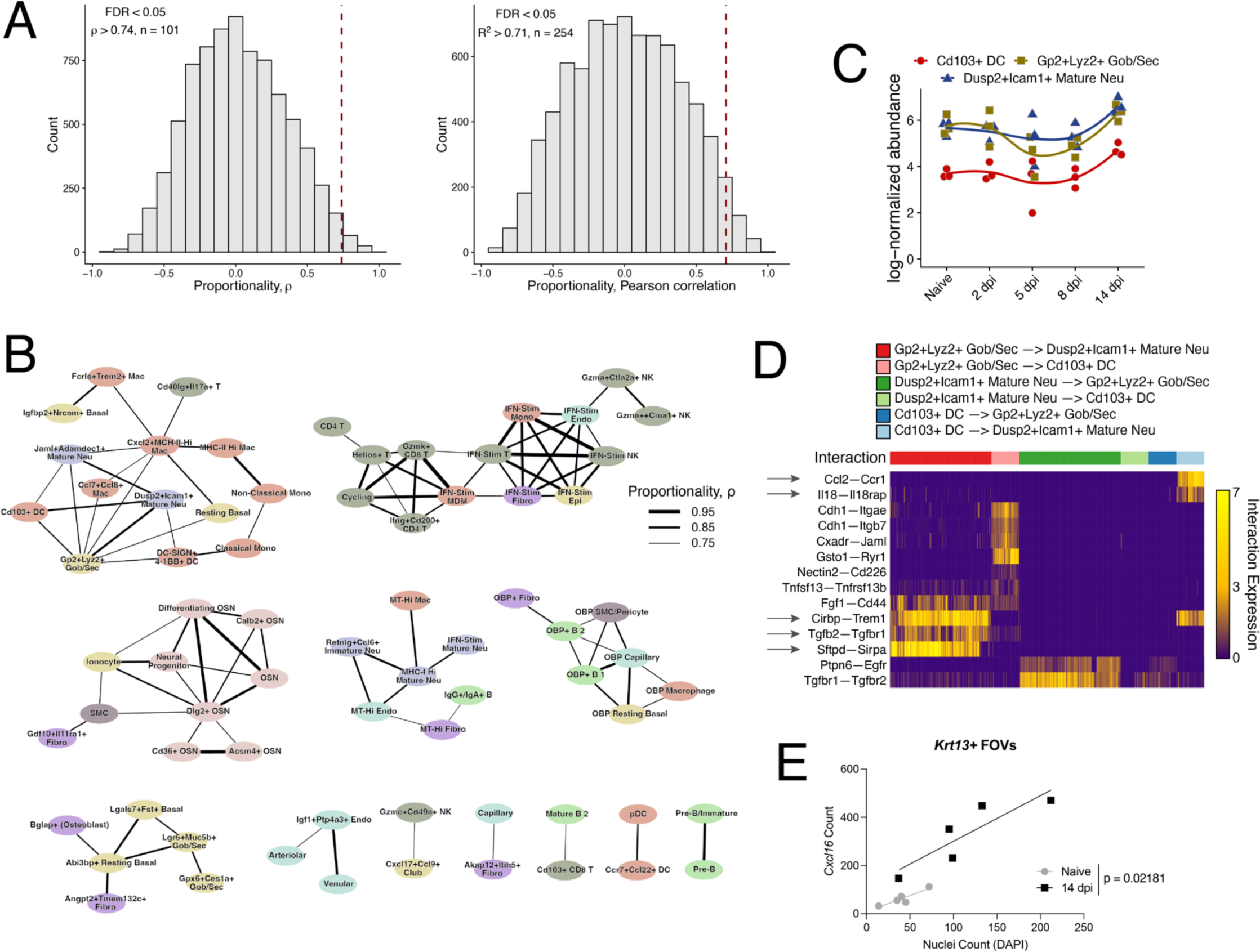
Proportionality and cell-cell communication analysis. (A) Histograms of calculated proportionality (left) and Pearson correlation (right) statistics across all RM sample replicates. The significance cutoff (FDR<0.05) for each statistic is marked by the red dashed line and was calculated from a background of 1000 permutations of the data. (B) Network of all significantly proportional (FDR<0.01) cell clusters across all RM replicate samples. Nodes are colored by cell type and edge weight is representative of proportionality. (C) Abundance plot of *Cd103*+ DCs, *Gp2*+*Lyz2*+ Gob/Sec cells, and *Dusp2*+*Icam1*+ Mature Neutrophils in replicate RM samples. Smoothed lines are calculated using local polynomial regression fitting. (D) Heatmap depicting a subset of differentially expressed receptor-ligand interaction pairs between single-cell pairs identified by NICHES (Raredon et al., 2023) for the clusters depicted in (C); see **Supplementary Table 2**. Interaction expression is the multiplicative expression of receptor and ligand gene expression for each member of a single-cell pair. Arrows highlight interactions described in the text. (E) Quantification of nuclei and *Cxcl16* spots within *Krt13*+ regions of interest across 10 RNAscope images in a naïve mouse and 14 dpi (n = 5/timepoint). Linear regression, p value reported for difference in intercepts.

**Figure S9:**
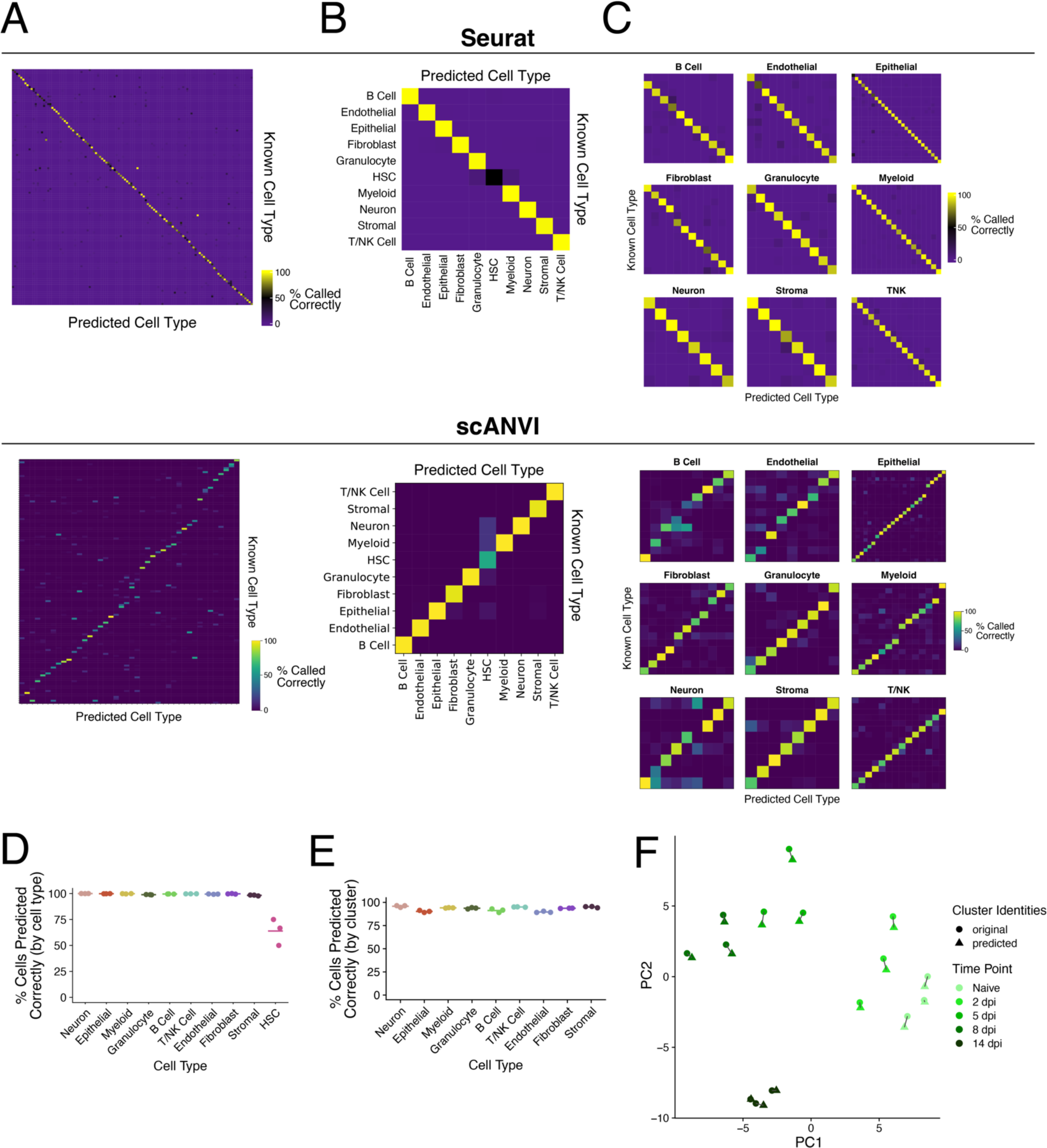
Validating label transfer methods to assign cell cluster identities to new nasal mucosa samples. Initially, one sample replicate per timepoint was separated from the RM dataset to be used as a query trained using a reference made from the remaining sample replicates and unclassified cells. The top and bottom depict results from Seurat and scANVI respectively. (A) Heatmaps depicting the per-cluster on-target prediction frequency when calculated across all 127 cluster labels. (B) Heatmaps depicting the per-cell-type on-target prediction frequency when calculated across the 9 cell type labels. (C) After predicting cell type labels, new query and reference pairs were generated within each cell type and label transfer was performed within each. Heatmaps depicting the per-cluster on-target prediction frequency when calculated within all clusters within each respective cell type. Following comparison of Seurat and scANVI, the label transfer approach was reiterated two more times in Seurat using other sets of replicates as the query and reference datasets. (D) Percentage of cells with an accurate cell type label by cell type. Each dot reflects a distinct query+reference comparison. (E) Percentage of cells with an accurate cluster label by cell type. Each dot reflects a distinct query+reference comparison. (F) Compositional PCA from Figure 2I where the query sample replicates were projected using the predicted cell cluster labels. Lines connect the query predicted sample compositions to their matching compositions as determined by the initial clustering and labeling.

**Figure S10:**
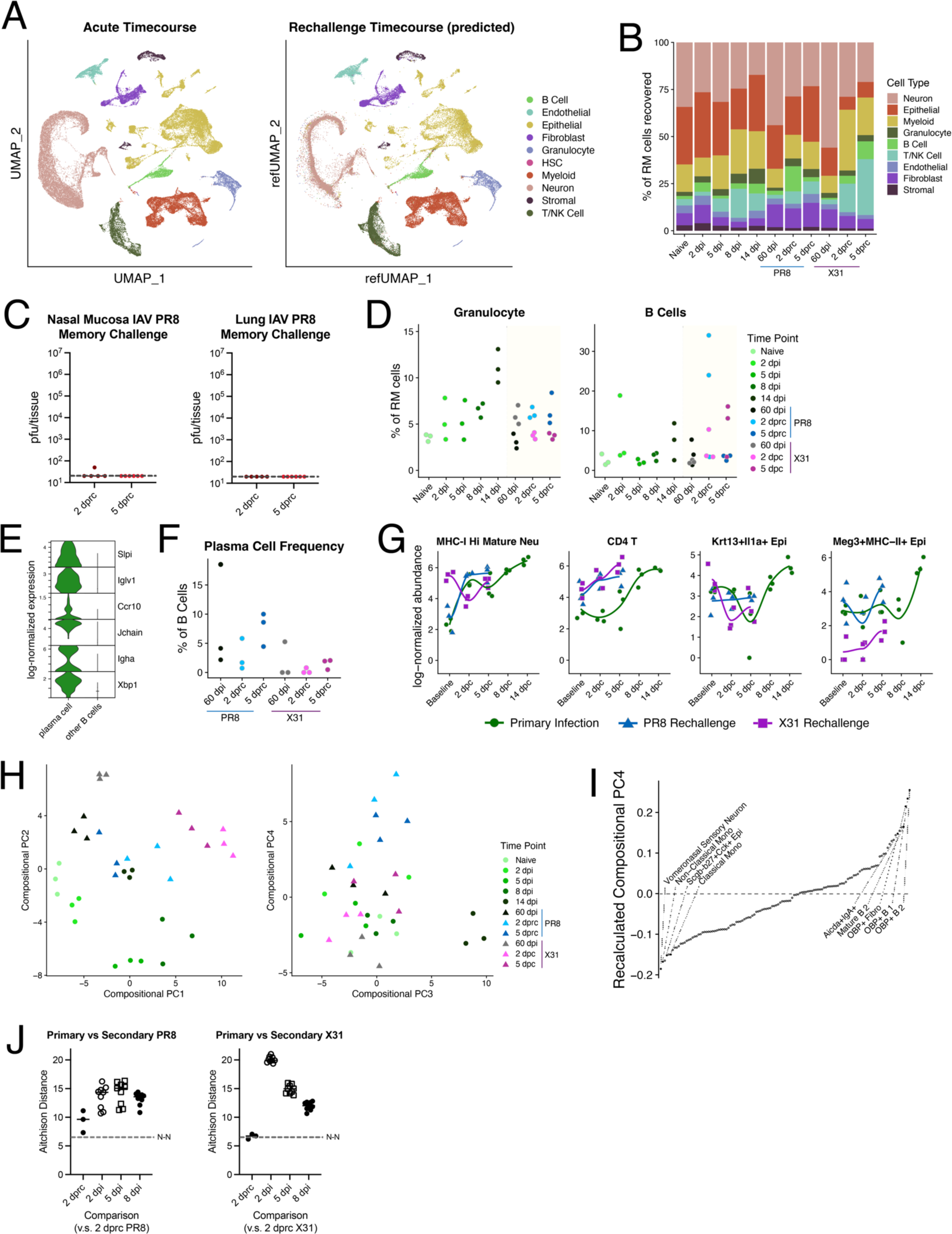
Changes in RM composition following secondary challenge. (A) UMAP of all RM cells from the primary infection dataset (left) and the projected UMAP of all cells in the rechallenge dataset (right) colored by cell type. (B) Stacked bar chart depicting the relative proportions of cells annotated for each cell type by time point. Cells from RM samples only. (C) Infectious IAV PR8 quantification in pfu of the entire nasal mucosa (left) and lung (right) during IAV PR8 rechallenge. (D) Relative frequencies of granulocytes and B cells as a proportion of all sequenced cells per RM replicate sample in primary infection and following rechallenge. (E) Violin plot of select plasma cell specific/enriched genes (FDR corrected p-values ≤ 10^–118^ by 1-vs-rest Wilcoxon Rank Sum Test). (F) Relative frequency of the plasma cell cluster as proportions of all B cells per RM replicate in rechallenge. Plasma cells were not found in the primary infection dataset. (G) Abundance plots of various clusters showing overlaid primary infection (green) and rechallenge responses with PR8 in blue and X31 in pink. Baseline refers to samples from naïve mice in primary infection and to samples from 60 dpi in rechallenge. dpc = days post challenge. Smoothed lines are calculated using local polynomial regression fitting. (H) Compositional PCA recalculated using both primary infection and secondary challenge RM sample replicates. PCs 1-2 (left) and 3-4 (right). (I) Cell cluster abundance loadings for PC3 (E). Cell cluster names for several of the most negative and most positive weights for each PC are depicted. (J) Heatmap depicting all pairwise Aitchison distances between all RM sample replicates.

**Supplementary Table 1: Differentially expressed genes across clusters within each cell type.** Differential expression analysis was performed across cells from all samples within each cell type using the Wilcoxon Rank Sum test.

**Supplementary Table 2: Cell cluster counts per sample replicate across all regions and time points.** Annotated counts for each sample replicate across all 127 clusters in the dataset. Abundances per sample, and the total number of cells recovered per reaction are also reported.

**Supplementary Table 3: Differentially expressed receptor-ligand pairs identified by NICHES.** Within each NICHES analysis, differential expression was performed across all cell-pairs at the specific timepoint of interest using the ROC test built into Seurat. See Figure 6.

**Supplementary Table 4: Differentially expressed genes between primary infection and secondary challenge.** Differential expression analysis (“bidomal” test) was performed between timepoints using cells from RM only within the specified clusters in Figure 7E.

## REFERENCES

Adamson, M. S., Nesic, S., Buness, A., Bayrak, K., Schmitz, S., Soler, S., Zillinger, T., Marx, S., Lambing, S., Andryka-Cegielski, K., et al. (2022). RIG-I activation primes and trains innate antiviral immune memory. 2022.10.27.514004.

Aegerter, H., Kulikauskaite, J., Crotta, S., Patel, H., Kelly, G., Hessel, E. M., Mack, M., Beinke, S. and Wack, A. (2020). Influenza-induced monocyte-derived alveolar macrophages confer prolonged antibacterial protection. Nat Immunol 21, 145–157.

Aitchison, J., Barceló-Vidal, C., Martín-Fernández, J. A. and Pawlowsky-Glahn, V. (2000). Logratio Analysis and Compositional Distance. Mathematical Geology 32, 271–275.

Angerer, P., Haghverdi, L., Büttner, M., Theis, F. J., Marr, C. and Buettner, F. (2016). destiny: diffusion maps for large-scale single-cell data in R. Bioinformatics 32, 1241–1243.

Ariotti, S., Hogenbirk, M. A., Dijkgraaf, F. E., Visser, L. L., Hoekstra, M. E., Song, J.-Y., Jacobs, H., Haanen, J. B. and Schumacher, T. N. (2014). Skin-resident memory CD8+ T cells trigger a state of tissue-wide pathogen alert. Science 346, 101–105.

Bastard, P., Rosen, L. B., Zhang, Q., Michailidis, E., Hoffmann, H.-H., Zhang, Y., Dorgham, K., Philippot, Q., Rosain, J., Béziat, V., et al. (2020). Autoantibodies against type I IFNs in patients with life-threatening COVID-19. Science 370,.

Bastard, P., Zhang, Q., Zhang, S.-Y., Jouanguy, E. and Casanova, J.-L. (2022). Type I interferons and SARS-CoV-2: from cells to organisms. Current Opinion in Immunology 74, 172–182.

Bosch, A. A. T. M., Biesbroek, G., Trzcinski, K., Sanders, E. A. M. and Bogaert, D. (2013). Viral and Bacterial Interactions in the Upper Respiratory Tract. PLOS Pathogens 9, e1003057.

Bosch-Camós, L., Alonso, U., Esteve-Codina, A., Chang, C.-Y., Martín-Mur, B., Accensi, F., Muñoz, M., Navas, M. J., Dabad, M., Vidal, E., et al. (2022). Cross-protection against African swine fever virus upon intranasal vaccination is associated with an adaptive-innate immune crosstalk. PLOS Pathogens 18, e1010931.

Boudreau, C. M. and Alter, G. (2019). Extra-Neutralizing FcR-Mediated Antibody Functions for a Universal Influenza Vaccine. Frontiers in Immunology 10,.

Bouvier, N. M. and Lowen, A. C. (2010). Animal Models for Influenza Virus Pathogenesis and Transmission. Viruses 2, 1530–1563.

Boyd, D. F., Allen, E. K., Randolph, A. G., Guo, X. J., Weng, Y., Sanders, C. J., Bajracharya, R., Lee, N. K., Guy, C. S., Vogel, P., et al. (2020). Exuberant fibroblast activity compromises lung function via ADAMTS4. Nature 587, 466–471.

Brandes, M., Klauschen, F., Kuchen, S. and Germain, R. N. (2013). A Systems Analysis Identifies a Feedforward Inflammatory Circuit Leading to Lethal Influenza Infection. Cell 154, 197–212.

Brann, D. H., Tsukahara, T., Weinreb, C., Lipovsek, M., Van den Berge, K., Gong, B., Chance, R., Macaulay, I. C., Chou, H.-J., Fletcher, R. B., et al. (2020). Non-neuronal expression of SARS-CoV-2 entry genes in the olfactory system suggests mechanisms underlying COVID-19-associated anosmia. Science Advances 6, eabc5801.

Broggi, A., Ghosh, S., Sposito, B., Spreafico, R., Balzarini, F., Cascio, A. L., Clementi, N., Santis, M. D., Mancini, N., Granucci, F., et al. (2020). Type III interferons disrupt the lung epithelial barrier upon viral recognition. Science 369, 706–712.

Büttner, M., Ostner, J., Müller, C. L., Theis, F. J. and Schubert, B. (2021). scCODA is a Bayesian model for compositional single-cell data analysis. Nat Commun 12, 6876.

Cao, Y., Lin, Y., Ormerod, J. T., Yang, P., Yang, J. Y. H. and Lo, K. K. (2019). scDC: single cell differential composition analysis. BMC Bioinformatics 20, 721.

Chapman, T. J. and Topham, D. J. (2010). Identification of a Unique Population of Tissue-Memory CD4+ T Cells in the Airways after Influenza Infection That Is Dependent on the Integrin VLA-1. The Journal of Immunology 184, 3841–3849.

Chen, X., Liu, S., Goraya, M. U., Maarouf, M., Huang, S. and Chen, J.-L. (2018). Host Immune Response to Influenza A Virus Infection. Front Immunol 9, 320.

Chen, K., Zhang, J., Huang, Y., Tian, X., Yang, Y. and Dong, A. (2022). Single-cell RNA-seq transcriptomic landscape of human and mouse islets and pathological alterations of diabetes. iScience 25, 105366.

Clark, S. E. (2020). Commensal bacteria in the upper respiratory tract regulate susceptibility to infection. Current Opinion in Immunology 66, 42–49.

Clark, R. A., Chong, B., Mirchandani, N., Brinster, N. K., Yamanaka, K., Dowgiert, R. K. and Kupper, T. S. (2006). The Vast Majority of CLA+ T Cells Are Resident in Normal Skin1. The Journal of Immunology 176, 4431–4439.

Costa-Martins, A. G., Mane, K., Lindsey, B. B., Ogava, R. L. T., Castro, Í., Jagne, Y. J., Sallah, H. J., Armitage, E. P., Jarju, S., Ahadzie, B., et al. (2021). Prior upregulation of interferon pathways in the nasopharynx impacts viral shedding following live attenuated influenza vaccine challenge in children. Cell Reports Medicine 2, 100465.

Crowl, J. T., Heeg, M., Ferry, A., Milner, J. J., Omilusik, K. D., Toma, C., He, Z., Chang, J. T. and Goldrath, A. W. (2022). Tissue-resident memory CD8+ T cells possess unique transcriptional, epigenetic and functional adaptations to different tissue environments. Nat Immunol 23, 1121–1131.

Dann, E., Henderson, N. C., Teichmann, S. A., Morgan, M. D. and Marioni, J. C. (2022). Differential abundance testing on single-cell data using k-nearest neighbor graphs. Nat Biotechnol 40, 245–253.

Darrah, P. A., Zeppa, J. J., Maiello, P., Hackney, J. A., Wadsworth, M. H., Hughes, T. K., Pokkali, S., Swanson, P. A., Grant, N. L., Rodgers, M. A., et al. (2020). Prevention of tuberculosis in macaques after intravenous BCG immunization. Nature 577, 95–102.

Davis, J. D. and Wypych, T. P. (2021). Cellular and functional heterogeneity of the airway epithelium. Mucosal Immunol 14, 978–990.

Denning, N.-L., Aziz, M., Murao, A., Gurien, S. D., Ochani, M., Prince, J. M. and Wang, P. (2020). Extracellular CIRP as an endogenous TREM-1 ligand to fuel inflammation in sepsis. JCI Insight 5,.

Deprez, M., Zaragosi, L.-E., Truchi, M., Becavin, C., Ruiz García, S., Arguel, M.-J., Plaisant, M., Magnone, V., Lebrigand, K., Abelanet, S., et al. (2020). A Single-Cell Atlas of the Human Healthy Airways. Am J Respir Crit Care Med 202, 1636–1645.

Dumm, R. E., Wellford, S. A., Moseman, E. A. and Heaton, N. S. (2020). Heterogeneity of Antiviral Responses in the Upper Respiratory Tract Mediates Differential Non-lytic Clearance of Influenza Viruses. Cell Reports 32,.

Fleming, S. J., Chaffin, M. D., Arduini, A., Akkad, A.-D., Banks, E., Marioni, J. C., Philippakis, A. A., Ellinor, P. T. and Babadi, M. (2022). Unsupervised removal of systematic background noise from droplet-based single-cell experiments using CellBender. 791699.

Gardai, S. J., Xiao, Y.-Q., Dickinson, M., Nick, J. A., Voelker, D. R., Greene, K. E. and Henson, P. M. (2003). By Binding SIRPα or Calreticulin/CD91, Lung Collectins Act as Dual Function Surveillance Molecules to Suppress or Enhance Inflammation. Cell 115, 13–23.

Gloor, G. B., Macklaim, J. M., Pawlowsky-Glahn, V. and Egozcue, J. J. (2017). Microbiome Datasets Are Compositional: And This Is Not Optional. Frontiers in Microbiology 8,.

Grieshaber-Bouyer, R., Radtke, F. A., Cunin, P., Stifano, G., Levescot, A., Vijaykumar, B., Nelson-Maney, N., Blaustein, R. B., Monach, P. A. and Nigrovic, P. A. (2021). The neutrotime transcriptional signature defines a single continuum of neutrophils across biological compartments. Nat Commun 12, 2856.

Habibi, M. S., Thwaites, R. S., Chang, M., Jozwik, A., Paras, A., Kirsebom, F., Varese, A., Owen, A., Cuthbertson, L., James, P., et al. (2020). Neutrophilic inflammation in the respiratory mucosa predisposes to RSV infection. Science 370,.

Hao, Y., Hao, S., Andersen-Nissen, E., Mauck, W. M., Zheng, S., Butler, A., Lee, M. J., Wilk, A. J., Darby, C., Zager, M., et al. (2021). Integrated analysis of multimodal single-cell data. Cell 184, 3573–3587.e29.

Harkema, J. R., Carey, S. A. and Wagner, J. G. (2006). The Nose Revisited: A Brief Review of the Comparative Structure, Function, and Toxicologic Pathology of the Nasal Epithelium. Toxicol Pathol 34, 252–269.

Heim, T. A., Lin, Z., Steele, M. M., Mudianto, T. and Lund, A. W. (2023). CXCR6 promotes dermal CD8+ T cell survival and transition to long-term tissue residence. 2023.02.14.528487.

Hu, M.-W., Kim, D. W., Liu, S., Zack, D. J., Blackshaw, S. and Qian, J. (2019). PanoView: An iterative clustering method for single-cell RNA sequencing data. PLOS Computational Biology 15, e1007040.

Hufford, M. M., Kim, T. S., Sun, J. and Braciale, T. J. (2015). The Effector T Cell Response to Influenza Infection. In Influenza Pathogenesis and Control - Volume II (ed. Oldstone, M. B. A.) and Compans, R. W.), pp. 423–455. Cham: Springer International Publishing.

Ibricevic, A., Pekosz, A., Walter, M. J., Newby, C., Battaile, J. T., Brown, E. G., Holtzman, M. J. and Brody, S. L. (2006). Influenza Virus Receptor Specificity and Cell Tropism in Mouse and Human Airway Epithelial Cells. Journal of Virology 80, 7469–7480.

Iijima, N. and Iwasaki, A. (2014). A local macrophage chemokine network sustains protective tissue-resident memory CD4 T cells. Science 346, 93–98.

Iuliano, A. D., Roguski, K. M., Chang, H. H., Muscatello, D. J., Palekar, R., Tempia, S., Cohen, C., Gran, J. M., Schanzer, D., Cowling, B. J., et al. (2018). Estimates of global seasonal influenza-associated respiratory mortality: a modelling study. Lancet 391, 1285–1300.

Iwasaki, A. and Medzhitov, R. (2015). Control of adaptive immunity by the innate immune system. Nat. Immunol. 16, 343–353.

Jiang, X., Clark, R. A., Liu, L., Wagers, A. J., Fuhlbrigge, R. C. and Kupper, T. S. (2012). Skin infection generates non-migratory memory CD8+ TRM cells providing global skin immunity. Nature 483, 227–231.

Johansson, C. and Kirsebom, F. C. M. (2021). Neutrophils in respiratory viral infections. Mucosal Immunol 14, 815–827.

Johnson, P. R., Feldman, S., Thompson, J. M., Mahoney, J. D. and Wright, P. F. (1986). Immunity to Influenza A Virus Infection in Young Children: A Comparison of Natural Infection, Live Cold-Adapted Vaccine, and Inactivated Vaccine. The Journal of Infectious Diseases 154, 121–127.

Johnson Jr., P. R., Feldman, S., Thompson, J. M., Mahoney, J. D. and Wright, P. F. (1985). Comparison of long-term systemic and secretory antibody responses in children given live, attenuated, or inactivated influenza A vaccine. Journal of Medical Virology 17, 325– 335.

Jung, S.-H., Hwang, B.-H., Shin, S., Park, E.-H., Park, S.-H., Kim, C. W., Kim, E., Choo, E., Choi, I. J., Swirski, F. K., et al. (2022). Spatiotemporal dynamics of macrophage heterogeneity and a potential function of Trem2hi macrophages in infarcted hearts. Nat Commun 13, 4580.

Kadoki, M., Patil, A., Thaiss, C. C., Brooks, D. J., Pandey, S., Deep, D., Alvarez, D., Andrian, U. H. von, Wagers, A. J., Nakai, K., et al. (2017). Organism-Level Analysis of Vaccination Reveals Networks of Protection across Tissues. Cell 171, 398–413.e21.

Kaufmann, E., Sanz, J., Dunn, J. L., Khan, N., Mendonça, L. E., Pacis, A., Tzelepis, F., Pernet, E., Dumaine, A., Grenier, J.-C., et al. (2018). BCG Educates Hematopoietic Stem Cells to Generate Protective Innate Immunity against Tuberculosis. Cell 172, 176–190.e19.

Kim, Y.-M. and Shin, E.-C. (2021). Type I and III interferon responses in SARS-CoV-2 infection. Exp Mol Med 53, 750–760.

Klinkhammer, J., Schnepf, D., Ye, L., Schwaderlapp, M., Gad, H. H., Hartmann, R., Garcin, D., Mahlakõiv, T. and Staeheli, P. (2018). IFN-λ prevents influenza virus spread from the upper airways to the lungs and limits virus transmission. eLife 7, e33354.

Kochs, G., García-Sastre, A. and Martínez-Sobrido, L. (2007). Multiple Anti-Interferon Actions of the Influenza A Virus NS1 Protein. J Virol 81, 7011–7021.

Krammer, F. (2019). The human antibody response to influenza A virus infection and vaccination. Nat Rev Immunol 19, 383–397.

Lavelle, E. C. and Ward, R. W. (2022). Mucosal vaccines — fortifying the frontiers. Nat Rev Immunol 22, 236–250.

Li, B., Gould, J., Yang, Y., Sarkizova, S., Tabaka, M., Ashenberg, O., Rosen, Y., Slyper, M., Kowalczyk, M. S., Villani, A.-C., et al. (2020). Cumulus provides cloud-based data analysis for large-scale single-cell and single-nucleus RNA-seq. Nat Methods 17, 793– 798.

Li, H., Wang, X., Wang, Y., Zhang, M., Hong, F., Wang, H., Cui, A., Zhao, J., Ji, W. and Chen, Y.-G. (2022). Cross-species single-cell transcriptomic analysis reveals divergence of cell composition and functions in mammalian ileum epithelium. Cell Regeneration 11, 19.

Li, J., Hubisz, M. J., Earlie, E. M., Duran, M. A., Hong, C., Varela, A. A., Lettera, E., Deyell, M., Tavora, B., Havel, J. J., et al. (2023). Non-cell-autonomous cancer progression from chromosomal instability. Nature 620, 1080–1088.

Liew, F., Talwar, S., Cross, A., Willett, B. J., Scott, S., Logan, N., Siggins, M. K., Swieboda, D., Sidhu, J. K., Efstathiou, C., et al. (2023). SARS-CoV-2-specific nasal IgA wanes 9 months after hospitalisation with COVID-19 and is not induced by subsequent vaccination. eBioMedicine 87,.

Lin, H. and Peddada, S. D. (2020). Analysis of microbial compositions: a review of normalization and differential abundance analysis. npj Biofilms Microbiomes 6, 1–13.

Linderman, G. C., Zhao, J., Roulis, M., Bielecki, P., Flavell, R. A., Nadler, B. and Kluger, Y. (2022). Zero-preserving imputation of single-cell RNA-seq data. Nat Commun 13, 192.

Liu, S.-Y., Sanchez, D. J., Aliyari, R., Lu, S. and Cheng, G. (2012). Systematic identification of type I and type II interferon-induced antiviral factors. Proceedings of the National Academy of Sciences 109, 4239–4244.

Lovell, D., Pawlowsky-Glahn, V., Egozcue, J. J., Marguerat, S. and Bähler, J. (2015). Proportionality: A Valid Alternative to Correlation for Relative Data. PLOS Computational Biology 11, e1004075.

Mack, M., Cihak, J., Simonis, C., Luckow, B., Proudfoot, A. E. I., Plachý, J., Brühl, H., Frink, M., Anders, H.-J., Vielhauer, V., et al. (2001). Expression and Characterization of the Chemokine Receptors CCR2 and CCR5 in Mice1. The Journal of Immunology 166, 4697–4704.

MacLean, A. J., Richmond, N., Koneva, L., Attar, M., Medina, C. A. P., Thornton, E. E., Gomes, A. C., El-Turabi, A., Bachmann, M. F., Rijal, P., et al. (2022). Secondary influenza challenge triggers resident memory B cell migration and rapid relocation to boost antibody secretion at infected sites. Immunity 55, 718–733.e8.

Major, J., Crotta, S., Llorian, M., McCabe, T. M., Gad, H. H., Priestnall, S. L., Hartmann, R. and Wack, A. (2020). Type I and III interferons disrupt lung epithelial repair during recovery from viral infection. Science 369, 712–717.

Manicassamy, B., Manicassamy, S., Belicha-Villanueva, A., Pisanelli, G., Pulendran, B. and García-Sastre, A. (2010). Analysis of in vivo dynamics of influenza virus infection in mice using a GFP reporter virus. PNAS 107, 11531–11536.

Mao, T., Israelow, B., Peña-Hernández, M. A., Suberi, A., Zhou, L., Luyten, S., Reschke, M., Dong, H., Homer, R. J., Saltzman, W. M., et al. (2022). Unadjuvanted intranasal spike vaccine elicits protective mucosal immunity against sarbecoviruses. Science 378, eabo2523.

Matsuoka, Y., Lamirande, E. W. and Subbarao, K. (2009). The mouse model for influenza. Curr Protoc Microbiol Chapter 15, Unit 15G.3.

McMaster, S. R., Wilson, J. J., Wang, H. and Kohlmeier, J. E. (2015). Airway-Resident Memory CD8 T Cells Provide Antigen-Specific Protection against Respiratory Virus Challenge through Rapid IFN-γ Production. The Journal of Immunology 195, 203–209.

Milner, J. J., Toma, C., Yu, B., Zhang, K., Omilusik, K., Phan, A. T., Wang, D., Getzler, A. J., Nguyen, T., Crotty, S., et al. (2017). Runx3 programs CD8+ T cell residency in non-lymphoid tissues and tumours. Nature 552, 253–257.

Montoro, D. T., Haber, A. L., Biton, M., Vinarsky, V., Lin, B., Birket, S. E., Yuan, F., Chen, S., Leung, H. M., Villoria, J., et al. (2018). A revised airway epithelial hierarchy includes CFTR-expressing ionocytes. Nature 560, 319–324.

Morens, D. M., Taubenberger, J. K. and Fauci, A. S. (2023). Rethinking next-generation vaccines for coronaviruses, influenzaviruses, and other respiratory viruses. Cell Host & Microbe 31, 146–157.

Morgan, A. J., Guillen, C., Symon, F. A., Birring, S. S., Campbell, J. J. and Wardlaw, A. J. (2008). CXCR6 identifies a putative population of retained human lung T cells characterised by co-expression of activation markers. Immunobiology 213, 599–608.

Moriyama, M., Chen, I.-Y., Kawaguchi, A., Koshiba, T., Nagata, K., Takeyama, H., Hasegawa, H. and Ichinohe, T. (2016). The RNA- and TRIM25-Binding Domains of Influenza Virus NS1 Protein Are Essential for Suppression of NLRP3 Inflammasome-Mediated Interleukin-1β Secretion. Journal of Virology 90, 4105–4114.

Ols, S., Yang, L., Thompson, E. A., Pushparaj, P., Tran, K., Liang, F., Lin, A., Eriksson, B., Karlsson Hedestam, G. B., Wyatt, R. T., et al. (2020). Route of Vaccine Administration Alters Antigen Trafficking but Not Innate or Adaptive Immunity. Cell Reports 30, 3964–3971.e7.

Onodera, T., Takahashi, Y., Yokoi, Y., Ato, M., Kodama, Y., Hachimura, S., Kurosaki, T. and Kobayashi, K. (2012). Memory B cells in the lung participate in protective humoral immune responses to pulmonary influenza virus reinfection. Proceedings of the National Academy of Sciences 109, 2485–2490.

Ordovas-Montanes, J., Dwyer, D. F., Nyquist, S. K., Buchheit, K. M., Vukovic, M., Deb, C., Wadsworth, M. H., Hughes, T. K., Kazer, S. W., Yoshimoto, E., et al. (2018). Allergic inflammatory memory in human respiratory epithelial progenitor cells. Nature 560, 649.

Ordovas-Montanes, J., Beyaz, S., Rakoff-Nahoum, S. and Shalek, A. K. (2020). Distribution and storage of inflammatory memory in barrier tissues. Nat Rev Immunol 1–13.

Park, A. and Iwasaki, A. (2020). Type I and Type III Interferons – Induction, Signaling, Evasion, and Application to Combat COVID-19. *Cell Host Microbe* 27, 870–878.

Patterson-Cross, R. B., Levine, A. J. and Menon, V. (2021). Selecting single cell clustering parameter values using subsampling-based robustness metrics. BMC Bioinformatics 22, 39.

Pizzolla, A., Nguyen, T. H. O., Smith, J. M., Brooks, A. G., Kedzierska, K., Heath, W. R., Reading, P. C. and Wakim, L. M. (2017). Resident memory CD8+ T cells in the upper respiratory tract prevent pulmonary influenza virus infection. Science Immunology 2,.

Quinn, T. P., Richardson, M. F., Lovell, D. and Crowley, T. M. (2017). propr: An R-package for Identifying Proportionally Abundant Features Using Compositional Data Analysis. Sci Rep 7, 16252.

Quinn, T. P., Erb, I., Richardson, M. F. and Crowley, T. M. (2018). Understanding sequencing data as compositions: an outlook and review. Bioinformatics 34, 2870–2878.

Raredon, M. S. B., Yang, J., Kothapalli, N., Lewis, W., Kaminski, N., Niklason, L. E. and Kluger, Y. (2023). Comprehensive visualization of cell–cell interactions in single-cell and spatial transcriptomics with NICHES. Bioinformatics 39, btac775.

Ratnasiri, K., Wilk, A. J., Lee, M. J., Khatri, P. and Blish, C. A. (2023). Single-cell RNA-seq methods to interrogate virus-host interactions. Semin Immunopathol 45, 71–89.

Rossen, R. D., Butler, W. T., Waldman, R. H., Alford, R. H., Hornick, R. B., Togo, Y. and Kasel, J. A. (1970). The Proteins in Nasal Secretion: II. A Longitudinal Study of IgA and Neutralizing Antibody Levels in Nasal Washings From Men Infected With Influenza Virus. JAMA 211, 1157–1161.

Roth, G. A., Abate, D., Abate, K. H., Abay, S. M., Abbafati, C., Abbasi, N., Abbastabar, H., Abd-Allah, F., Abdela, J., Abdelalim, A., et al. (2018). Global, regional, and national age-sex-specific mortality for 282 causes of death in 195 countries and territories, 1980–2017: a systematic analysis for the Global Burden of Disease Study 2017. *The Lancet* 392, 1736–1788.

Russell, M. W., Moldoveanu, Z., Ogra, P. L. and Mestecky, J. (2020). Mucosal Immunity in COVID-19: A Neglected but Critical Aspect of SARS-CoV-2 Infection. Frontiers in Immunology 11,.

Rutigliano, J. A., Morris, M. Y., Yue, W., Keating, R., Webby, R. J., Thomas, P. G. and Doherty, P. C. (2010). Protective Memory Responses Are Modulated by Priming Events prior to Challenge. J Virol 84, 1047–1056.

Rutigliano, J. A., Sharma, S., Morris, M. Y., Oguin, T. H., McClaren, J. L., Doherty, P. C. and Thomas, P. G. (2014). Highly Pathological Influenza A Virus Infection Is Associated with Augmented Expression of PD-1 by Functionally Compromised Virus-Specific CD8+ T Cells. Journal of Virology 88, 1636–1651.

Schenkel, J. M., Fraser, K. A., Beura, L. K., Pauken, K. E., Vezys, V. and Masopust, D. (2014). Resident memory CD8 T cells trigger protective innate and adaptive immune responses. Science 346, 98–101.

Schneider, M. A., Brühl, H., Wechselberger, A., Cihak, J., Stangassinger, M., Schlöndorff, D. and Mack, M. (2005). In vitro and in vivo properties of a dimeric bispecific single-chain antibody IgG-fusion protein for depletion of CCR2+ target cells in mice. European Journal of Immunology 35, 987–995.

Shinya, K., Ebina, M., Yamada, S., Ono, M., Kasai, N. and Kawaoka, Y. (2006). Influenza virus receptors in the human airway. Nature 440, 435–436.

Slütter, B., Pewe, L. L., Kaech, S. M. and Harty, J. T. (2013). Lung Airway-Surveilling CXCR3hi Memory CD8+ T Cells Are Critical for Protection against Influenza A Virus. Immunity 39, 939–948.

Slütter, B., Van Braeckel-Budimir, N., Abboud, G., Varga, S. M., Salek-Ardakani, S. and Harty, J. T. (2017). Dynamics of influenza-induced lung-resident memory T cells underlie waning heterosubtypic immunity. Science Immunology 2, eaag2031.

Smillie, C. S., Biton, M., Ordovas-Montanes, J., Sullivan, K. M., Burgin, G., Graham, D. B., Herbst, R. H., Rogel, N., Slyper, M., Waldman, J., et al. (2019). Intra- and Inter-cellular Rewiring of the Human Colon during Ulcerative Colitis. Cell 178, 714–730.e22.

Sposito, B., Broggi, A., Pandolfi, L., Crotta, S., Clementi, N., Ferrarese, R., Sisti, S., Criscuolo, E., Spreafico, R., Long, J. M., et al. (2021). The interferon landscape along the respiratory tract impacts the severity of COVID-19. Cell 184, 4953–4968.e16.

Stary, G., Olive, A., Radovic-Moreno, A. F., Gondek, D., Alvarez, D., Basto, P. A., Perro, M., Vrbanac, V. D., Tager, A. M., Shi, J., et al. (2015). VACCINES. A mucosal vaccine against Chlamydia trachomatis generates two waves of protective memory T cells. Science 348, aaa8205.

Steinbach, K., Vincenti, I., Kreutzfeldt, M., Page, N., Muschaweckh, A., Wagner, I., Drexler, I., Pinschewer, D., Korn, T. and Merkler, D. (2016). Brain-resident memory T cells represent an autonomous cytotoxic barrier to viral infection. Journal of Experimental Medicine 213, 1571–1587.

Sterlin, D., Mathian, A., Miyara, M., Mohr, A., Anna, F., Claër, L., Quentric, P., Fadlallah, J., Devilliers, H., Ghillani, P., et al. (2021). IgA dominates the early neutralizing antibody response to SARS-CoV-2. Science Translational Medicine 13, eabd2223.

Steuerman, Y., Cohen, M., Peshes-Yaloz, N., Valadarsky, L., Cohn, O., David, E., Frishberg, A., Mayo, L., Bacharach, E., Amit, I., et al. (2018). Dissection of Influenza Infection In Vivo by Single-Cell RNA Sequencing. Cell Syst 6, 679–691.e4.

Takaba, H., Morishita, Y., Tomofuji, Y., Danks, L., Nitta, T., Komatsu, N., Kodama, T. and Takayanagi, H. (2015). Fezf2 Orchestrates a Thymic Program of Self-Antigen Expression for Immune Tolerance. Cell 163, 975–987.

Tang, B. M., Shojaei, M., Teoh, S., Meyers, A., Ho, J., Ball, T. B., Keynan, Y., Pisipati, A., Kumar, A., Eisen, D. P., et al. (2019). Neutrophils-related host factors associated with severe disease and fatality in patients with influenza infection. Nat Commun 10, 3422.

Trombetta, J. J., Gennert, D., Lu, D., Satija, R., Shalek, A. K. and Regev, A. (2014). Preparation of Single-Cell RNA-Seq Libraries for Next Generation Sequencing. Current Protocols in Molecular Biology 107,.

Tse, S.-W., Radtke, A. J., Espinosa, D. A., Cockburn, I. A. and Zavala, F. (2014). The Chemokine Receptor CXCR6 Is Required for the Maintenance of Liver Memory CD8+ T Cells Specific for Infectious Pathogens. The Journal of Infectious Diseases 210, 1508– 1516.

Türei, D., Korcsmáros, T. and Saez-Rodriguez, J. (2016). OmniPath: guidelines and gateway for literature-curated signaling pathway resources. Nat Methods 13, 966–967.

Ualiyeva, S., Lemire, E., Wong, C., Perniss, A., Boyd, A. A., Avilés, E. C., Minichetti, D. G., Maxfield, A., Roditi, R., Matsumoto, I., et al. (2024). A nasal cell atlas reveals heterogeneity of tuft cells and their role in directing olfactory stem cell proliferation. Science Immunology 9, eabq4341.

Uddbäck, I., Michalets, S. E., Saha, A., Mattingly, C., Kost, K. N., Williams, M. E., Lawrence, L. A., Hicks, S. L., Lowen, A. C., Ahmed, H., et al. (2024). Prevention of respiratory virus transmission by resident memory CD8+ T cells. Nature 626, 392–400.

Wein, A. N., McMaster, S. R., Takamura, S., Dunbar, P. R., Cartwright, E. K., Hayward, S. L., McManus, D. T., Shimaoka, T., Ueha, S., Tsukui, T., et al. (2019). CXCR6 regulates localization of tissue-resident memory CD8 T cells to the airways. Journal of Experimental Medicine 216, 2748–2762.

Weisberg, S. P., Ural, B. B. and Farber, D. L. (2021). Tissue-specific immunity for a changing world. Cell 184, 1517–1529.

Wellford, S. A., Moseman, A. P., Dao, K., Wright, K. E., Chen, A., Plevin, J. E., Liao, T.-C., Mehta, N. and Moseman, E. A. (2022). Mucosal plasma cells are required to protect the upper airway and brain from infection. Immunity 55, 2118–2134.e6.

Weltzin, R., Traina-Dorge, V., Soike, K., Zhang, J.-Y., Mack, P., Soman, G., Drabik, G. and Monath, T. P. (1996). Intranasal Monoclonal IgA Antibody to Respiratory Syncytial Virus Protects Rhesus Monkeys against Upper and Lower Respiratory Tract Infection. The Journal of Infectious Diseases 174, 256–261.

Wickenhagen, A., Sugrue, E., Lytras, S., Kuchi, S., Noerenberg, M., Turnbull, M. L., Loney, C., Herder, V., Allan, J., Jarmson, I., et al. (2021). A prenylated dsRNA sensor protects against severe COVID-19. Science 374, eabj3624.

Wiley, J. A., Hogan, R. J., Woodland, D. L. and Harmsen, A. G. (2001). Antigen-Specific CD8+ T Cells Persist in the Upper Respiratory Tract Following Influenza Virus Infection. J Immunol 167, 3293–3299.

Woof, J. M. and Mestecky, J. (2005). Mucosal immunoglobulins. Immunological Reviews 206, 64–82.

Xie, X., Shi, Q., Wu, P., Zhang, X., Kambara, H., Su, J., Yu, H., Park, S.-Y., Guo, R., Ren, Q., et al. (2020). Single-cell transcriptome profiling reveals neutrophil heterogeneity in homeostasis and infection. Nat Immunol 21, 1119–1133.

Xu, C., Lopez, R., Mehlman, E., Regier, J., Jordan, M. I. and Yosef, N. (2021). Probabilistic harmonization and annotation of single-cell transcriptomics data with deep generative models. Molecular Systems Biology 17, e9620.

Yao, Y., Jeyanathan, M., Haddadi, S., Barra, N. G., Vaseghi-Shanjani, M., Damjanovic, D., Lai, R., Afkhami, S., Chen, Y., Dvorkin-Gheva, A., et al. (2018). Induction of Autonomous Memory Alveolar Macrophages Requires T Cell Help and Is Critical to Trained Immunity. Cell 175, 1634–1650.e17.

Zhang, Q., Bastard, P., Liu, Z., Pen, J. L., Moncada-Velez, M., Chen, J., Ogishi, M., Sabli, I. K. D., Hodeib, S., Korol, C., et al. (2020). Inborn errors of type I IFN immunity in patients with life-threatening COVID-19. Science 370,.

Zhang, H., Alford, T., Liu, S., Zhou, D. and Wang, J. (2022a). Influenza virus causes lung immunopathology through down-regulating PPARγ activity in macrophages. Frontiers in Immunology 13,.

Zhang, K., Erkan, E. P., Jamalzadeh, S., Dai, J., Andersson, N., Kaipio, K., Lamminen, T., Mansuri, N., Huhtinen, K., Carpén, O., et al. (2022b). Longitudinal single-cell RNA-seq analysis reveals stress-promoted chemoresistance in metastatic ovarian cancer. Science Advances 8, eabm1831.

Zheng, H. B., Doran, B. A., Kimler, K., Yu, A., Tkachev, V., Niederlova, V., Cribbin, K., Fleming, R., Bratrude, B., Betz, K., et al. (2021). A Treatment-Naïve Cellular Atlas of Pediatric Crohn’s Disease Predicts Disease Severity and Therapeutic Response. 2021.09.17.21263540.

Zheng, H. B., Doran, B. A., Kimler, K., Yu, A., Tkachev, V., Niederlova, V., Cribbin, K., Fleming, R., Bratrude, B., Betz, K., et al. (2023). Concerted changes in the pediatric single-cell intestinal ecosystem before and after anti-TNF blockade. eLife 12,.

Ziegler, C. G. K., Allon, S. J., Nyquist, S. K., Mbano, I. M., Miao, V. N., Tzouanas, C. N., Cao, Y., Yousif, A. S., Bals, J., Hauser, B. M., et al. (2020). SARS-CoV-2 Receptor ACE2 Is an Interferon-Stimulated Gene in Human Airway Epithelial Cells and Is Detected in Specific Cell Subsets across Tissues. Cell 181, 1016–1035.e19.

Ziegler, C. G. K., Miao, V. N., Owings, A. H., Navia, A. W., Tang, Y., Bromley, J. D., Lotfy, P., Sloan, M., Laird, H., Williams, H. B., et al. (2021). Impaired local intrinsic immunity to SARS-CoV-2 infection in severe COVID-19. Cell 184, 4713–4733.e22.

